# Automated high-throughput fabrication of patient-specific vessel-on-chips enables a generative AI digital twin—Cascade Learner of Thrombosis (CLoT) for personalized thrombosis prediction

**DOI:** 10.64898/2026.03.03.709446

**Authors:** Zihao Wang, Yunduo Charles Zhao, Haimei Zhao, Arian Nasser, Nicole Alexis Yap, Yanyan Liu, Allan Sun, Wei Chen, Ken S Butcher, Timothy Ang, Lining Arnold Ju

## Abstract

We developed an integrated platform combining high-throughput automated biofabrication, systematic patient-derived tissue experiments, and specialized artificial intelligence to enable patient-specific computational “digital twins” for thrombosis prediction. Our automated manufacturing platform fabricates 80 fully assembled, patient-specific vessel-on-chips within 10 hours from clinical imaging—a ∼100-fold improvement over manual methods—achieving sub-micron precision through novel two-stage pneumatic motion control and integrated optical feedback. Using these chips, we systematically captured thrombosis across 491 high-fidelity videos spanning 6 patient-derived vascular geometries, 5 distinct anatomical injury sites, and 14 anticoagulant/antiplatelet interventions, establishing a “physical twin” experimental corpus. We trained CLoT (Cascade Learner of Thrombosis), a conditional video diffusion model efficiently adapted via lightweight Low-Rank Adaptation (LoRA) to generate realistic thrombosis videos conditioned on patient-specific geometry, injury location, and drug treatment. Rigorous benchmarking against state-of-the-art commercial models (Sora, Wan, Kling, Seedance, Hailuo, Hunyuan) reveals CLoT achieves 7.38-fold superior temporal biological consistency and 5.3-fold higher spatial morphological fidelity. Prospective validation on unseen patients demonstrates >90% temporal accuracy. This integrated paradigm—combining automated fabrication with domain-specialized generative AI—establishes proof-of-concept for personalized medicine enabled by digital twins trained on human-derived vascular anatomy, enabling pre-treatment antithrombotic evaluation while providing a replicable template for translating tissue engineering into clinical practice.

## 1. Introduction

Over the past decade, microfluidic organ-on-chip systems have emerged as transformative platforms for recapitulating human tissue complexity and drug responses with superior fidelity to animal models (*1–5*). Pioneered by international colleagues (*1, 6, 7*) with the lung-on-a-chip, and expanded to vascular (*8–15*), cardiac (*16–18*), neural (*19*), liver (*20*), and kidney (*21, 22*) systems, these devices exploit microfluidic principles to recreate critical features of in vivo biology: physiologic shear stress, cell–cell communication, 3D tissue architecture, and temporal dynamics of pathological processes. Among these systems, patient-specific vessel-on-chip platforms are particularly relevant for thrombotic diseases, leading cause of mortality globally, with thrombotic events—including stroke, myocardial infarction, and stent thrombosis—accounting for over 17 million deaths annually (*23*), as they enable the recapitulation of thrombogenesis—a multifactorial process governed by the classical Virchow’s triad (endothelial injury, blood stasis/flow disturbance, blood hypercoagulability) (*13*). Recent vessel-on-chip studies have demonstrated human-relevant thrombosis kinetics (*14, 24–29*), platelet-count-independent microthrombi formation (*30*), and predictive drug responses (*31*) that diverge from animal models.

Predicting and preventing thrombosis in patient-specific vascular geometries requires understanding how vessel morphology, hemodynamic shear stress, and pharmacological interventions interact to modulate platelet aggregation and thrombus stability (*32–35*). However, a critical bottleneck limits this clinical translation: existing vessel-on-chip fabrication relies on photolithography, soft-lithography, and other labor-intensive processes (*36–39*), which limits its throughput, agility and adaptability. Scaling to personalized medicine—producing patient-specific chips for each individual—demands orders-of-magnitude improvements of throughput and reproducibility (*39–41*).

Parallel to tissue-engineering advances, digital twin technology has revolutionized engineering and manufacturing by creating high-fidelity computational mirrors of physical systems (*42–44*). Originally developed for aerospace and automotive applications, digital twins are now being adapted for personalized medicine (*45–47*). The concept is powerful: if comprehensive experimental data from a patient’s “physical twin” (e.g., a patient-specific organ-on-a-chip) can be acquired, then AI models trained on such data can predict outcomes for new conditions, optimizing treatment without exhausting biological resources.

Recent advances in generative AI, particularly Diffusion Transformers (*48–50*), have demonstrated remarkable capability to synthesize realistic, temporally coherent video sequences. OpenAI’s Sora model (*51*), trained on vast natural-video corpora, generates photorealistic videos from text prompts with unprecedented coherence (*52*). However, Sora and comparable models (Alibaba Wan (*53*), Kaishou Kling (*54*), Bytedance Seedance (*55*), MiniMax Hailuo (*56*), Tencent Hunyuan (*57*)) are optimized for natural scenes; their generalization to biological fluorescence microscopy, hemodynamic phenomena, and patient-specific geometry remains untested. Finetuning such pretrained backbones via parameter-efficient techniques (e.g., Low-Rank Adaptation; LoRA) offers a path to adapt foundation models to specialized domains (*58*) without retraining from scratch, democratizing AI deployment for resource-constrained laboratories.

We address these barriers via an integrated platform: (1) a fully automated vessel-on-chips manufacturing system that translates clinical vascular imaging into patient-specific microfluidic devices at high throughput within hours (Fig. 1A), (2) high-fidelity blood-perfusion experiments on patient-specific vessel-on-chips establishing a “physical twin” dataset of thrombotic dynamics across diverse clinical phenotypes and pharmacological interventions (Fig. 1B), and (3) a specialized generative AI model (CLoT; Cascade Learner of Thrombosis) that learns from physical-twin videos to predict thrombotic outcomes under untested conditions, effectively creating a computational “digital twin” (Fig. 1C).

**Figure 1.**
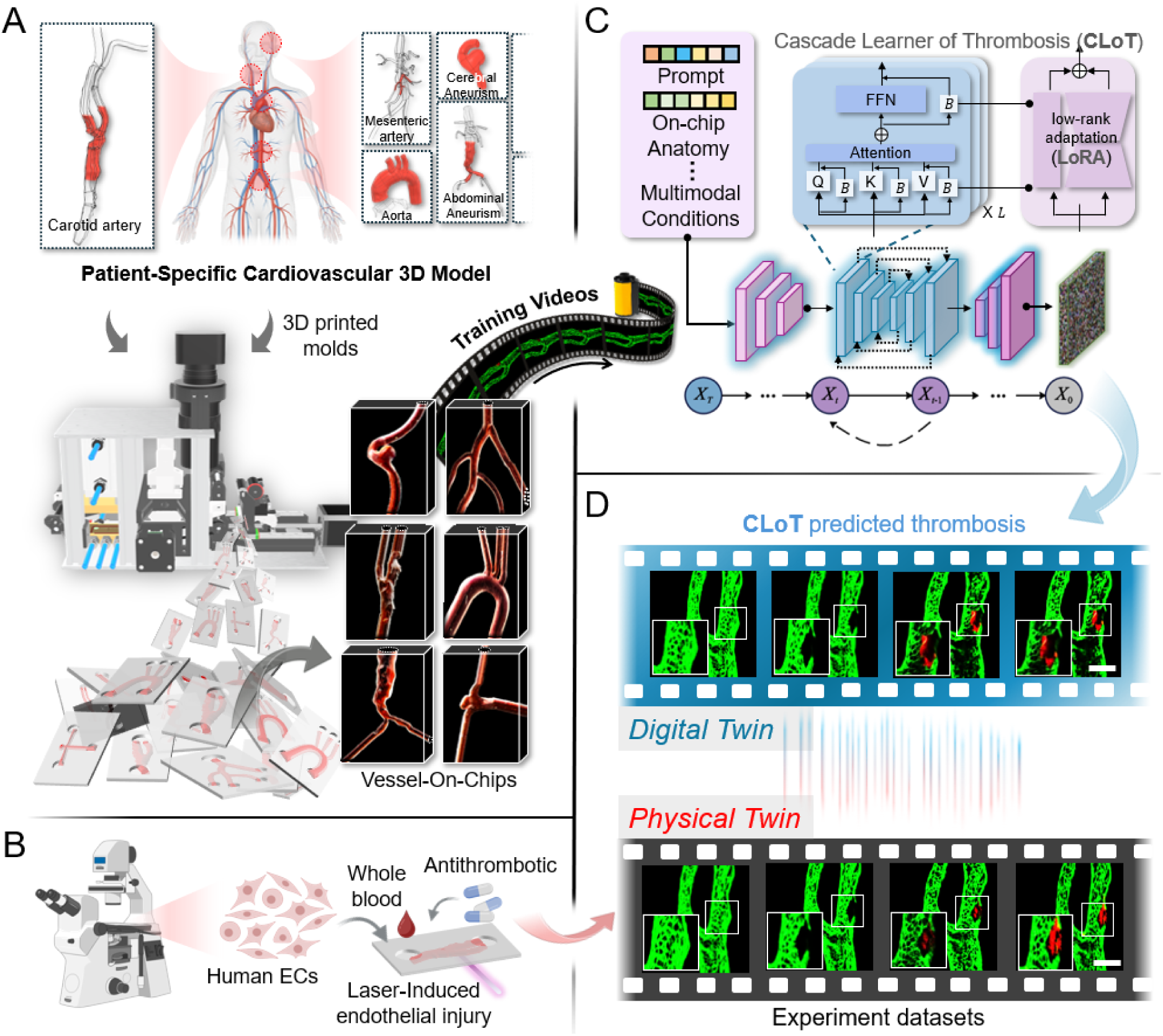
Integration of automated patient-specific vessel-on-a-chip manufacturing with generative AI digital twin thrombosis modeling. (**A**) Overview of the patient-specific thrombosis modeling workflow. Three-dimensional vascular geometries were reconstructed from clinical imaging (e.g., carotid, mesenteric, aortic, and cerebral vasculatures) and micro-3D printed as positive molds. These molds were processed by the automated manufacturing system to generate SEBS-based vessel-on-chip devices at high-throughput corresponding to each patient’s vasculature. The resulting chips were endothelialized and perfused with whole blood under physiological flow, producing training videos for the AI model. (**B**) Experimental setup for physical twin thrombus formation. Endothelialized chips were perfused with whole blood containing pharmacological agents, and thrombus initiation was triggered by targeted laser ablation under real-time confocal microscopy. (**C**) Schematic of the multimodal AI model, Cascade Learner of Thrombosis (CLoT). The model incorporates text descriptions, vascular geometry, and vessel-on-chip experimental conditions to learn thrombosis dynamics through a diffusion transformer framework combining attention and feed-forward layers with a lightweight cascade-learning module. (**D**) Comparison between AI-generated and experimentally observed clot formation. The AI-generated videos (top) reproduce clot morphology and growth trajectories consistent with physical experiments (bottom), establishing a one-to-one correspondence between the digital twin and physical twin representations of patient-specific thrombotic events.

Our automated fabrication platform involves five coordinated modules—pneumatic, film-feeding, thermoforming, punching, and alignment—coordinated by a centralized controller for real-time control. The platform achieves sub-micron level precision during optical alignment of microfluidic channel halves, enabling production of 80 fully assembled, patient-specific vessel chips within 10 hours of clinical imaging acquisition. This represents a ∼100 × improvement over conventional manual fabrication pipeline.

To train the digital twin, we systematized blood perfusion assays on patient-specific chips, performing controlled endothelial injury at anatomically distinct sites and evaluating responses to 14 anticoagulants/antiplatelet agents. This experimental corpus (n = 491 videos (90% of 546 qualified videos in total); 6 patient geometries; 5 injury sites; 14 drugs) serves as training datasets for CLoT, a conditional video diffusion model adapted from the Wan backbone via LoRA finetuning. CLoT learns to synthesize realistic thrombosis videos conditioned on vessel geometry, injury location, and drug treatment, effectively capturing patient-specific and site-specific hemodynamic coupling to platelet biology (Fig. 1D). Detailed video dataset collection and construction procedure can be found in Tables 1 and S1.

## 2. Results

### 2.1. Automated fabrication platform enables rapid, patient-specific vessel-on-a-chip production

The automated fabrication platform functions as an integrated pipeline that transforms a styrene–ethylene–butylene–styrene (SEBS) thermoplastic elastomer film and a glass-based master mold—3D printed via our previously established microprinting method (*28*)—into fully assembled, patient-specific vessel-on-a-chip devices (Fig. 2A). The platform performs three sequential steps: thermoforming, punching, and alignment. During thermoforming, the SEBS film is hot-embossed against the 3D-printed glass mold to replicate the vascular geometry. The punching module then perforates inlet and outlet ports to establish perfusion interfaces. Finally, the molded film containing two mirror-placed half-channel geometries is folded and precisely aligned to form a sealed, full-lumen vessel-on-chip (Fig. 2B). The entire production process is fully automated and only requires minimal post process after the chip production (movie S1, S2). The completed chips are subsequently biofunctionalized by endothelial cells (ECs) seeding and static culture to generate a confluent EC monolayer (Fig. 2C). Under optimized operating conditions, the platform can produce 80 fully assembled patient-specific chips within 10 hours from clinical image acquisition—achieving more than a 100-fold increase in throughput compared with conventional manual fabrication (Fig. 2D, *upper*). Scanning electron microscopy confirmed high-fidelity replication of vascular features (Fig. 2D, *lower*), validating the precise geometry transfer of our platform.

**Figure 2.**
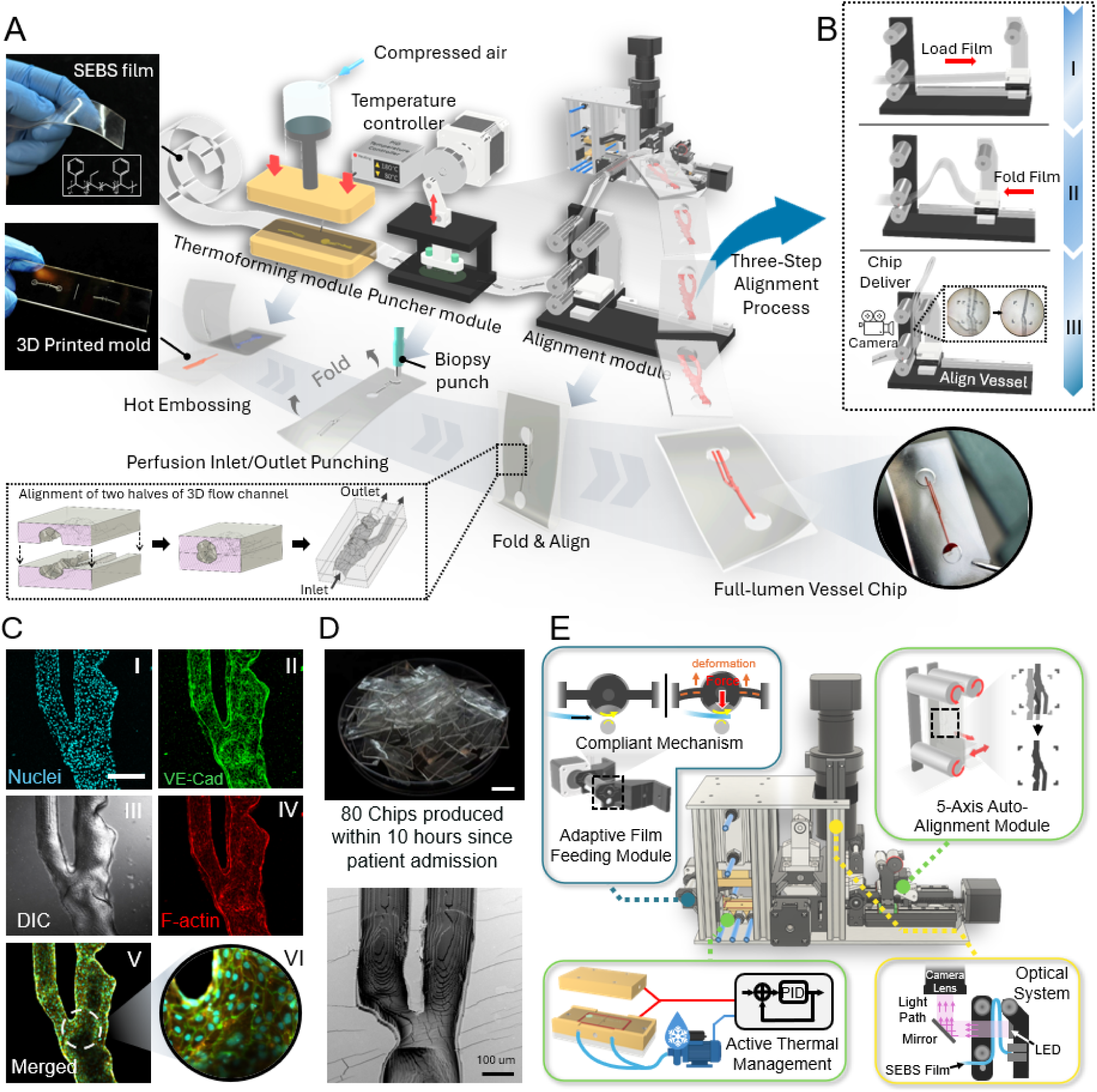
Automated fabrication platform for high-throughput patient-specific vessel-on-a-chip production. (**A**) Overview of the integrated vessel-on-a-chip manufacturing pipeline. Patient-specific vascular molds are first fabricated via digital light processing (DLP) micro-3D printing, followed by automated SEBS chip fabrication through sequential thermoforming, punching, and alignment modules. Each module operates under semi-close loop control to enable rapid, continuous production of perfusable 3D vessel-on-chips. (**B**) Sequential schematic of the three-step alignment process. I, SEBS film is grasped by the film grabber and loaded onto the alignment stage. II, The folding stage moves inward, folding the film until its two halves overlap; the peak of the folded film is grabbed and tensioned by the stationary and motive rollers to stabilize the structure. III, The horizontal stage, tension rollers, and feeding roller operate in coordination under camera feedback to achieve precise alignment between the upper and lower channel halves. (**C**) Representative imaging of a fabricated carotid artery-chip demonstrating geometric fidelity and confluent endothelialization. I, nuclei (*blue*); II, VE-cadherin (*green*); III, F-actin (*red*); IV, differential interference contrast (DIC); V, merged fluorescence channels; VI, magnified view of the merged image showing continuous endothelial junctions along the bifurcation. (**D**) Top: A total of 80 fully assembled patient-specific vessel-on-chips were fabricated within 10 hours from clinical image acquisition. Bottom: scanning electron micrograph (SEM) of the inner lumen of a thermoformed carotid artery-chip, demonstrating high structural fidelity and accurate replication of fine mold features. Scale bars: top, 10 mm; bottom, 100 µm. (**E**) Schematic of key subsystems in the automated platform, including the adaptive film-handling mechanism, PID-controlled thermal module, five-axis auto-alignment module, and optical path for image-guided alignment and automated chip delivery. The film-handling module comprises an active stepper-driven roller and a passive roller mounted on a compliant 3D-printed mechanism. As the SEBS film enters, the compliant support deforms upward to apply a uniform counter-force, maintaining stable traction and preventing slippage. The optical subsystem uses a single LED mounted on the folding arm to illuminate the film, with reflected light directed to a camera through a 45° mirror, allowing simultaneous visualization of both film halves during alignment. Scale bar: 300 µm.

To smoothly and automatically perform these three steps, the platform consists of five modules that operate in continuous coordination through a central control board, translating clinical imaging data into functional vessel-on-a-chip devices that recapitulate patient-specific vascular geometries (Fig. 2E, fig. S1). The SEBS film is first introduced via an adaptive film-feeding module comprising a motorized active roller and a passive roller mounted on a compliant mechanism. Upon film insertion, the passive roller deforms upward to apply a uniform downward pressure, ensuring firm contact with the active roller and prevent slippage during feeding (Fig. 2E, *upper left*).

The film-feeding module subsequently advances the SEBS film into the thermoforming module for hot embossing onto the glass-mounted 3D-printed mold. This module integrates dual brass heating blocks equipped with cartridge heaters and thermocouples for precise temperature regulation, a thermal bridge and a thermal isolation pad for heat insulation, and a pneumatic actuator to deliver controlled compression (fig. S2A). During the embossing process, the SEBS film is placed over the mold, which is then sandwiched between the two heat blocks and compressed by the pneumatic actuator to replicate the mold geometry. The final chip thickness—critical for ensuring that the channel depth remains within the working distance of standard microscope objectives—is defined by the cavity depth of the bottom block in combination with the mold dimensions (fig. S2B). A vacuum-lock interface embedded within the bottom heat block secures the glass mold during embossing while allowing rapid mold exchange and maintaining precise positioning (fig. S2C).

To ensure reproducible molding, the top and bottom blocks are independently temperature-regulated through a Proportional-Integral-Derivative (PID) controller coupled with active water cooling (Fig. 2E, *lower left* and fig. S2D). Thermal characterization confirmed that this combined PID control and heat isolation architecture maintains temperature stability within ±2 °C under idle conditions and within ±11 °C during continuous high-throughput operation (fig. S3, S4, A and B). The integrated active water-cooling circuit embedded within the bottom-block stabilizes its temperature during extended production, enabling more than 12 consecutive fabrication cycles without significant temperature drift (fig. S4, *B* and C). The use of 3D-printed and ceramic polymer components for thermal isolation reduced parasitic heat transfer to pneumatic cylinder by 51% compared with aluminum counterparts while preserving overall temperature uniformity (fig. S4D, S5, A to D, S6). These design features collectively enable prolonged, thermally stable, and reproducible molding performance.

**Figure 3.**
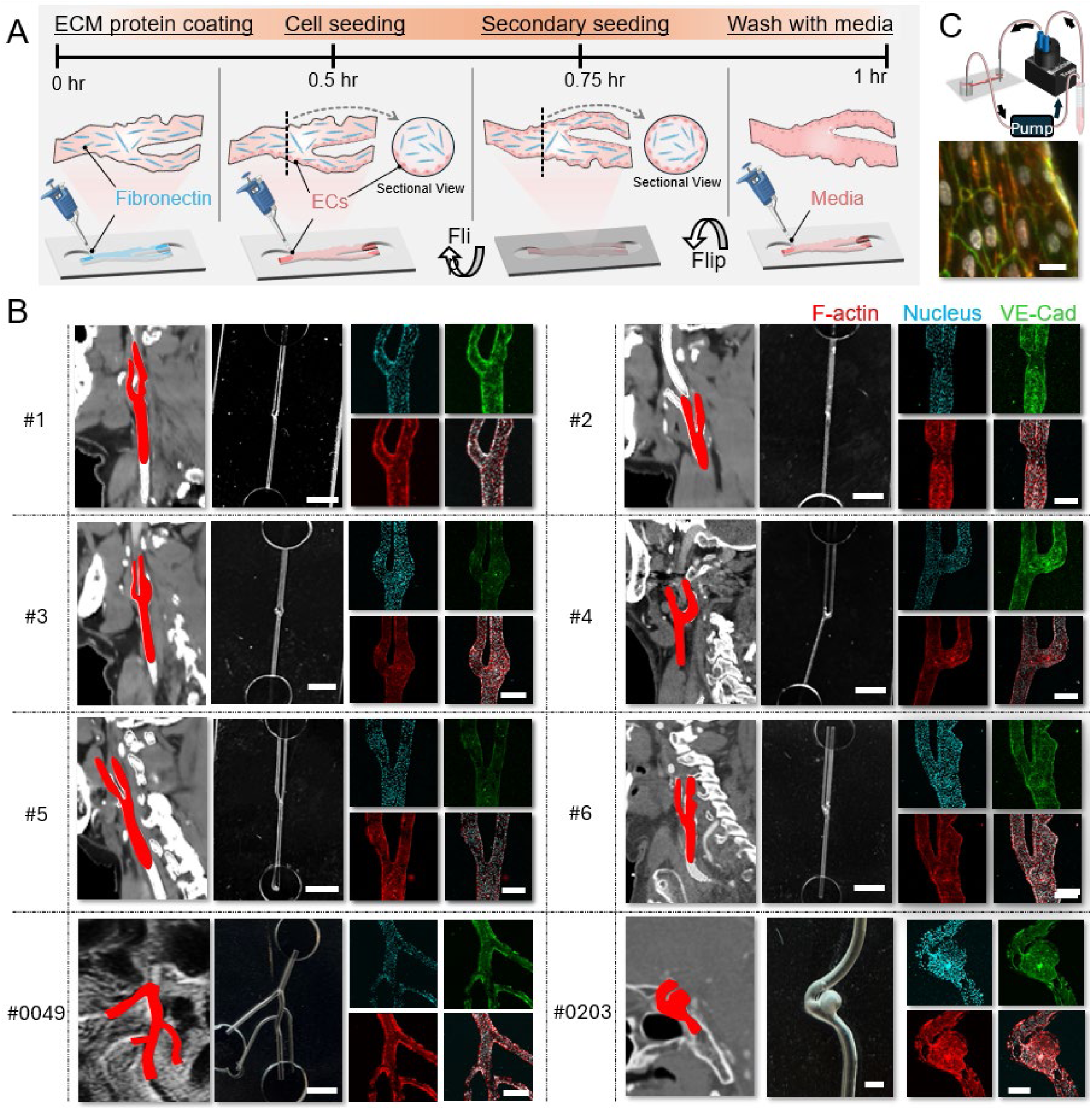
Endothelialization of the patient-specific vessel-on-a-chip models. (**A**) Stepwise schematic of the endothelial seeding and culture process. The microchannel surfaces were first coated with fibronectin (0.5 hr). Human umbilical vein endothelial cells (HUVECs) were then introduced for the first seeding (0.25 h) to establish attachment on one side of the channel. The chip was subsequently flipped (0.25 h) to achieve bilayer endothelial coverage. Finally, the device was perfused with culture medium and incubated overnight to form a confluent endothelium. (**B**) Representative examples demonstrating the system’s capability to reproduce anatomically accurate, patient-specific vascular geometries (denoted as #1–#6 for carotid artery patients and #0049, #0203 as their legacy name in Stanford VASCULAR MODEL REPOSITORY database). For each patient, the first column shows clinical CT angiography (CTA) scans of the targeted vessel region with 3D-reconstructed vascular models highlighted in red. The second column presents photographs of the corresponding vessel-on-chips. The third and fourth columns display fluorescence immunostaining of the vessel lumen, showing endothelial distribution and morphology. Blue, nuclei (Hoechst); green, VE-cadherin (endothelial junction marker); red, F-actin (cytoskeletal structure). Merged channels confirm confluent endothelial coverage and alignment consistent with patient-specific vessel geometries. (**C**) Illustration of dynamic cell culture setup and the fluorescence immunostaining image of the cell cultured under 2 days of dynamic culture. Color codes are same as those in (B). Scale bars: Panel B, third column (except bottom right), 2 mm; bottom right, 300 µm; fourth and fifth columns, 300 µm; Panel C, 50 µm.

**Figure 4.**
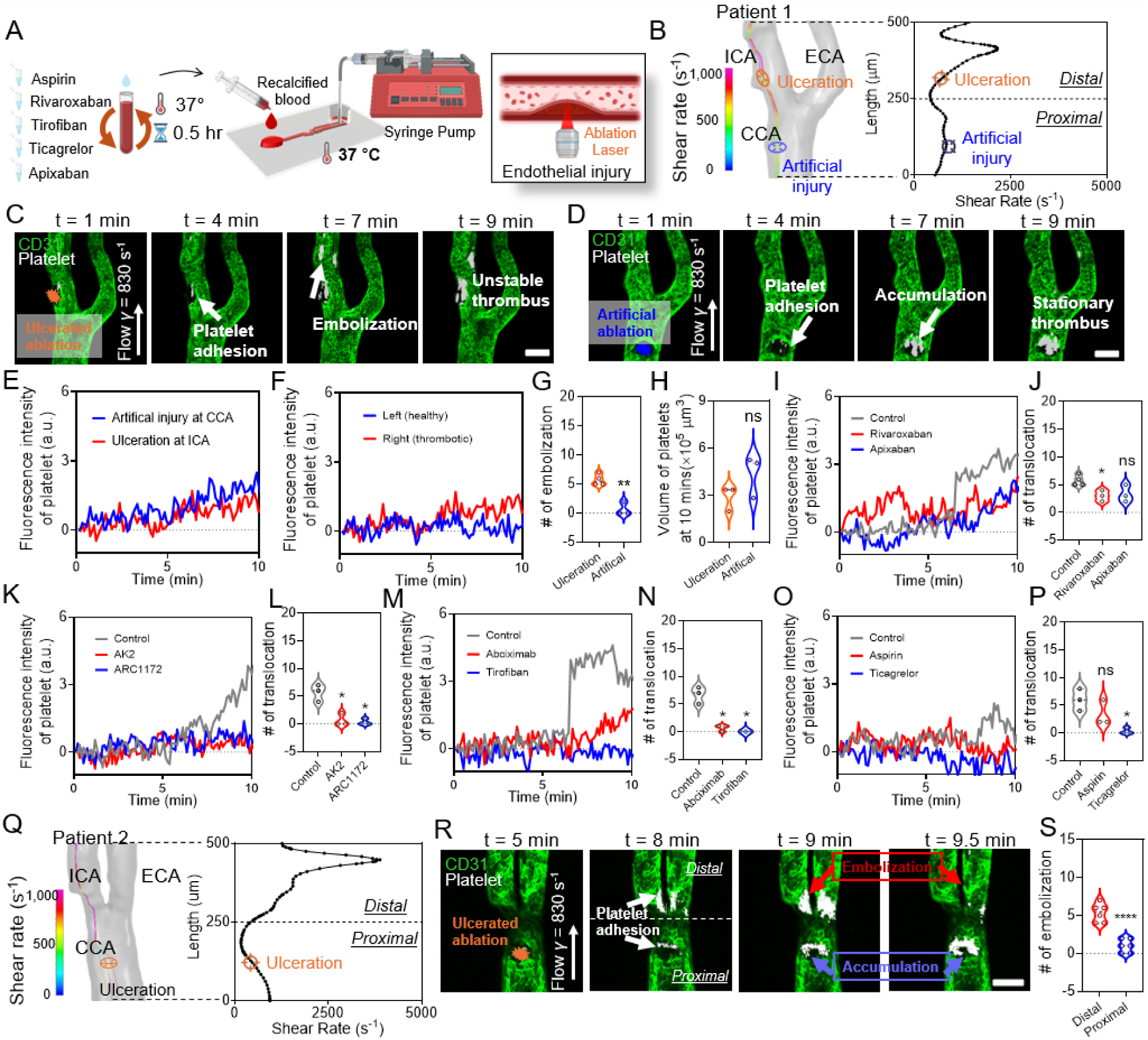
Patient-specific carotid artery-on-chip platform for assessing site-dependent thrombosis and drug responses. (**A**) Schematic of the whole-blood perfusion and intervention workflow. Recalcified human blood, pre-incubated for 30 min at 37 °C with antithrombotic agents, was perfused through the vessel-on-chips. Localized endothelial injury was induced in real time by laser ablation. (**B**) Hemodynamic reconstruction for Patient 1 showing shear-rate distribution along the carotid bifurcation and the locations of the natural ulceration at the internal carotid artery (ICA, *orange*) and the artificial injury at the common carotid artery (CCA, *blue*). Line plot (*right*) shows the corresponding shear profile from proximal to distal segments. (**C–D**) Representative confocal time-lapse images (CD31, *green*; platelets, *white*) showing thrombus formation at the ulcerated ICA (C) and at the artificial CCA site (D) under 830 s⁻¹ bulk shear. Ulcerated sites exhibited repeated platelet adhesion and embolization, whereas artificial sites formed stationary thrombi. Scale bars: 200 µm. (**E–F**) Time-resolved platelet fluorescence intensity at ulcerated versus artificial sites (E) and at the left (healthy) versus right (thrombotic) carotid branches (F), illustrating higher instability at the pathological regions. (**G**) Violin plot quantifying the number of embolization events over 10 min shows significantly more embolization at ulcerated than artificial sites. (**H**) Platelet volume at 10 min was comparable between the two sites, indicating that flow localization governs stability rather than total mass. (**I–J**) Effects of oral anticoagulants (rivaroxaban, apixaban): time-lapse platelet accumulation (I) and embolization counts (J) show that rivaroxaban exhibited greater inhibition of platelet dynamics relative to the control, while apixaban produced a lesser and statistically non-significant effect (ns). (**K–L**) Blockade of VWF–GPIb using AK2 and ARC1172 markedly reduced platelet accumulation (K) and significantly lowered translocation events (L). (**M–N**) Integrin αIIbβ3 inhibition (abciximab, tirofiban) almost abolished thrombus growth (M) and reduced embolization to near baseline (N), confirming fibrinogen/αIIbβ3-dependent consolidation under these shear conditions. (**O–P**) Antiplatelet agents targeting TXA₂ (aspirin) or P2Y₁₂ (ticagrelor) showed differential effects: ticagrelor more effectively blunted platelet accumulation (O) and reduced embolization (P), whereas aspirin was modest. (**Q**) Hemodynamic mapping for Patient 2 identifying an ICA ulceration located at a distal, higher-shear region. (**R**) Time-lapse imaging in Patient 2 shows that distal ulceration favors embolizing thrombi, while proximal segments with lower shear support stable accumulation. Scale bar: 200 µm. (**S**) Violin plot comparing distal versus proximal sites in Patient 2 confirms significantly more embolization at distal, high-shear ulcerations. Statistical significance is indicated by * = p < 0.05, ** = p < 0.01, *** = p < 0.0001 assessed by unpaired, two-tailed Student’s t-test.

**Figure 5.**
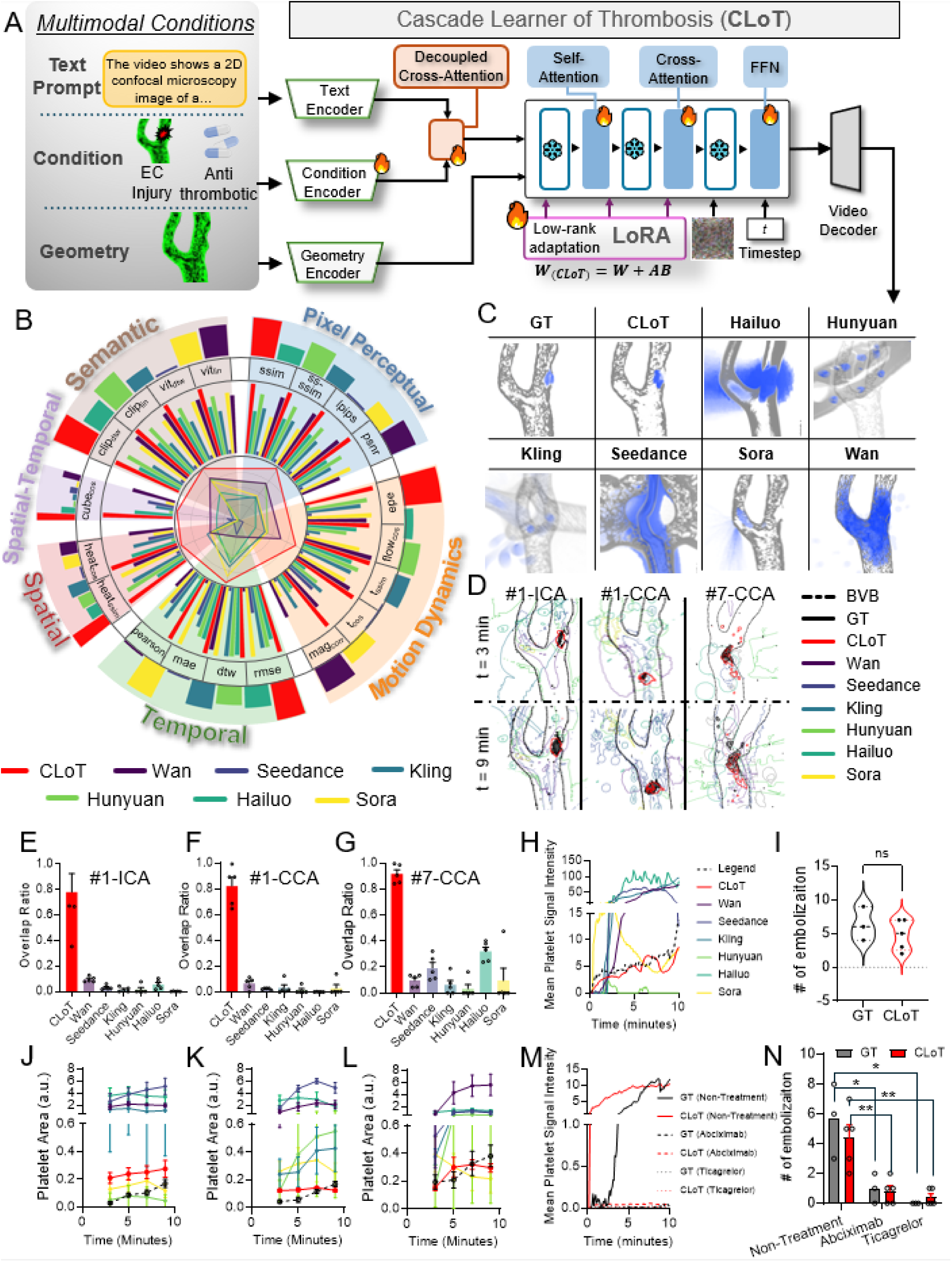
Generative AI model for thrombosis video synthesis and performance evaluation. (**A**) Model architecture. The Cascade Learner of Thrombosis (CLoT) integrates blood vessel geometry, injury site, pharmacological treatment, and natural-language experimental descriptors as multimodal conditions for video generation. Geometry, condition, and text encoders feed into a decoupled cross-attention transformer whose weights are adapted from a pretrained foundation video-generation model through lightweight cascade learning (*W*_CLoT=*W*+*AB*). The system generates microscopy-style videos depicting thrombosis dynamics conditioned on patient-specific and pharmacological contexts. (**B**) Circular, category-structured evaluation of video-generation quality with embedded radar. We compare CLoT (ours), Sora, Wan, Kling, Seedance, Hailuo, and Hunyuana across six metric families: Pixel Perceptual (PSNR, SSIM, MS-SSIM, LPIPS), Motion Dynamics (EPE, flow cosine similarity, flow-magnitude correlation, temporal SSIM, temporal RGB cosine), Temporal signal (Pearson’s r, DTW, MAE, RMSE), Spatial pattern (heatmap SSIM, heatmap cosine), Spatiotemporal Bio-pattern (occupancy-cube cosine), Semantic Consistency (ViT/CLIP linear similarity and DTW-aligned similarity), and temporospatial. Each spoke corresponds to a metric (short labels shown tangentially). For every metric, methods are rendered as adjacent inner radial bars; greater radial height indicates better relative performance after per-metric min–max normalization across methods. Metrics where “lower is better” (EPE, LPIPS, DTW, MAE, RMSE) are inverted before scaling; a *γ*-correction (*γ* = 0.6) enhances mid-range contrast. Category sectors are lightly tinted for readability. An inner radar summarizes the mean score of each method within every category (higher radius = better). The outer ring encodes the category mean rescaled to 10–100 (same color as inner bars), enabling quick, family-level comparison. Overall, CLoT occupies the largest radar area and shows consistently taller outer-ring segments, indicating superior performance across most categories. (**C**) Temporal-trajectory (“long-exposure”) visualization of platelet activation for Patient #1. Representative videos from ground truth (GT) and seven AI models (CLoT, Hailuo, Hunyuan, Kling, Seedance, Sora, Wan) were processed to generate cumulative persistence maps. Vascular structure (green channel) is rendered in gray; platelet signals (red channel) are rendered in blue. Darker regions indicate prolonged signal presence across the 10-minute observation period; lighter regions indicate transient activation. Each image represents the composite spatiotemporal propagation pattern of a single video replicate. (**D**) Spatial distribution of platelet activation at injury sites visualized as contour overlays at t = 3 min and t = 9 min. Left to right: Patient #1 ICA injury site, Patient #1 CCA injury site, Patient #7 CCA injury site. Ground-truth platelet regions are shown as filled black masks (70% opacity); CLoT as red contours (2-pt thickness); other AI models as colored contours (1-pt thickness, viridis colormap). Blood vessel boundaries (BVB, black dash-dot line) extracted from the first frame of reference. CLoT videos provide anatomical context. Representative videos selected from each model’s five replicates. (**E–G**) Spatial overlap ratios between AI-generated and ground-truth platelet distributions at 9 mins of equivalent experiment time. Individual data points represent single video replicates (n = 5 per AI model). Overlap ratio calculated as |At ∩ Gt| / max(|At|, |Gt|), where At and Gt denote binary platelet masks from AI and ground-truth frames at time t. Bars represent mean ± S.E.M. (**H**) Temporal evolution of average platelet signal intensity for Patient #7. Each colored trace represents a single representative video from ground truth (dashed black line) or AI models (CLoT: red, Wan: purple, Seedance: blue, Kling: cyan, Hunyuan: green, Hailuo: teal, Sora: yellow). Intensity values are baseline-subtracted (first frame = 0) and calculated within the 140×140-pixel Intensity ROI centered on the injury site. Inset: magnified view of 0–15 intensity range. Time normalized to 10-minute real-time experimental duration. (**I**) Embolization (platelet translocation) counts comparing GT and CLoT, indicating no significant difference in predicted embolic instability under matched conditions. (**J–L**) Platelet area dynamics over time for ground truth and seven AI models. Platelet area (count of valid red pixels in 360×360-pixel Analysis ROI) plotted as mean ± S.E.M. across all replicates (ground truth n = 3, AI models n = 5 each). Time normalized to 10-minute duration. (J) Patient #1, ICA injury; (K) Patient #1, CCA injury; (L) Patient #7, CCA injury. (**M–N**) Prospective digital-twin prediction of drug responses. Mean platelet-signal trajectories (M) and embolization counts (N) for non-treatment, abciximab (αIIbβ3 blockade), and ticagrelor (P2Y_12_ antagonism) show that CLoT reproduces GT drug-specific suppression of thrombus growth and embolization. Statistical significance is indicated by * = p < 0.05, ** = p < 0.01, *** = p < 0.001, **** = p < 0.0001; ns, not significant.

The pneumatic cylinder drives the top heat block toward the bottom block to press the SEBS film onto the master mold under tightly regulated force. A dual-stage pneumatic control circuit governs three sequential phases—preheat, emboss, and release—to ensure precise softening and forming of SEBS without mechanical shock (fig. S7). During preheating (step 1), low-pressure air (∼1 bar) gently lowers the top block, softening the SEBS above its glass-transition temperature (T_g_ ≈ 100 °C). In the hot embossing phase (step 2), high-pressure actuation (∼5 bar) produces ∼950 N of compressive force, fully filling complex vascular cavities. The release phase (step 3) reverses airflow, venting high-pressure air and reintroducing low-pressure air to lift the top block smoothly and prepare for the next cycle.

Failure-mode analysis confirmed the robustness of this two-stage pneumatic–thermal process and thermal control design (fig. S8 and S9). Insufficient heating yielded incomplete cavity filling and rounded channel edges (fig. S8, B and C, fig. S9, A to C), whereas excessive heating caused SEBS bubbling and mold outgassing (fig. S9, D to F). Adequate bottom-block temperature was critical for successful demolding, minimizing adhesion between the molded film and master mold (fig. S9 G and H). Importantly, the soft-start/soft-stop motion prevented catastrophic glass fracture from abrupt pneumatic shocks (fig. S9I).

Following thermoforming, the film enters the puncher module, which converts servo-driven rotary motion into precise linear displacement via a 3D-printed crankshaft linkage (fig. S10, A and B). This mechanism drives dual biopsy punches to form perfusion inlet and outlet with tunable force balance and cutting depth (fig. S10C). The servo-controlled cycle allows programmable cutting periods to accommodate varying film thicknesses and material stiffness (fig. S10D). A depth-tuning design ensures partial material retention at the punch site, preventing debris accumulation during continuous operation (fig. S11).

The hot-embossed SEBS film, now containing the molded channel geometries and perfusion inlets and outlets, is then transferred to the final stage—the alignment module—for assembly into a full-lumen, perfusable vessel-on-a-chip device. The alignment module integrates five motorized axes that provide complete three-dimensional spatial control of film positioning during chip alignment (fig. S12). The alignment workflow comprises three sequential phases: pre-alignment, alignment, and post-alignment. During pre-alignment, the film is first advanced and folded. Because the SEBS film is soft and flexible, dual stepper-driven tension rollers apply controlled stretching forces to tension and stabilize it, completing the pre-alignment phase (fig. S13A). In the subsequent alignment phase, a real-time optical feedback loop detects fiducial markers to achieve submicron-scale registration between the two vessel halves (fig. S13B; figs. S14 and S15). After alignment, the tension rollers act as compression rollers, gently pressing the layers together during delivery to yield a seamless, bubble-free, and fully enclosed vessel-on-a-chip device with embedded patient-specific luminal geometries suitable for subsequent endothelialization (fig. S13C).

### 2.2 Patient-specific vascular geometry reproduction across diverse clinical phenotypes

We next demonstrated the biofunctionalization of patient-specific vessel-on-chips with endothelial cells (Fig. 3A) and validated the system’s ability to reproduce anatomically complex vascular geometries derived from both clinical and open-source datasets (Fig. 3B). To further assess device performance under physiological flow, we conducted dynamic cell culture experiments using the fabricated chips, demonstrating that the system supports the formation of polarized endothelial monolayers within the perfused channels (Fig. 3C). Starting from clinical CT angiography (CTA) images, 3D vascular models were reconstructed, converted into 3D-printed glass molds, and used as templates for automated thermoforming. The resulting chips faithfully recapitulated the distinct bifurcation and curvature features of the original vascular anatomies, as confirmed by microscopic inspection (fig. S17).

Vessel-on-chips were fabricated using carotid artery geometries from different patients exhibiting diverse bifurcation morphologies with each geometry numerically labeled from #1 to #9, along with six additional cardiovascular segments—including one aortic arch, two cerebral aneurysms, one mesenteric artery network, and two abdominal aortic aneurysms—sourced from the Vascular Model Repository (VMR, Stanford University), identified using their original legacy names (Fig. 3B; fig. S17). This combined dataset encompasses a broad anatomical spectrum, ranging from straight arterial conduits to tortuous, branching, and aneurysmal geometries, thus establishing the system’s capacity to generalize across heterogeneous vascular topologies.

The SEBS-based vessel-on-chips exhibited high optical transparency suitable for live-cell imaging and maintained excellent dimensional integrity after thermoforming. Following endothelial seeding and static culture, confocal immunostaining revealed confluent monolayers throughout the vascular lumen. VE-cadherin staining delineated continuous intercellular junctions, while F-actin and nuclear markers confirmed cytoskeletal alignment and uniform cell distribution across bifurcation and curvature regions (Fig. 3B; fig. S17). These results demonstrate that the integrated fabrication platform robustly reproduces patient-derived and anatomically diverse vascular architectures with high geometric fidelity and reproducible biofunctionality.

### 2.3 Physical Twin: Patient-specific recapitulation of thrombosis pathophysiology

Blood perfusion experiments were performed on patient-specific vessel chips to establish a “physical twin” dataset recapitulating in vivo thrombosis mechanisms under controlled experimental conditions. We selected two representative patients (Patient #1, Patient #2) with distinct carotid geometries and performed comprehensive pharmacological perturbations (Fig. 4A).

CFD simulations revealed distinct shear rate distributions at the carotid artery bifurcations for patients 1 and 2 (Fig. 4, B and Q). For patient 1, we observed elevated shear rates at the site of endothelial injury (Fig. 4B). In contrast, patient 2 exhibited a low shear zone at the injury site (Fig. 4Q), highlighting the variable hemodynamic environments among patients experiencing vessel injury and plaque rupture.

Recalcified whole blood was perfused through patient-specific vessel chips at physiological shear rates. Controlled endothelial injury was induced at distinct anatomical sites—the internal carotid artery (ICA), and common carotid artery (CCA)—via laser ablation (Fig. 4, C to E, movie S3 and S4). ICA injury—characterized by high stenosis and complex flow patterns—triggered rapid platelet accumulation and unstable thrombus formation (Fig. 4C, t=7–9 min). In contrast, CCA injury sites with lower flow complexity exhibited more gradual, stationary thrombus development (Fig. 4D, t=7-9 min). We compared platelet signals at the thrombotic and healthy sites under physiological recalcification and found them to be similar (Fig. 4F). However, thrombotic response varied markedly by injury location, reflecting local hemodynamic and geometric effects. Quantification revealed significantly fewer platelet translocation events at the artificial injury site compared to the ulceration site (Fig. 4G), while the overall platelet accumulation volume remained similar (Fig. 4H).

We then evaluated the efficacy of various antithrombotic drugs in the Carotid Artery-Chip molded from Patient #1:

- Group 1, Anticoagulants: rivaroxaban exhibited greater inhibition of platelet signal, while apixaban showed limited effects in reducing platelet accumulation and translocation compared to controls (Fig. 4, I and J).
- Group 2, GPIb and VWF blockers: AK2 and ARC1172 significantly reduced platelet adhesion and translocation (Fig. 4, K and L) by disrupting the VWF-GPIb interaction.
- Group 3, αIIbβ3 antagonists: Abciximab and tirofiban exhibited strong inhibitory effects on platelet aggregation (Fig. 4M) and significantly reduced platelet translocation (Fig. 4N), demonstrating their effectiveness in preventing platelet-platelet interactions necessary for thrombus growth.
- Group 4, TxA_2_ pathway inhibitors: Aspirin showed no significant change compared to controls (Fig. 4, O and Q), consistent with previous reports that aspirin is less effective in preventing biomechanical platelet aggregation under high shear stress conditions(*67, 68*).
- Group 5, P2Y_12_ antagonists: Ticagrelor significantly reduced platelet accumulation and translocation (Fig. 4, O and Q).

For Patient #2, we observed distinct platelet behavior in the proximal and distal regions of the injury site. In the distal region, platelets accumulated faster and translocated multiple times, while in the proximal region, they formed more stable thrombi (Fig. 4R). Time-lapse fluorescence intensity measurements confirmed less platelet accumulation in the proximal ulceration site compared to the distal region (fig. S18). Statistical analysis showed a significant difference in platelet translocation events between these regions (Fig. 4S).

We then evaluated the efficacy of various antithrombotic agents in Patient #2 models. In contrast with our findings for Patient #1, anticoagulants (rivaroxaban and apixaban) both did not significantly alter platelet accumulation or translocation compared to control conditions (fig. S19, A and B). GPIb blockers (AK2 and ARC1172) and GPIIb/IIIa antagonists (abciximab and tirofiban) significantly reduced platelet adhesion and translocation in both patients (fig. S19, C to F). Among the antiplatelet agents, ticagrelor significantly reduced platelet accumulation and translocation, while aspirin showed no significant effect (fig. S19, G and H).

These spatiotemporal signatures varied systematically with patient geometry, injury site, and drug treatment, creating a diverse corpus for AI model training.

### 2.4 Digital Twin: Generative AI Model for Patient-Specific Thrombosis Prediction

To construct a digital twin that can predict thrombosis dynamics beyond experimentally tested conditions, we developed Cascade Learner of Thrombosis (CLoT), a conditional video diffusion model specialized for vessel-on-chip fluorescence microscopy (Fig. 5A). CLoT is built on a pretrained Diffusion Transformer (DiT) backbone and adapted to the thrombosis domain via low-rank adaptation (LoRA), enabling efficient finetuning while preserving the generative priors of the foundation model. The model is explicitly multimodal: patient-specific vascular geometry provides fixed anatomical context; endothelial injury is encoded as a spatial binary mask; and pharmacological perturbations are encoded through drug-condition prompts. These condition streams are separately embedded and fused through a decoupled cross-attention interface, allowing the geometry, injury location, and drug context to modulate all transformer blocks during diffusion denoising (Fig. 5A). This design enforces vessel-constrained synthesis and supports controlled generation of thrombosis videos under specified anatomic and experimental settings. A decoupled cross-attention module aligns these heterogeneous condition embeddings with the DiT latent sequence to modulate all transformer blocks during diffusion-based video denoising.

CLoT was trained on a curated physical-twin corpus of 491 high-quality thrombosis videos spanning multiple patient-specific carotid geometries, distinct injury sites, and 14 antithrombotic interventions (fig. S20). We introduced an automatic video quality check and prompt generation pipeline (fig. S21). The pipeline automatically excludes clips with unexpected conditions or poor quality, e.g., out-of-focus/blurry frames, debris-laden flow, incomplete endothelial coverage, or channel artifacts. By freezing the pretrained DiT backbone and optimizing only LoRA parameters inserted into attention and feed-forward projections, the finetuning process reduced trainable parameters by >95% relative to full model retraining, while retaining sufficient capacity to learn fluorescence microscopy statistics and hemodynamics-coupled platelet behavior. After training, CLoT generates thrombosis sequences conditioned on unseen combinations of vessel anatomy, injury site, and drug treatment, thereby functioning as a computational digital twin linked to the patient’s physical twin (fig. S22, movie S5).

### 2.5 Generative performance and biological fidelity

To comprehensively assess quality, we evaluated both general video fidelity and biologically grounded fidelity. To comprehensively assess quality, we evaluated both general video quality and biologically grounded fidelity.

#### Overall video quality evaluation

We quantified similarity between generated videos and biological ground-truth sequences using five independent replicates and five videos per method, benchmarking CLoT against six generators—Sora, Wan, Kling, Hunyuan, Hailuo, and Seedance (movie S6). To ensure biological plausibility, we required strict preservation of vessel anatomy and realistic hemodynamics. Accordingly, we assessed fidelity with a comprehensive battery spanning: pixel-level quality (PSNR, SSIM, MS-SSIM, LPIPS); semantic consistency (ViTScore, CLIPScore); spatial biological agreement (morphology of platelet distributions); temporal biological consistency (thrombus-growth kinetics; DTW); and spatio-temporal distributional alignment (see Methods). Metrics were min–max normalized across methods (lower-is-better metrics inverted) before aggregation.

Across all categories, CLoT matched or outperformed comparators and enclosed the largest area on the radar plot (Fig. 5B; Table 1), indicating balanced, multidimensional gains over commercial baselines. Sora produced smooth sequences and comparable temporal scores (0.3% vs. 0.3% MAE; 0.5% and 0.6% RMSE for CLoT and Sora) but showed weaker pixel perceptual agreement and frequent deviations from vessel anatomy, suggesting limited transfer from natural-video pretraining to hemodynamic scenes. In contrast, the LoRA-adapted CLoT mitigated these limitations, achieving a normalized DTW of ∼0.1% over 64-frame sequences while maintaining strong pixel and semantic fidelity (SSIM 0.94; CLIPScore 0.97), surpassing Sora (SSIM 0.62; CLIPScore 0.89). The fused circular plot (Fig. 5B) places per-metric bars around category sectors and encodes category means in the outer ring. CLoT encloses the largest radar area and dominates most family means, indicating balanced gains rather than a trade-off between pixel quality, motion realism, and biological plausibility. Representative generated frames for Patients #1 and #7 show qualitatively realistic platelet distributions, thrombus morphology, and spatiotemporal evolution (fig. S23-24).

**Table 1.** Multi-metric evaluation score comparison between CLoT and other commercial video generators based on 5 replicate generated videos for each generator. The best results are displayed as bold and the second is with underline.

| Metrics | CLoT | Sora | Wan | Kling | Seedance | Hailuo | Hunyuan |
| --- | --- | --- | --- | --- | --- | --- | --- |
| Pixel Perceptual-PSNR (↑) | **27.62** | <u>22.71</u> | 18.97 | 18.09 | 12.80 | 18.16 | 21.98 |
| Pixel Perceptual-SSIM (↑) | **0.94** | 0.62 | <u>0.85</u> | 0.68 | 0.4 | 0.64 | 0.83 |
| Pixel Perceptual-MS-SSIM (↑) | **0.95** | 0.63 | <u>0.86</u> | 0.66 | 0.39 | 0.65 | 0.82 |
| Pixel Perceptual-LPIPS (↓) | **0.05** | 0.29 | <u>0.15</u> | 0.27 | 0.45 | 0.26 | 0.17 |
| Motion Dynamic – EPE (↓) | **0.07** | <u>0.16</u> | 0.25 | 0.34 | 0.86 | 0.87 | 0.22 |
| Motion Dynamic - tSSIM (↑) | **0.90** | 0.84 | <u>0.87</u> | 0.85 | 0.73 | 0.83 | <u>0.87</u> |
| Motion Dynamic – flow cosine (↑) | <u>0.0009</u> | 0.0007 | **0.001** | -0.0001 | -0.0003 | 0.0005 | 0.0003 |
| Motion Dynamic – flow magnitude correlation (↑) | **0.40** | 0.07 | <u>0.17</u> | 0.15 | 0.03 | 0.06 | 0.07 |
| Motion Dynamic - t-cosine (RGB) (↑) | **0.32** | 0.13 | <u>0.23</u> | 0.21 | 0.18 | 0.15 | 0.18 |
| Temporal Bio-Pearson r (↑) | **0.61** | 0.46 | 0.30 | <u>0.49</u> | 0.38 | 0.39 | 0.23 |
| Temporal Bio-DTW distance (↓) | **0.0001** | <u>0.002</u> | 0.3131 | 0.058 | 2.6417 | 0.1711 | 0.0713 |
| Temporal Bio-MAE (↓) | **0.0003** | **0.0003** | 0.0018 | 0.001 | 0.0014 | 0.0007 | 0.0008 |
| Temporal Bio-RMSE (↓) | **0.0005** | <u>0.0006</u> | 0.0108 | 0.0024 | 0.0089 | 0.0039 | 0.0024 |
| Spatial Bio- Heatmap SSIM (↑) | **0.99** | 0.94 | 0.85 | 0.89 | 0.69 | 0.77 | <u>0.95</u> |
| Spatial Bio- Heatmap cosine (↑) | **0.26** | 0.05 | 0.13 | 0.12 | 0.06 | <u>0.14</u> | 0.02 |
| SpatioTemporal Bio-Cube cosine (↑) | **0.26** | 0.03 | <u>0.13</u> | 0.05 | 0.07 | 0.11 | 0.03 |
| Semantic-ViTScore (linear) (↑) | **0.94** | 0.87 | 0.85 | 0.76 | 0.61 | 0.8 | <u>0.91</u> |
| Semantic-ClipScore (linear) (↑) | **0.96** | 0.9 | <u>0.92</u> | 0.80 | 0.72 | 0.83 | 0.91 |
| Semantic-ViTScore (DTW) (↑) | **0.96** | <u>0.89</u> | <u>0.89</u> | 0.78 | 0.64 | 0.83 | <u>0.89</u> |
| Semantic-ClipScore (DTW) (↑) | **0.97** | 0.89 | <u>0.95</u> | 0.80 | 0.72 | 0.84 | 0.93 |

#### Biological fidelity evaluation

In addition to the overall video quality assessment, we next assessed whether CLoT-generated videos preserve both vessel anatomy and thrombosis biology at the pixel, spatial, and temporal levels, benchmarking against six state-of-the-art commercial video generators. Time-integrated “long-exposure” platelet-signal maps accumulated from 1–10 min revealed that CLoT maintained vessel-constrained, injury-localized platelet deposition patterns closely matching the physical twin, whereas commercial models frequently exhibited spatial drift, off-target platelet placement, or diffuse deposition inconsistent with the injury geometry (Fig. 5C, fig. S25 to S27). This indicates that domain-aligned conditioning and LoRA finetuning are necessary to anchor generative dynamics to patient-specific vascular context.

To further interrogate spatial fidelity over time, we extracted representative platelet-distribution snapshots at *t* = 3 and 9 minutes and overlaid segmented platelet masks onto the vessel boundary for direct multi-model comparison (Fig. 5D). Across three representative anatomical conditions—Patient #1 ICA ulceration, Patient #1 CCA artificial injury, and Patient #7 CCA injury—CLoT wireframe contours consistently colocalized with the filled ground-truth regions, reproducing both thrombus morphology and its evolution relative to the bifurcation geometry. In contrast, commercial models produced contoured fields that were frequently displaced from the injury zone, fragmented, or expanded into anatomically implausible regions (Fig. 5D). Quantification of spatial agreement using the overlap ratio confirmed that CLoT achieved substantially higher overlap with ground truth for all three conditions by a wide margin (Fig. 5E–G). These results demonstrate that CLoT learns geometry- and site-specific spatial priors governing platelet accrual.

Temporal thrombotic fidelity was evaluated by comparing mean platelet-signal intensity time courses derived from generated and experimental videos. For Patient #7, CLoT closely tracked ground-truth kinetics, capturing onset timing, accumulation rate, and platelet behavior, whereas other models displayed systematic temporal deviations including delayed activation, attenuated growth, or unstable plateaus (Fig. 5H). Importantly, embolic instability—a key pathological signature in carotid thrombosis—was also preserved: predicted embolization (platelet translocation) counts from CLoT were statistically indistinguishable from ground truth (ns), indicating that the model captures detachment dynamics rather than only bulk accumulation (Fig. 5I). Consistently, time-resolved platelet-area trajectories across all three conditions showed that CLoT maintained accurate growth profiles over the full 10-min assay window, while commercial generators diverged in either magnitude or temporal structure as the sequences progressed (Fig. 5J–L).

### 2.6 Patient-specific drug screening and personalized thrombosis risk stratification

Having established that CLoT reproduces untreated thrombosis phenotypes with high spatial and temporal fidelity (Fig. 5C–L), we next evaluated whether the digital twin can prospectively capture pharmacological modulation of thrombus growth and instability. We focused on two mechanistically distinct antiplatelet interventions that were not used as direct conditioning exemplars for the evaluated sequences: αIIbβ3 inhibition (abciximab) and P2Y₁₂ antagonism (ticagrelor). For each condition, CLoT was prompted with the corresponding drug context and injury-site mask on the patient geometry, and its predictions were compared against matched physical-twin ground-truth experiments (movie S7).

Across drug treatments, CLoT-generated platelet-signal trajectories closely recapitulated experimental kinetics (Fig. 5M). In the non-treatment setting, CLoT reproduced the rapid rise and sustained accumulation of platelet signal characteristic of the physical twin. Under abciximab, CLoT predicted near-complete suppression of thrombus growth, consistent with blockade of platelet–platelet consolidation at high shear. Under ticagrelor, CLoT predicted a marked attenuation of accumulation relative to non-treatment, matching the physical-twin reduction in ADP-amplified platelet recruitment (Fig. 5M). Importantly, this agreement extended beyond bulk growth to embolic instability, predicted embolization counts followed the same direction and rank order as ground truth, with both abciximab and ticagrelor reducing translocation events compared to non-treatment (Fig. 5N).

Together, these results demonstrate that CLoT functions as a pharmacology-aware digital twin capable of translating drug-specific perturbations into patient-specific thrombosis predictions. This proof-of-concept agreement with physical-twin assays supports the feasibility of coupling automated vessel-on-chip datasets with generative digital twins to guide personalized antithrombotic screening and thrombosis risk stratification.

## 3. Conclusion

We have demonstrated an integrated approach—combining automated high-throughput biofabrication with domain-specialized generative AI—that addresses longstanding barriers to personalized tissue engineering. The 100-fold increase in fabrication throughput (80 patient-specific chips in 10 hours versus ∼10 chips per week with manual methods) enables rapid production of patient cohorts sufficiently large to permit rigorous statistical validation and clinical translation. More significantly, the physical-to-digital twin paradigm enables quantitative, mechanistic prediction of thrombotic outcomes and drug efficacy without requiring repeated biological experiments. Our demonstration that LoRA-adapted video diffusion models achieve superior performance on biological consistency metrics compared to general-purpose foundation models (Sora, Wan, Kling, Seedance, Hailuo, Hunyuan) reveals an important principle: parameter-efficient specialization of pretrained models is more effective than both full retraining and direct zero-shot application to biomedical domains. This insight has immediate practical implications—researchers can download CLoT and rapidly customize it for new patient populations, geometries, or drugs through modest computational investment, democratizing personalized-medicine computational tools.

The benchmarking against Sora, Hunyuan, Kling, Seedance, Hailuo, and Wan reveals a critical insight: general-purpose video diffusion models, despite impressive performance on natural images, struggle to capture the specialized spatiotemporal dynamics of biological phenomena. The baseline model Wan, trained on diverse YouTube videos and synthetic CGI, achieves high pixel-level perceptual quality (SSIM 85% vs. CLoT 94% vs. Kling 68% vs. Seedance 39% vs. Hailuo 64% vs. Hunyuan 83%) but exhibits lower temporal consistency (DTW distance 31.31% vs. CLoT 0.01% vs. Sora 0.2% Kling 5.8% vs. Seedance 264%% vs. Hailuo 17.11% vs. Hunyaun 7.13%). This discrepancy suggests that natural-video pretraining encodes generic principles (edge preservation, color coherence, temporal continuity) but misses domain-specific constraints (platelet mechanics, hemodynamic coupling, coagulation cascade kinetics).

CLoT overcomes this via LoRA-adapted finetuning on 491 vessel-on-chip videos, requiring only ∼76.7M additional parameters and 4.65% GPU time compared to full-model retraining. This parameter efficiency matters strategically: it enables rapid deployment of specialized models without massive computational infrastructure, democratizing AI adaptation for under-resourced biomedical laboratories. The methodology is general; similar LoRA finetuning could generate specialized models for cardiac arrhythmia prediction, neural network activity patterns, hepatic drug metabolism, or kidney filtration dynamics—each trainable on modest video corpora from corresponding tissue-chip platforms.

Last but not least, our case studies (Patient 1 and Patient 2) illustrate the clinical utility of personalized thrombosis prediction. These divergent responses, not predicted from genetics or coagulation assays alone, highlight the importance of geometry-dependent hemodynamic coupling to pharmacological effects. In current practice, anticoagulant selection for stroke prevention in atrial fibrillation or post-stent thrombosis relies on population-averaged efficacy, with 3–8 % of patients experiencing inadequate protection or major hemorrhage (*69–71*). Personalized chips enable pre-treatment screening: measuring each patient’s thrombotic phenotype on their own vascular geometry, under their own anticoagulant candidate, predicts in vivo response with higher confidence than population averages. While our prospective validation cohort is modest, the 87 % similarity against ground truth demonstrates feasibility. Larger studies (n = 50–100) are required to establish clinical thresholds (e.g., which baseline platelet-adhesion values predict ischemic vs. hemorrhagic complications).

## 4. Methods

### 4.1 Automated vessel-on-a-chip manufacturing system

#### System Architecture and Design Principles

The automated vessel-on-chip manufacturing platform was designed as an integrated pneumatic–electromechanical system incorporating temperature, pressure, and vision-based positional feedback for real-time semi-closed-loop control. The system comprises five coordinated functional modules (pneumatic, film-feeding, thermoforming, puncher, and alignment) operating sequentially under coordination by a control module to complete the full fabrication cycle: hot embossing of patient-specific vascular geometries, formation of perfusion inlets/outlets via automated punching, and optical alignment and bonding of the two channel halves. All modules were mounted on a custom-designed aluminum frame (dimensions: 30 cm × 12.5 cm × 15.8 cm) to ensure structural rigidity, geometric precision, and modular accessibility for maintenance and calibration (fig. S1). Electronic control was managed by a commercial 3D printer controller board (Azteeg X3 Pro) running custom firmware, with centralized MATLAB-based host interface coordinating stepper motors, pneumatic solenoids, temperature controllers, and camera systems (fig. S16).

#### Film-Feeding Module

The film-feeding unit consisted of an active motor-driven roller (5 mm diameter ZrO₂ ceramic rod; rotational speed 0–100 rpm) and a compliant passive roller (12 mm diameter stainless steel) mounted on a 3D-printed flexible frame incorporating compliant hinges (Fig. 2E, upper left). As the SEBS film (Flexdym™, Eden Tech; 25 m length, 170 mm width supplied, trimmed to 20 mm width, 150–200 μm thickness) passed between the rollers, the compliant frame deflected upward under material tension, generating a stable downward force that pressed the film against the active roller to prevent slippage. The active roller was driven by a stepper motor (SM-42BYG011-25, Mercury Motor) and precisely advanced or retracted the film via programmed step sequences, providing accurate film position control throughout the system.

#### Pneumatic Module

The pneumatic circuit employed a dual-stage pressure-regulation system with independent low-and high-pressure branches (fig. S7A). A main regulator reduced the primary air supply (from compressor) to approximately 5 bar, which was split into two pathways through a Y-shaped junction. In the first pathway, a secondary regulator further reduced the air to ∼1 bar and directed it through a check valve to serve as the low-pressure line. The second pathway retained 5 bar pressure and was controlled by a solenoid shut-off valve. Both branches recombined through a second Y-junction before entering a 5/2-way solenoid valve (6V0510M5B050, AirTAC) connected to a pneumatic cylinder (CQ2KB50-40DZ, SMC). This architecture enabled seamless switching between high- and low-pressure operation during thermoforming, yielding a smooth soft-start/soft-stop motion profile that protected the glass mold from mechanical impact while maintaining stable, uniform compression during the hot-pressing cycle.

#### Thermoforming Module

The thermoforming module consisted of custom-machined brass heat blocks (thermal conductivity: 109 W/m·K; specific heat: 380 J/kg·K) designed for precise and uniform thermal performance. The bottom heat block (dimensions: 80 mm × 35 mm × 11.7 mm with 1.7 mm cavity depth) was fitted with a single 24 V, 40 W heating cartridge (unbranded, 24V, 40W), while the top heat block (dimensions: 60 mm × 35 mm × 10 mm) incorporated two identical cartridges to ensure faster and more homogeneous heating. Each block was instrumented with a K-type thermocouple (RS PRO 268-4513) connected to a PID temperature controller (RS Pro 222-8114) for closed-loop thermal regulation (fig. S2, A and D).

The top heat block was mounted to the pneumatic cylinder through a 3D-printed thermal bridge (material: FormLabs™ Rigid 10K resin; thermal conductivity: 0.621 W/m·K) providing both vertical motion and heat isolation. The bottom heat block was attached to the aluminum frame via a thermal-insulation pad (material: aluminum silicate ceramic fiber board; thickness: 6 mm; thermal conductivity: 0.132 W/m·K) to minimize conductive heat transfer and enhance temperature stability (fig. S2A).

The bottom heat block featured an integrated vacuum cavity sealed by an O-ring (red silicone), enabling vacuum-locking of the 3D-printed glass mold during thermoforming. The block integrated dual internal channels for circulating cooling water (flow rate: 1100 mL/min) supplied by a miniature diaphragm pump (JSB1523018, TCS Tec). The pump was automatically activated by the PID controller when the bottom block temperature exceeded 85 °C, maintaining the block within the 75–85 °C target range (fig. S4C). Water was supplied and circulated from a glass beaker placed in room temperature.

The SEBS film was loaded as a sandwich structure between an FEP (fluorinated ethylene propylene) release film and a glass slide containing the 3D-printed mold (borosilicate glass, dimensions: 75 mm × 25 mm, thickness: 1 mm). The cavity depth of the bottom heat block and glass-slide thickness jointly determined the final channel height and wall thickness of the vessel chip, tuned to remain within the 600 μm working distance of the microscope objective (Olympus LUCPLFLN 10X air objective) used for downstream imaging (fig. S2B).

#### Thermoforming Cycle

The fabrication process begins with loading the glass mold slide into the bottom heat block, aligned such that its centerline matches the midpoint of the bottom block. Then the slide is vacuum locked to prevent movement and ensure intimate thermal contact during forming (fig. S2C).

The complete thermoforming cycle comprises three consecutive steps: Pre-Heating, Hot-Press, and Release (fig. S7B). The detailed process inside each step is introduced below:

##### Pre-Heating (Step 1; Duration: 20 s)

Low-pressure air (∼1 bar) from the pneumatic system actuates the pneumatic cylinder, gently driving the top heat block downward at ∼35 mm/s. As the film contacts the vacuum-fixed glass slide, a sandwich structure forms, entering the pre-heating phase. The SEBS film is heated above its glass-transition temperature (Tg ≈ 100 °C) under approximately 130 N of compressive force, softening the polymer for molding.

##### Hot-Press (Step 2; Duration: 40 s)

The shut-off valve opens, allowing high-pressure air (∼5 bar) to enter the pneumatic cylinder, generating approximately 950 N of compressive force. This force fully presses the molten SEBS into the mold cavities with consistent, uniform pressure distribution. Internal pressure is monitored via a pressure transducer (range 0–10 bar, ±0.5% accuracy) and logged continuously.

##### Release (Step 3; Duration: ∼10 s)

The solenoid valve reverses airflow, exhausting the high-pressure line and supplying low-pressure air to retract the top heat block smoothly at ∼35 mm s⁻¹. The formed SEBS film is then cooled under ambient air via natural convection for approximately 10 s. During this period, the film temperature drops below its glass-transition point, shifting the material behavior from viscoelastic to elastic-dominant. As the polymer solidifies, the SEBS retains its molded geometry with high dimensional fidelity.

##### Film Demold

After cooling, the film-feeding module advances the strip outward, peeling the formed SEBS cleanly from the mold via controlled speed (∼2 mm/s). The resulting half-channel geometry (thickness: 0.7 mm, channel height: ∼150 μm, wall thickness: ∼500 μm) proceeds to the puncher module.

#### Puncher Module

The puncher module converts servo-driven rotary motion into linear displacement through a custom-designed crankshaft linkage (fig. S10, A and B). The crankshaft mechanism was 3D-printed in-house (Form 3+, Formlabs; material: Rigid 10k) and driven by a high-torque servo motor (HB7545GS; peak torque: 450 N·cm). The crankshaft translates rotary motion into linear displacement along two precision linear guide rails (MGN7C, HIWIN), driving dual biopsy punch assemblies (KAI Medical; punch diameter: 3 mm; customizable spacing). Punch holders and cutting mats are modular and interchangeable to accommodate different mold layouts.

The servo motor speed and phasing are adjustable to tune the cutting period (time interval during which the punch blade engages the film), enabling optimization for SEBS films of varying thickness (500–3000 μm). A manually adjustable screw mechanism (thread pitch: 0.7 mm; adjustment range: 8 mm) allows precise control of the gap between the punch head and cutting mat, enabling repeatable, consistent cutting depth across production runs (fig. S10C). Punch depth is calibrated at system startup by monitoring film penetration depth.

The puncher does not fully perforate the film but instead cuts to a controlled depth (∼90 % film thickness), leaving the punched segment partially attached. This design prevents loose debris from accumulating within the punching chamber, eliminating the need for manual material removal and substantially reducing maintenance requirements (fig. S11).

After punching, the patterned film advances to the alignment module via stepper-motor through step-counting position control.

#### Alignment Module

The alignment module integrates five motorized axes arranged as a compact 3D gantry providing full positional control of the two vessel halves during folding and bonding (fig. S12).

##### Tension Rollers

Two tension rollers are driven by stepper motors (station roller: 20HS34 stepper motor, HANPOSE; motive roller: PG15S-D20-HHB9 stepper motor, NMB Tech) via friction wheel made with silicone rubber to apply stretching forces to tension the SEBS film. The friction wheel also acts as a “clutch” allowing slippage during the pre-alignment phase to prevent over-stretching and film rupture (fig. S12A). Roller diameter: 12 mm; active surface: ground stainless steel. Roller motors operate at 0–30 rpm, generating linear surface velocities of 0– 15.7 mm/s. These rollers subsequently function as vertical adjustment elements in coordination with the feeding roller during alignment, and later act as compression and delivery rollers to consolidate and feed the aligned chip to the user.

##### Film-Feeding Roller

The feeding roller, driven by a stepper motor (Creality 42-40), transmits motion via a timing belt (GT2, 5 mm wide) and pulley system. The active feeding roller incorporates a compliant mechanism similar to the film-feeding module, accommodating variations in film thickness and minimizing slippage.

##### Folding Axis

The moving stage (folding stage) is actuated by a stepper motor (17LS13-0704J-120E) coupled to a precision lead screw with linear translation guided by dual precision rails (MGN7C, HIWIN; length: 100 mm) for smooth, stable motion. Travel range: 60 mm; positioning resolution: 12 μm.

##### Horizontal Alignment Axis

A secondary horizontal stage (LX30-L, MISI; travel range: 10 mm) driven by a miniature stepper motor (Nidec; step resolution: 2 um) provides micrometer-level lateral adjustment perpendicular to the primary folding motion, enabling precise 2D registration of vessel halves (fig. S12).

##### Film Grabber

A solenoid-actuated, 3D-printed film grabber (material: FormLabs™ Rigid 4k; solenoid: 5 V, 7 W) mounted on the moving stage securely holds the SEBS film during folding and alignment operations via clamping jaws (contact force: ∼0.85 N).

#### Alignment Procedure

The alignment workflow proceeds through pre-alignment, alignment, and post-alignment phases:

##### Pre-Alignment (fig. S13A)

As the moving stage approaches the stationary tension rollers, the film grabber opens (solenoid energized, jaws released), while the feeding roller simultaneously advances the SEBS film from the upstream puncher module. When the film reaches the grab zone, the grabber closes (solenoid de-energized) to secure the film. The grabber then moves outward by 40 mm, synchronized with the feeding roller to prevent slack formation or overstretching. As the stage retracts (moving inward), the film folds while the feeding roller continues advancing, creating an upward arch. When the folded segment contacts the dual tension rollers, they rotate inward, generating an upward force through friction that tensions and stabilizes the film (fig. S13A, step III to IV).

The camera is pre-focused on the tangent plane between the feeding roller and the stationary tension roller. Once the film is properly tensioned, the section nearest the stationary roller aligns with the camera’s focal plane, allowing clear visualization of the vessel geometry and alignment markers (fig. S15). The rollers halt automatically once these features are recognized by the onboard vision system. During this process, the friction wheel functions as a clutch, permitting controlled slippage when the film is fully engaged and stretched, thereby preventing mechanical stress or tearing. This sequence completes the pre-alignment phase and prepares the system for active optical alignment.

##### Alignment (fig. S13B)

The vessel region contains four L-shaped alignment markers (printed as part of the SEBS mold; marker size: 300 μm × 300 Μm; thickness: 100 um; spacing: 2 mm) positioned around the area of interest to assist computer-vision-based registration. A single white illumination LED mounted on the folding mechanism illuminates the vessel area, and a reflection mirror directs the light path to an integrated CMOS camera (HY-800B, HAYEAR; resolution: 4k, 8 megapixels; frame rate: 25 fps; sensor size: 1/1.8 inch) (Fig. 2E, *lower right*, fig. S12 and S13). The camera captures both top and bottom channel halves through reflection imaging, revealing initial misalignment.

Real-time image processing via OpenCV-based software identifies alignment markers in both reflected images, calculates the spatial offset (Δx, Δy, Δθ) between the two halves, and computes corrective motor commands. The controller then directs coordinated 2D motion: the station tension roller and feeding roller produce vertical adjustments (fig. S13B, III and V), while the horizontal stage provides lateral motion (fig. S13B, IV). After each correction, the controller advances the two halves slightly closer together, which may introduce transient misalignment; this closed-loop process repeats until the spatial offset falls below a threshold (±20 μm alignment tolerance) and the two halves contact, completing the alignment sequence (fig. S14 and S15).

##### Post-Alignment (fig. S13C)

Following successful alignment, the moving stage advances continuously for 3 mm, driving the two tension rollers to compress the aligned chip (force ∼30 N). Once compression begins, the film grabber releases (solenoid energized, jaws open), and the tension rollers, together with the feeding roller, advance the aligned chip upward. During this process, the chip is further compressed between the tension rollers, which removes interfacial air bubbles and ensures uniform bonding between the two vessel halves. The completed chip is then manually separated from the film by the user, while the residual film retracts automatically to its initial position, resetting the system for the next alignment cycle (movie S1).

Sub-micron alignment accuracy is achieved through iterative closed-loop correction, with typical alignment requiring 3–5 feedback cycles per device.

#### System Integration and Real-Time Control

All motion and process control are managed by the custom Azteeg X3 Pro controller board running custom firmware developed in Arduino IDE. The controller board governs stepper motors (via DRV8825 drivers), pneumatic solenoids (via relay modules), film-grabber solenoids, and pressure regulators through serial communication with the MATLAB-based host interface. Temperature regulation is handled by dedicated PID controllers (one for top block, one for bottom block). Positional feedback is derived from calibrated step counts determined during system calibration routines performed at startup. The alignment camera is integrated into the MATLAB control environment, enabling synchronized image acquisition and motion commands during the critical alignment phase.

A safety interlock system monitors all pneumatic, thermal, and electrical parameters. If any parameter deviates outside safe operating bounds (e.g., excessive pressure > 6 bar, temperature >200 °C, motor stall detection), production is halted and the system returns to a safe resting state.

### 4.2 Patient-Specific Vascular Geometry Acquisition and 3D Reconstruction

#### Clinical imaging and image processing

Clinical and anatomical vascular data were obtained from two complementary sources: (i) computed tomography angiography and digital subtraction angiography (DSA) scans of three patients with carotid artery–related cerebrovascular disease at Royal Prince Alfred Hospital (RPAH), and (ii) four additional vascular geometries (cerebral aneurism, aorta, mesenteric artery, and abdominal aneurism) from the Vascular Model Repository (VMR, Stanford University), an open-access database of validated cardiovascular models (*73*). Together, these datasets provided a total of fifteen vascular geometries, enabling the generation of patient-specific and reference vascular models for experimental fabrication and AI training.

Patient inclusion criteria for blood perfusion experiments were based on the availability of high-quality CTA and DSA datasets (New South Wales Telestroke Service) suitable for accurate 3D reconstruction and hemodynamic analysis. Patients with severe systemic disease, coagulation abnormalities, or prior vascular interventions were excluded to ensure consistency across cases. All clinical data were collected in accordance with institutional ethical guidelines and patient consent protocols approved by the Sydney Local Health District Human Research Ethics Committee (HREC–RPAH Zone; X23-0267 and 2023/ETH01607).

#### Acquisition and Processing of Clinical CTA Images

CTA imaging was performed using a GE Optima CT660 scanner following standardized protocols to ensure reproducibility. Scans were acquired with a slice thickness of 0.625 mm and pixel spacing of 0.539 × 0.539 mm. Each dataset was enhanced using an 80 mL bolus of iodinated contrast agent (4–7 mL s⁻¹) followed by saline flush to optimize lumen definition and detect stenotic or ulcerative lesions. Images were transferred to a secure research repository for post-processing.

CTA datasets were imported into SimVascular (Cardiovascular Biomechanics Computation Lab, Stanford University) for 3D reconstruction. Vessel lumens were initially delineated using region-growing algorithms, followed by manual refinement to correct artifacts and calcification-induced discontinuities under clinical supervision. Ulcerations and lumen boundaries were validated for continuity using slice-wise inspection tools. The final 3D reconstructions were exported as STL files for 3D mold design and fabrication preparation.

#### 3D Mold Design and Fabrication

The 3D mold reconstruction was performed as previously described (*26–28*). In brief, the STL mesh of the patient-specific vessel geometry was first imported into computer-aided design (CAD) software (SpaceClaim, Ansys) for surface smoothing and model reconstruction. The reconstructed model was then imported into a secondary CAD environment (Fusion 360, Autodesk) for pre-print modifications, which included the following:

Vessel segmentation and mirroring: Vessel branches were elongated and bisected along the mid-plane. The resulting two half-vessel geometries were mirror-positioned along the central axis (each 13 mm in length, offset by 8.5 mm from the centerline).

Alignment markers: Four L-shaped alignment markers (300 × 300 μm area; 100 μm width; 80 μm depth) were positioned symmetrically around the vessel region to assist optical alignment. Perfusion inlet/outlet markers: Two circular inlet and outlet markers (3 mm diameter, 13 mm apart) were placed at either end of the vessel to define perfusion cutout locations.

The finalized design was converted into a digital light processing (DLP) printing format with an XY resolution of 10 μm and a Z resolution of 5 μm (VoxelDance Additive 4.0). The detailed 3D printing method was performed as previously described (*28*).In brief, molds were fabricated using a DLP micro 3D printer (microArch® S240, BMF) with high-temperature resin (HTL) printed directly on chemically treated soda-lime glass substrates. Post-print curing was performed under UV exposure at 80 °C to enhance mechanical stability and surface hardness. The mold height (typically ∼150 μm) was optimized to produce a vessel-chip wall thickness of ∼600 μm after thermoforming, balancing optical transparency for microscopy with sufficient structural rigidity.

### 4.3 Endothelial cell culture and seeding

#### Cell Preparation

Human umbilical vein endothelial cells (HUVECs, P5-9) were obtained from Lonza Bioscience (Lonza, C2519A). Cells were cultured in complete EGM-2 medium (Lonza, CC-3162) supplemented with hydrocortisone, hFGF-B, VEGF, R3-IGF-1, ascorbic acid, hEGF, gentamycin, and amphotericin B, maintained at 37 °C and 5 % CO₂. Cells were passaged when reaching 80–90% confluency. Medium was changed every 2 days.

Prior to seeding into vessel chips, cells were detached from T75 culture flasks (Corning) using TrypLE express enzyme (Thermofisher; incubation time: 2 min at 37 °C) and resuspended in EGM-2 medium at a concentration of 1× 10^7^ cells/mL. Approximately 2 μL of cell suspension (yielding 2 x 10^4^ cells per chip) was injected into the vessel channel via a 10 μL pipette tip positioned at a 45° angle (Fig. 3A).

#### Seeding Protocol

For chips with complex branching geometries, resistance-dependent preferential flow was observed, with most cells flowing into the lower-resistance branch. In such cases, a second injection was performed from the opposite inlet/outlet to ensure confluent coverage across all branches. Residual cell suspension was removed from inlets/outlets to prevent unintended fluid flow. Chips were placed in 90 mm petri dishes lined with damp paper towels and sealed with parafilm, then incubated for 30 min at 37 °C in 5 % CO₂ to allow initial cell adhesion.

To promote confluent endothelial coverage on both halves of the circular channel, double-sided tape was used to attach the glass slide holding the channel to the bottom of the petri dish. Chips were then flipped upside-down in-place, and the injection procedure was repeated for the second channel half. Residual cells were washed with warm EGM-2 and inlets were filled with fresh medium. Chips were incubated overnight (typically 16–18 hours) at 37 °C in 5 % CO₂ to achieve confluent endothelial monolayer formation before downstream experiments.

#### Immunofluorescence Staining

Following the designated culture period, chips were fixed with 4 % paraformaldehyde (PFA; Electron Microscopy Sciences) for 10 min at room temperature. Cells were permeabilized with 0.5% (v/v) Triton X-100 (Sigma-Aldrich) in PBS for 10 min. Non-specific binding was blocked with 5 % bovine serum albumin (BSA; Sigma-Aldrich) in PBS for 30 min at room temperature.

Primary antibody against VE-cadherin (1:200 dilution, Thermofisher 53-1449-42, mouse monoclonal, clone 16B1) was applied in 2.5 % BSA/PBS and incubated for 1 hour at room temperature. Following three washes (5 min each) with PBS.

Phalloidin 555 antibody (1:5000 dilution, Thermofisher A30106) for F-actin visualization and nuclear counterstain (Hoechst 33342, 1:3000 dilution, Thermofisher H1399) were applied sequentially for 30 min and 10 min, respectively, at room temperature, with PBS washes between steps. Inlets were filled with PBS to prevent channel dehydration during imaging.

#### Fluorescence microscopy

Imaging was performed with an Olympus IX83 inverted microscope equipped with a Fluoview FV3000 fluorescent confocal scanning unit (Olympus, Japan) and a Plan Apo 10X air lens (NA: 0.4). Z-stack acquisition was done with 5 μm optical section thickness for a total Z-range of 300 μm, covering the full vessel cross-section. Z-stacks were projected using maximum intensity for analysis..

### Blood perfusion and thrombosis assay

#### Blood collection and handling

Blood collection from healthy donors was approved by the University of Sydney Human Research Ethics Committee (HREC, project 2023/HE000582). Donors provided written informed consent and were screened for appropriate age, weight, and absence of anticoagulant or anti-inflammatory medication use. Blood was collected via a 19G butterfly needle into a syringe containing 3.8% sodium citrate anticoagulant. Whole blood was stained with anti-CD41 and anti-fibrin antibodies. Immediately prior to perfusion experiments, whole blood was recalcified by adding CaCl₂ (Sigma-Aldrich; final concentration: 2 mM) to restore coagulation cascade function. Blood temperature was maintained at 37 °C throughout handling and experimentation.

#### Endothelial injury induction

Localized thrombosis was induced by laser ablation on the Carotid Artery-Chip at user-specified locations (ICA, CCA, or ECA regions of the bifurcation). A RAPP UGA-42 Caliburn 355/42 module (355 nm pulsed DPSS; 42 µJ pulse⁻¹ at 1 kHz; 42 mW average) was coupled to the confocal microscope and operated at 70% output. The beam traced a short linear path over the ulceration region; power, dwell time and spot size were fixed across samples. Injury magnitude was standardized to loss of ≈5–7 endothelial cells; chips outside this range were excluded. Immediately after ablation, recalcified whole blood was perfused under patient-scaled flow, and platelet accrual/embolization were recorded in real time by confocal imaging. Timing: ablation was performed at t = 0 min, immediately prior to blood perfusion initiation.

#### Perfusion setup and hemodynamic conditions

Recalcified blood was perfused through patient-specific vessel chips via a programmable syringe pump (Harvard Apparatus PHD 2000; flow rates: 50–130 μL/min, corresponding to physiologic shear rates of 830 s⁻¹ depending on vessel geometry and cross-sectional area).

Shear rate (γ^•^) was calculated from flow rate (Q) and channel cross-sectional area (A) via: γ^•^ = Q / (A × d), where d is the channel depth. For patient-derived bifurcations, cross-sectional area varies spatially; local shear rates were estimated from computational fluid dynamics simulations (ANSYS Fluent; see below).

#### Pharmacological interventions

Anticoagulant or antiplatelet drugs were added to recalcified blood at therapeutic concentrations 30 minutes before perfusion. Rivaroxaban was obtained from Sigma Aldrich, #SML2844-5MG, and reconstituted in DMSO to 4.59 µM. Apixaban was obtained from Cayman Chemical, #15427, and reconstituted in DMSO to 10.88 mM. AK2 was obtained from Sapphire Bioscience, #GTX76458-250UG, 1 mg/mL. Aptamer ARC1172 was chemically synthesised (IDT DNA Tech., Coralville, USA) to 1 mM. Tirofiban was obtained from Sigma Aldrich, #30165, and reconstituted in saline to 0.25 mg/mL. Abciximab was obtained from Invitrogen, #MA5-47865, 1 mg/mL. Aspirin (acetylsalicylic acid) was obtained from Sigma Aldrich, #A5376-1KG, and reconstituted in ethanol to 100 mM; Ticagrelor was obtained from Sigma Aldrich, # SML2482-50MG, and reconstituted in DMSO to 1 mM. Clexane was obtained from Sanofi Medical, 100 mg/mL (10,000U). Prostaglandin E1 and Prostacyclin were obtained from Sigma Aldrich. Indomethacin was obtained from ChemSupply Australia #GA3090-25G, and reconstituted in DMSO to 20 mM. MRS2179 was obtained from Sapphire Bioscience, #10011450-10MG, and reconstituted in DMSO to 100 mM. Control experiments used recalcified blood without pharmacological agents.

#### Platelet Staining and Imaging

To visualize thrombotic events in real time, blood samples were pre-labeled with a fluorescent platelet marker (CD41 antibody, 1 μL per mL blood; incubation: 15 min at 30 °C in the dark prior to recalcification).

An Olympus FV3000RS confocal microscope with a 10× UPlanXApo objective was used for real-time imaging. The focal plane was set 30 μm above the bottom of the vessel lumen. Platelet and fibrin signals were acquired using the default pinhole size, while the CD31 signal was observed using an 800 μm pinhole for comprehensive endothelial wall visualization. Quantitative analysis of fluorescently labeled platelets and fibrin was performed frame-by-frame using IMARIS software (Bitplane AG, Oxford Instruments). Video acquisition: frame rate 0.17 fps; acquisition duration: typically 10 min per experiment.

#### Hemodynamic Simulation and Local Shear Rate Estimation

Patient-specific carotid geometries (STL files used for chip fabrication) were imported into ANSYS Fluent 2020 R1 for pre-processing, meshing and simulation. A tetrahedral-dominant mesh (global max element size 50 µm) was generated with local refinement at the bifurcation (sphere-of-influence, 5 µm elements). Inflation layers were applied on vessel walls (growth rate 1.2; up to 5 layers) to resolve near-wall shear.

Blood was modeled as an incompressible Newtonian fluid (ρ = 1060 kg m⁻³; μ = 0.00345 Pa·s). Steady, laminar flow was assumed. Velocity inlets were prescribed from patient flow estimates; pressure outlets were set at ICA and ECA; walls were rigid with no-slip. Simulations used the pressure-based solver with SIMPLE coupling and second-order upwind spatial schemes. Convergence required residuals <10⁻⁶ and outlet mass-flow balance within 0.1% of the inlet.

To determine microfluidic operating conditions, we first simulated the in-vivo-scale model using a representative human CCA flow to obtain the target shear-rate field. We then calculated the chip’s initial inlet flow from its CCA branch diameter to match bulk shear. The microfluidic model was run and the inlet was iteratively adjusted until the overall shear-rate distribution closely reproduced the in-vivo pattern. This optimized inlet was used in experiments.

Post-processing extracted shear-rate maps and velocity streamlines to visualize regions of disturbed flow and elevated gradients. These outputs validated that the chips reproduced patient-relevant hemodynamics and guided placement of laser-injury sites for thrombosis assays. Correlating the local shear environment with platelet accrual and drug responses enabled mechanistic interpretation across patient geometries.

### AI Model Architecture and Training

#### CLoT (Cascade Learner of Thrombosis) architecture

CLoT builds upon a conditional video diffusion model architecture based on a Diffusion Transformer (DiT) backbone pretrained for natural video synthesis (Wan; Alibaba; pretraining billions of videos and images from public datasets and resources from Internet). The model accepts four categories of input:

1. <u>Vascular Geometry Context:</u> We initialize the first frame from patient-specific vascular geometry (mesh/2D projection) and encode the full clip using Wan-VAE. Wan-VAE compresses spatiotemporal dimensions by 4×8×8 (time×height×width), yielding a latent of shape [1+T/4,H/8,W/8,C] with C=16 channels; the first frame is only spatially down sampled to preserve image fidelity. We use 512×512 resolution videos generated from blood perfusion experiments.
2. <u>Multimodal Conditions:</u> - Injury Site: Encoded as a binary mask image overlay on the geometry image (spatial resolution matching video frame size: 512×512 pixels; values: 1 = injury location, 0 = healthy vessel) processed via CLIP image encoder. - Drug Treatment: The drug effect is encoded in text embeddings.
3. <u>Text Prompts:</u> Instructions such as “*The video shows a 2D microscopy image of a patient’s carotid artery bifurcation. The single larger branch is the common carotid artery, which lies upstream, the two branches which branch from the bifurcation are the internal and external carotid artery respectively, which lie downstream. The video has a black background, and the vessel shape is visualized using green meshes, which represent endothelial cells lining the vessel walls. There is a laser cut injury on the vessel wall, which is visualized with a black gap in the green meshes, where endothelial cells have fallen off. Blood is flown through the vessel structure, red fluorescent signals representing platelets, from the single common carotid branch to the branched internal and external carotid branches. Platelets accumulate and detach at the laser injury zone and regions cell fallen off*” are embedded via a pretrained text encoder (umT5; multilingual Universal Transformer T5; output: 4069 dimensional embedding).
4. <u>Timestep Information:</u> Diffusion timestep t ∈50 is embedded via sinusoidal positional encoding and passed to all transformer blocks.

#### Backbone architecture

Tokenization / patchification. We operate in the VAE latent space. Given an input clip of T frames at resolution H×W (we use H=W=512 in experiments), the Wan-VAE compresses time by 4× and space by 8×, producing a latent grid 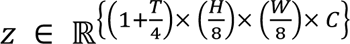 with C=16. We then tokenize z using a 3D convolution with kernel/stride (1,2,2) (no temporal down sampling), flattening to a sequence (B, L, D) with

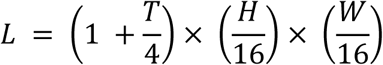

This preserves the VAE’s temporal resolution while matching a 16×16 effective spatial stride. Tokens are linearly projected to the model width D_model=5120.

#### Diffusion Transformer

The denoiser is a Diffusion Transformer (DiT) with N=40 blocks. Each block applies (i) multi-head self-attention with n_h=40 heads (head dim 128), (ii) cross-attention that fuses conditioning streams (text and, when present, image/video conditions), (iii) layer normalizations, and (iv) a feed-forward network with hidden size 13,824 and GELU activation. Timestep embeddings are injected into every block. This design supports long prompts and rich conditional control while maintaining high throughput in latent space.

#### Unpatchification / decoding

The transformer output sequence is reshaped back to the latent grid and decoded by the VAE to full-resolution frames H×W for T steps (e.g., 512×512, T=65 in our main experiments).

#### Finetuning via LoRA

Motivation. A Wan-pretrained video backbone does not directly capture fluorescence microscopy statistics or vessel hemodynamics. We therefore adapt DiT with lightweight Low-Rank Adaptation (LoRA), keeping all base weights frozen.

#### Parameterization

For a weight W ∈ ℝ^{d_out_× d_in_}^, LoRA introduces a low-rank update:

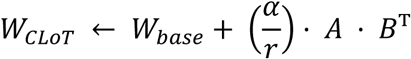

with rank r=4 and scale α=4 (effective scale α/r=1). Here *A* ∈ ℝ^{*d_out_*× *r*}^ and B ∈ ℝ^{d_in_× r}^. We up-cast LoRA parameters to FP32 for stability while keeping forward/backward in bfloat16.

LoRA is injected into every DiT block’s attention projections (q, k, v, o) and MLP linears (the first and last FFN linears, i.e., ffn.0, ffn.2). This yields a small, fixed overhead yet provides sufficient capacity to bridge the domain gap.

#### Training objective and schedule

We use standard diffusion training with a 1,000-step noise schedule. At each iteration, we sample t, perturb the latent x₀ to x_t, and optimize a weighted MSE:

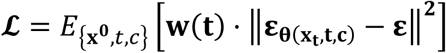

where c denotes conditioning (text, and optional image/video features). Optimization uses AdamW (learning rate 1×10⁻⁵). We freeze the VAE and all encoders as well as all base DiT weights; only the LoRA parameters are trainable. Optional gradient-checkpointing/offload can be enabled for memory savings without changing the formulation.

#### Implementation notes (for reproducibility)

- Model width 5120; heads 40 (head dim 128); FFN 13,824.
- VAE latent channel C=16; VAE compression 4× (time) and 8×8 (space).
- Patchify kernel/stride (1,2,2); token length *L* = (1 + *T*/4) × (*H*/16) × (*W*/16).
- Text encoder output shape (B, 512, 4096).
- Main experiments: H=W=512, T=65 frames.

#### Training data and corpus composition

We deployed a fully automated video-quality assessment (VQA) pipeline to filter experimental videos prior to model use (fig. S20). Videos exhibiting poor illumination, debris-related artifacts, or aberrant temporal dynamics were removed. Candidate prompts were produced by an LLM and vetted by human experts, ensuring that only high-fidelity videos—and their standardized prompts—were included in the training, validation, and test partitions. Among 712 videos, we collected 546 qualified videos for training and validating our model.

#### Training Procedure

– Optimizer: AdamW (learning rate: 1.0× 10⁻5;)
– Learning Rate Schedule: Cosine annealing with warm-up (total epochs: 8)
– Batch Size: 2 videos per batch given limited GPU memory
– Hardware: 2 GPUs (model: NVIDIA GeForce RTX 4090; total memory: 24 GB per GPU)
– Training Duration: 12 hours per epoch; total training: 8 epochs = 4 days

#### Inference Procedure

At inference, users provide:

1. Patient-specific vessel geometry
2. Injury site (ICA/CCA/ECA)
3. Drug treatment (or untreated)
4. Desired video duration (number of frames)
5. Natural-language prompt for additional context

The CLoT model performs iterative denoising starting from Gaussian noise *x*_*T* ∼ *N*(0, *I*), progressively refining via:

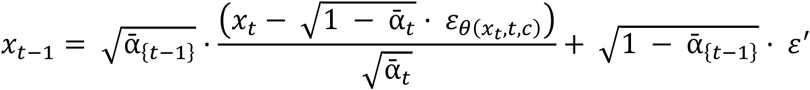

where ε^′^∼ *N*(0, *I*) and the denoising schedule follows standard DDPM (Denoising Diffusion Probabilistic Models) conventions.

#### Inference speed and latency

Video generation requires T = 50 denoising steps (sampling 5–10 % of the standard 1000-step diffusion schedule via consistency distillation or guidance-based acceleration), yielding inference times of 10 minutes per video on a single NVIDIA GeForce RTX 4090 GPU.

### Quantitative Evaluation of AI-Generated Thrombosis Videos

#### Video Dataset Composition

The evaluation dataset consisted of fluorescence time-lapse microscopy videos acquired during platelet activation following laser-induced endothelial injury in endothelialized, patient-specific vessel-on-chip devices. Experimental ground-truth data comprised three biological replicates. AI-generated videos were produced using our CLoT model and six state-of-the-art commercial generative video models—Sora, Hailuo, Hunyuan, Kling, Seedance, and Wan—with five independently generated videos per model, yielding 35 synthetic videos in total. All videos were encoded with 8-bit color depth (0–255 intensity range per RGB channel) and followed a standardized filename structure encoding patient ID, vascular injury site location, model name, and replicate index, enabling automated sorting during analysis (e.g., P1-ICA-Clot-03.mp4). Videos were grouped by patient index (e.g., P1 and P2) for model-level comparisons.

#### Quantitative Evaluation of CLoT Performance

We quantitatively compare CLoT with six commercial video generators (Sora, Wan, Kling, Seedance, Hailuo, Hunyuan) against a “physical-twin” ground truth (fluorescence microscopy). To ensure a fair protocol, all methods are evaluated on the same preprocessed videos (resize to 512×512, light denoising; optional ECC-based stabilization on the green channel). Biology awareness measures use a red-channel platelet proxy (HSV thresholding with morphological opening). Temporal curves are DTW-aligned and resampled to 65 points. We report: pixel/perceptual fidelity, motion fidelity, temporal consistency (platelet coverage vs. time), spatial consistency (occupancy heatmaps), spatiotemporal occupancy (3D space–time cube), and semantic alignment (ViTScore/CLIPScore, linear and DTW). Unless noted, **↑** denotes higher is better; **↓** denotes lower is better. Formal definitions and implementation details for all metrics are provided in table 1.

### Comprehensive comparison

CLoT leads on 18/20 metrics and is second on the remaining two—MAE (marginally behind Sora by ∼0.7%; 0.000271 vs 0.000269) and flow cosine (second to Wan). The radar in Fig. 5b (min–max normalized per metric with lower-is-better metrics inverted) shows the CLoT trace enclosing the largest area, indicating balanced gains across pixel/perceptual quality, motion dynamics, temporal and spatial biology, spatiotemporal occupancy, and semantic alignment.

#### Pixel / perceptual quality (↑)

CLoT attains PSNR/SSIM/MS-SSIM = 27.62 dB / 0.94 / 0.95, improving over Sora by +4.91 dB / +0.319 / +0.329. On LPIPS (↓), CLoT 0.051 represents an ∼82.5% reduction vs. Sora (0.292), consistent with sharper vessel context and platelet textures.

#### Motion dynamics

Using optical-flow-based measures, CLoT achieves EPE (↓) = 0.066 (a 58% reduction vs. Sora’s 0.159) and the highest flow-magnitude correlation (↑) = 0.392. Temporal-difference SSIM (↑) = 0.902 for CLoT is slightly above Wan (0.872). CLoT is second on flow cosine (↑) (CLoT 0.00087 vs. Wan 0.00110).

#### Temporal quality (platelet growth kinetics)

From the red-coverage curves, DTW (↓) is lowest for CLoT (0.000122), ∼94% lower than Sora (0.00207) and far below Wan (0.313). Pearson r (↑) is highest for CLoT (0.610; second best Kling 0.486). On absolute-error metrics, MAE (↓) is 0.000271 for CLoT (second to Sora 0.000269, a ∼0.7% gap), while RMSE (↓) is best for CLoT (0.000548, ∼17% lower than Sora 0.000663).

#### Spatial quality (occupancy heatmaps)

CLoT yields the highest heatmap SSIM (↑) = 0.988 and heatmap cosine (↑) = 0.256, improving over Sora (0.945 and 0.047) by +0.043 SSIM and +0.209 cosine, confirming anatomically plausible platelet placement and density.

#### Spatiotemporal occupancy

On the space–time cube metric, cube cosine (↑) is 0.258 for CLoT vs. 0.033 for Sora and 0.134 for Wan—∼7.9× higher than Sora—reflecting more faithful where–when thrombus evolution.

#### Semantic alignment

With ViT-B/16 and CLIP ViT-B/32 encoders, CLoT tops both linear and DTW variants: ViT linear 0.939 (+0.067 vs. Sora), CLIP linear 0.961 (+0.064), ViT DTW 0.963 (+0.071), CLIP DTW 0.975 (+0.076), indicating superior preservation of global scene semantics and structural cues (vascular geometry, injury context).

#### Biological Fidelity Metric Extraction

All raw videos were processed using a custom MATLAB pipeline (video_data_extraction_v4.m). Videos were first resized (if necessary) to 512×512 pixels using bilinear interpolation. A 360×360-pixel analysis window was then cropped around a user-specified center (default: frame center at X = 256, Y = 256) to focus analysis on the bifurcation region (Analysis ROI). For each frame, red (platelet activation) and green (endothelial cell boundary) channels were separated. Red-channel pixels were classified as valid platelet signal only when red intensity exceeded 10 and was at least 1.5-fold greater than green intensity, filtering non-specific fluorescence and co-localized signals. Three primary quantitative metrics were computed frame-by-frame:

1. Average Platelet Intensity: Mean red-channel intensity within a 140×140-pixel Intensity ROI centered on the injury site, calculated by summing intensities of valid red pixels and dividing by total ROI area. The first frame served as baseline; all subsequent frames were background-subtracted to normalize intensity traces to zero at t = 0.
2. Platelet Area (Analysis ROI): Total count of valid red pixels across the entire 360×360-pixel Analysis ROI, capturing global platelet accumulation.
3. Spatial Overlap Ratio (AI vs. Ground Truth): For each frame at normalized time t, binary platelet masks from AI and ground-truth videos were spatially aligned. The overlap ratio was calculated as:

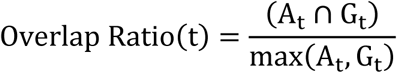

where A_t_ and G_t_ denote valid pixel sets from AI-generated and ground-truth frames at time t, respectively. This formulation normalized for differing thrombus sizes and video lengths. All per-frame metrics were exported to Excel spreadsheets for downstream analysis.

#### Temporal Normalization and Summary Statistics

To enable cross-video comparison despite varying frame counts, time was normalized by converting frame indices to percentage of total video duration, with 100% corresponding to 10 minutes of real-time experiment. A secondary MATLAB script (analyze_timepoint_data_v4.m) extracted metrics at four standardized time points (30%, 50%, 70%, and 90% of video duration) by identifying frames closest to each target percentage. Data were formatted in both tabular spreadsheet format and GraphPad Prism–compatible tables, with separate sheets generated for each metric (Average Platelet Intensity, Platelet Area, and Overlap Ratio). Each AI model was allocated five replicate columns to preserve experiment-level variance for statistical analysis.

#### Spatial Visualization of Platelet Activation Patterns

Representative spatial snapshots were generated using visualize_timepoint_snapshots_v4.m to visualize platelet distribution at key temporal landmarks. For each anatomical category and selected representative video per model, frames at 30%, 50%, 70%, and 90% of video duration were extracted, resized to 512×512 pixels, and cropped to the 360×360-pixel Analysis ROI. Activated-platelet regions were identified using the same validation criteria (red intensity >10 and purity ratio >1.5×green) and rendered as contours with model-specific visual encoding:

- Ground truth: filled black regions (70% opacity)
- CLoT: red wireframe contours (2-pt line thickness)
- Other AI models: colored wireframe contours (1-pt thickness, viridis colormap)

To provide anatomical context, the outer vessel boundary was extracted from the first frame of a reference CLoT video by thresholding the green channel (intensity >10), applying morphological hole-filling to obtain the vessel perimeter, and overlaying the result as a black dash-dot contour. This multi-model overlay enabled direct visual comparison of spatiotemporal platelet activation patterns relative to vessel geometry.

#### Temporal-Trajectory (“Long-Exposure”) Visualization

To depict cumulative spatiotemporal propagation of signals across entire video sequences, all videos were processed using visualize_pixel_trajectories_v2.m. Each video was resized to 512×512 pixels and cropped to a configurable 360×360-pixel Analysis ROI. For each pixel location, the number of frames exhibiting valid signal (green for vessel, red for platelets) was counted and normalized by total frame count, producing a persistence map with values ranging from 0 (never active) to 1 (active in all frames).

Signals were composited onto a white background using alpha-blended rendering:

- Vessel structure (green channel, intensity >60): rendered as gray (RGB: [0.55, 0.55, 0.55]) with α ≤ 0.99
- Platelet activation (red channel, meeting validation criteria): rendered as blue (RGB: [0, 0.2, 1]) with α ≤ 0.90

Darker regions indicated prolonged or repeated signal presence; lighter regions indicated transient activation. This single-image visualization captured global thrombus-propagation dynamics across the entire 10-minute observation period, analogous to long-exposure photography.

#### Individual Intensity-Trajectory Analysis

To quantify temporal activation kinetics at the replicate level, baseline-subtracted Average Platelet Intensity was plotted as continuous time-series traces using plot_intensity_trends.m. For each anatomical category, separate comparison figures were generated pairing each AI model with ground-truth replicates. Time was normalized to percentage of video duration (0–100%, mapped to 0-10min representing experiment time). Ground-truth traces were rendered with 2-pt line thickness; AI-generated traces used 1-pt thickness. This visualization enabled assessment of replicate-to-replicate variability, identification of AI models whose temporal dynamics most closely matched experimental behavior, and detection of systematic deviations in activation kinetics.

#### Video Similarity Quantification

We report 1−NMAE as the similarity score (higher is better). To compare temporal profiles, we first time-normalize each video’s AvgRedIntensity to a 0–100% timeline and select 10 anchor points at 10, 20, …, 100%, yielding a 10-dimensional vector per video. Vectors are then averaged within each cohort (e.g., RC-ICA-Clot-ABC, RC-ICA-GT-ABC, RC-ICA-Clot-Tica, and RC-ICA-GT-Tica to obtain cohort-level 10-point profiles. Given two cohort profiles *x*, *y* ∈ *R*^10^, we compute:

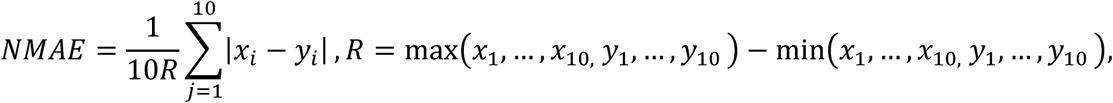

(using *R*=1 if values are already in [0,1]), and report

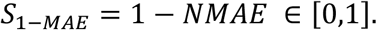

Thus, *S*_1−MAE_ = 1 indicates identical cohort trajectories, while lower values reflect larger average absolute deviations, expressed as a fraction of the joint dynamic range. We evaluate the similarity between the ground truth videos and Clot generated videos under two antiplatelet treatments i.e., Abc and Tica (84% and 96.6%,), achieving a similarity score *S*_1−*MAE*_ of 90.3%.

#### Platelet Detachment Event Quantification

Clot detachment events—indicative of embolic risk—were manually enumerated from all videos. Detachment was defined as an abrupt, sustained decrease in red signal intensity following a period of accumulation (increasing red area). Each detachment event represented a discrete embolus release and was tabulated per video for subsequent statistical comparison with AI-generated predictions.

### Statistical Analysis

#### Sample size and experimental design

For physical-twin blood perfusion experiments, a sample size of n ≥ 3 replicates per condition. Experiments were performed on vessel chips from 3 distinct patients with 10-20 conditions per patient, yielding approximately 500 total experiments.

#### Statistical tests and multiple comparisons

Platelet-adhesion metrics (fluorescence intensity) and kinetic parameters (embolization frequency) were compared across conditions via two-way ANOVA (factors: injury site, drug treatment; repeated measures across patients). Statistical significance is indicated by * = p < 0.05, ** = p < 0.01, ** = p < 0.0001 assessed by unpaired, two-tailed Student’s t-test.

## Acknowledgment

We thank Freda Passam for clinical guidance and support. We thank Peter Su, Yiyao Chen, Yinyan Wang for providing critical suggestions. This work was supported by MRFF Cardiovascular Health Mission Grants (MRF2016165– L.A.J. and Z.L.; MRF2023977 – L.A.J., F.P., Z.L., K.S.B., and T.A.); MRFF Early to Mid Career Researchers Grant (MRF2028865 – L.A.J. and Z-H.W.; MRF2037779 – L.A.J.); Lining Arnold Ju is a National Heart Foundation Future Leader Fellow Level 2 (105863) and a Snow Medical Research Foundation Fellow (2022SF176). Yunduo Charles Zhao is a National Heart Foundation PhD Scholar (106879) and NHMRC PhD Scholar (GNT2022247) All experiments were performed in accordance with relevant guidelines and approved by the University of Sydney Human Research Ethics Committee. Informed consent were obtained from human participants of this study. All data that support the findings of this study are available on request from the corresponding author.

## Author contributions

These authors contributed equally to this work: Z.W, Y.C.Z, and H.Z. L.A.J. and Y.C.Z designed the research, analyzed data and wrote the manuscript. Z.W designed the chip, developed the automated manufacturing system, made the chips, wrote the manuscript, and performed data analysis. Y.C.Z co-designed the chip, performed blood perfusion, conducted analysis and interpretation of data; Y.L, A.N and N.A.Y helped culture the endothelial cells and vascularized the chips. Y.L and A.N performed immunostaining of the vascularized chips; and N.A.Y. performed data analysis and wrote the manuscript; K.S.B provided clinical expertise, co-wrote and revised the paper. T.A. provided the CTA images and contributed to patient data analysis. L.A.J is the senior and corresponding author.

## Data and materials availability

All data are available in the main text or the supplementary materials.

---

**Figure S1.**
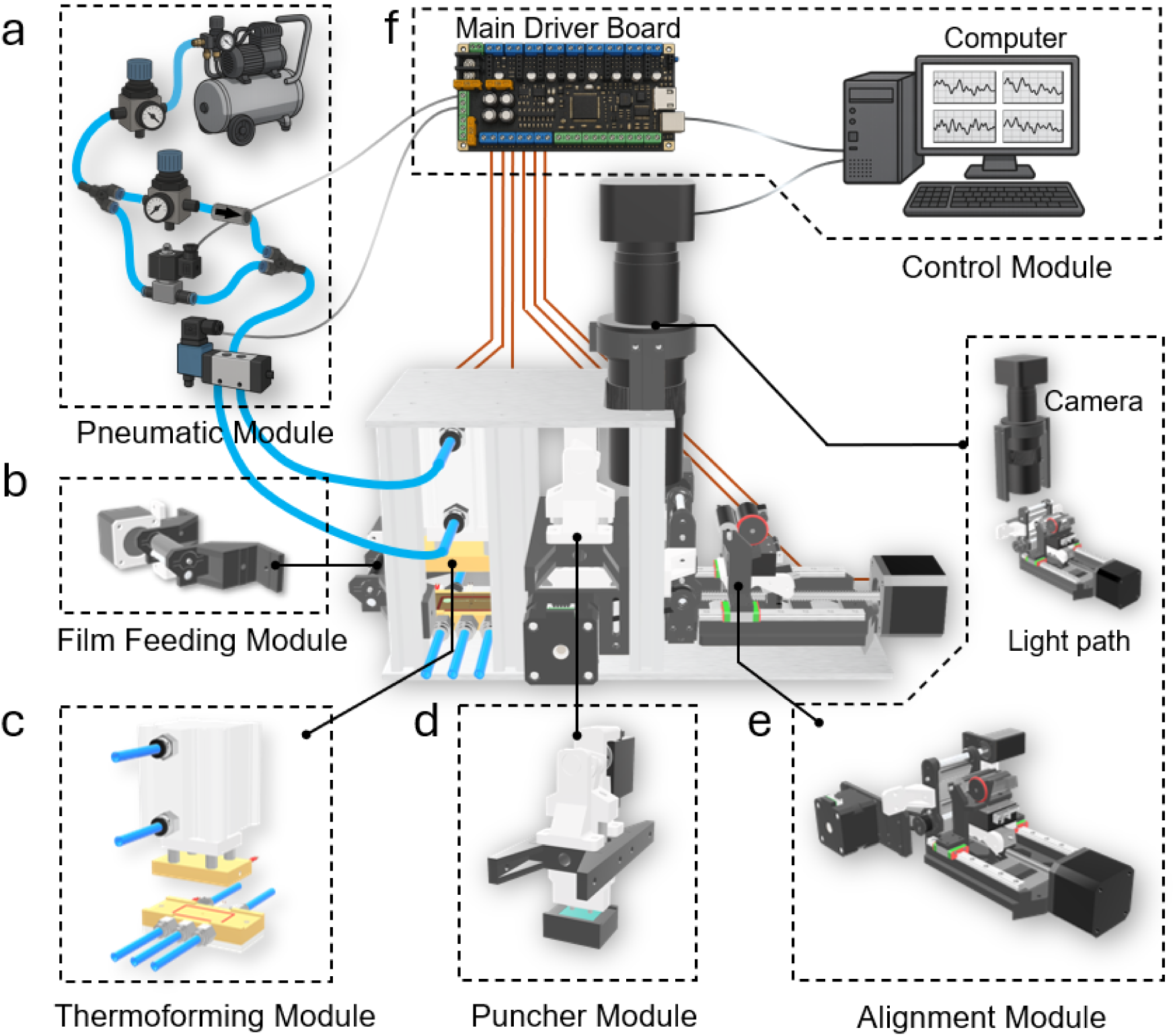
Architecture of the automated vessel-on-a-chip production system. The complete system integrates six functional modules enabling continuous, high-throughput fabrication. (**A**) Pneumatic module supplying independently regulated low-pressure (∼1 bar, soft-start/stop) and high-pressure (∼5 bar, high-force molding) air streams. (**B**) Film-feeding module with active motor-driven roller and compliant passive roller, maintaining consistent SEBS film tension during feeding. (**C**) Thermoforming module with dual independently controlled heating blocks (top: 2 cartridges, 80 W total; bottom: 1 cartridge, 40 W) achieving rapid thermal ramp-up (∼40 °C/min) and precision temperature control (±3 °C) via PID feedback and integrated K-type thermocouples. (**D**) Puncher module with servo-motor-driven crankshaft linkage converting rotary motion to linear reciprocating punch displacement, creating perfusion inlets/outlets with controlled penetration depth. (**E**) Alignment module integrating five motorized axes (folding, horizontal, dual tension-roller, feeding-roller) for real-time optical feedback-guided chip assembly. (**F**) Control module comprising main driver board (Azteeg X3 Pro), temperature controllers (PID), pneumatic driver relays, and MATLAB-based host interface coordinating all subsystems.

**Figure S2.**
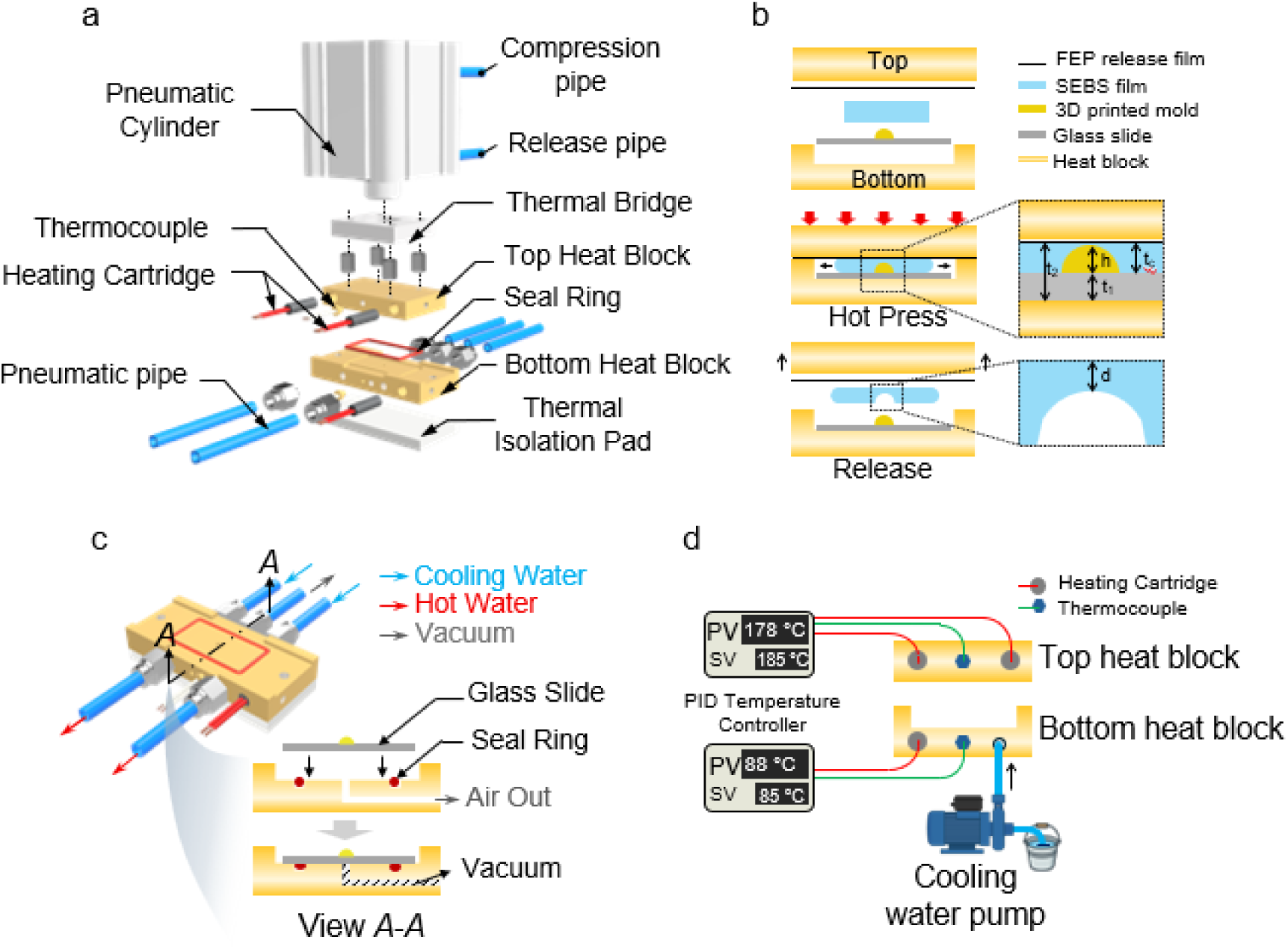
Structure and thermal operation of the thermoforming module. (**A**) Exploded schematic of the thermoforming module structure showing the pneumatic cylinder, dual heating cartridges in the top block, single heating cartridge in bottom block, thermocouples, thermal bridges (3D-printed polymer), and thermal-isolation pad separating the bottom block from the structural frame. (**B**) Schematic of the hot-press process. A multilayer sandwich (FEP release film, SEBS film, 3D-printed mold on glass slide) is positioned between heat blocks. During hot-pressing, the softened SEBS is deformed into the mold cavity. Final chip thickness (*t*_c_) is determined by bottom block cavity depth (*t*_2_) and glass-slide thickness (*t*_1_): *t*_c_ = *t*_2_ – *t*_1_. Minimum distance between channel and chip surface (*d*) affects imaging quality: *d* = *t*_c_ – *h*, where h is mold height. (**C**) Cross-sectional view showing integrated water cooling (circulating cooling water at room temperature ∼26 °C through dual internal channels) and vacuum-locking system (sealed cavity with air evacuation path enabling vacuum locks the glass slide against peeling forces during film demolding). (**D**) PID temperature-control circuitry with independent channels for top and bottom heat blocks, enabling precise ±3 °C unloaded thermal regulation.

**Figure S3.**
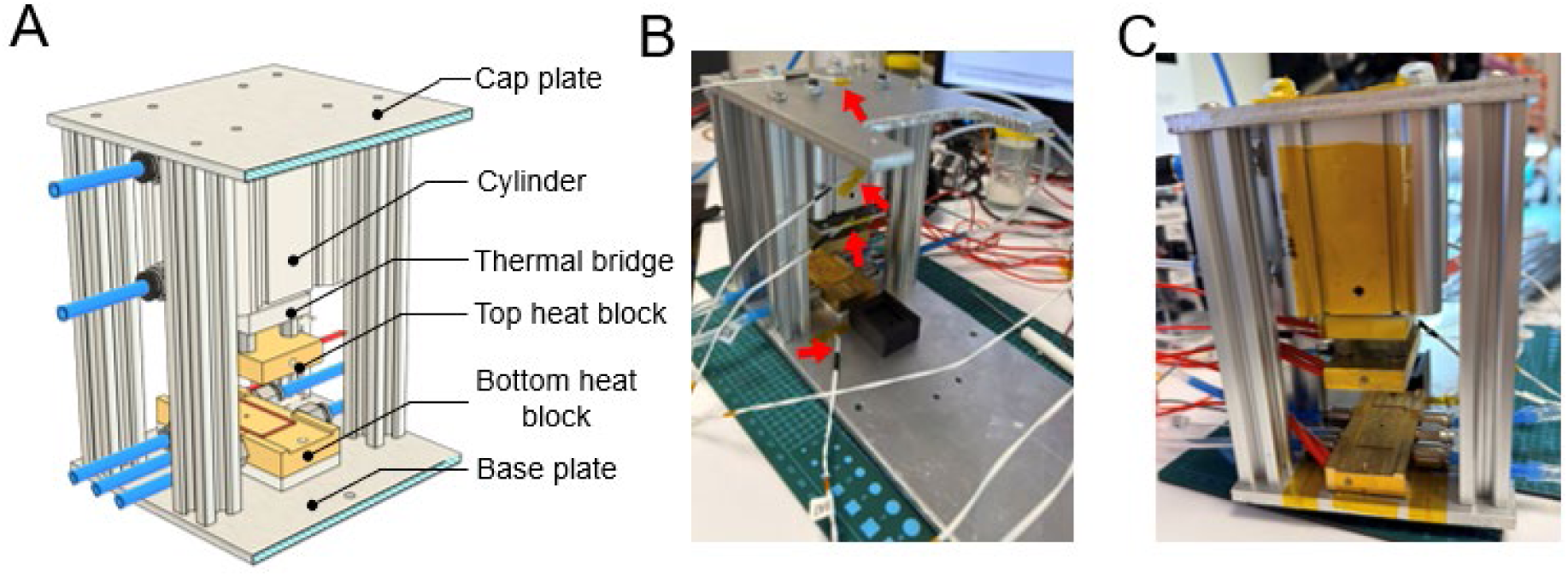
Structural terminology and thermographic preparation of the thermoforming module. (**A**) Schematic showing major structural components: cap plate, pneumatic cylinder, thermal bridge, top/bottom heat blocks, base plate, frame. These terms correspond to temperature-distribution plots in subsequent figures. (**B**) Photograph indicating thermistor positions (red arrows) for monitoring temperature at key structural points: top heat block, bottom heat block, cylinder, thermal bridge, cap plate, base plate. (**C**) All exposed metal surfaces are covered with Kapton tape (emissivity ε ≈ 0.76) to standardize emissivity and enable accurate infrared thermographic imaging.

**Figure S4.**
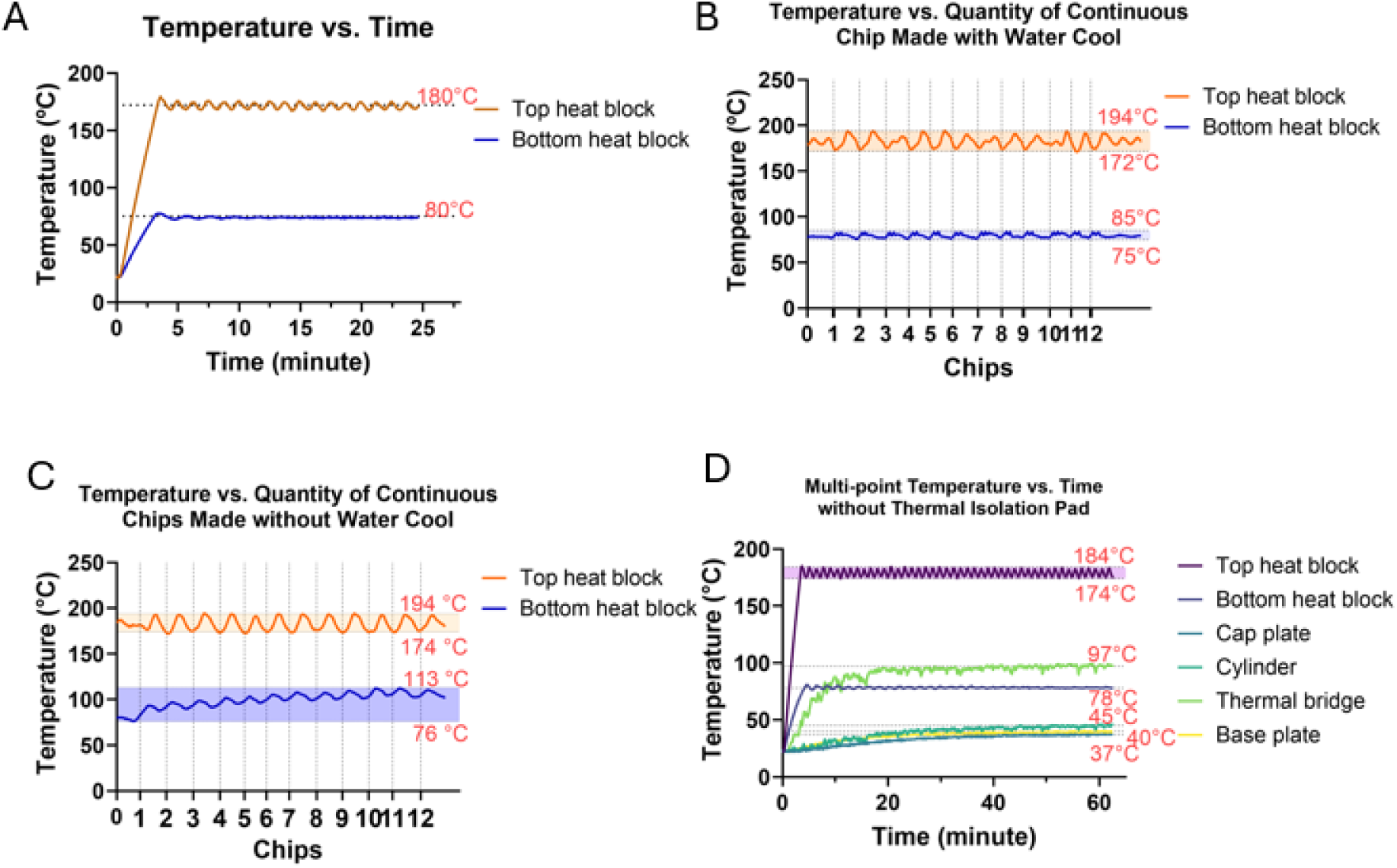
Temperature profiles of the thermoforming module under different operating conditions. (**A**) Temperature–time curve after system power activation showing rapid ramp to setpoint temperatures: top block 180 °C, bottom block 80 °C, with stability achieved within ∼4 min. (**B**) Multi-point temperature monitoring (top/bottom blocks, cap plate, cylinder, thermal bridge, base plate) for module without thermal isolation pad, showing distinct thermal gradients and elevated parasitic heating in non-target regions. (**C**) Temperature stability of top and bottom blocks during continuous high-throughput production with active water cooling applied. Bottom block maintained within 75–85 °C range across 10+ consecutive molding cycles, demonstrating stable processing. (**D**) Temperature evolution under identical continuous production conditions but without active water cooling. Bottom block temperature rises progressively, reaching ∼113 °C after 12 cycles, indicating significant thermal accumulation and need for active cooling during extended production runs.

**Figure S5.**
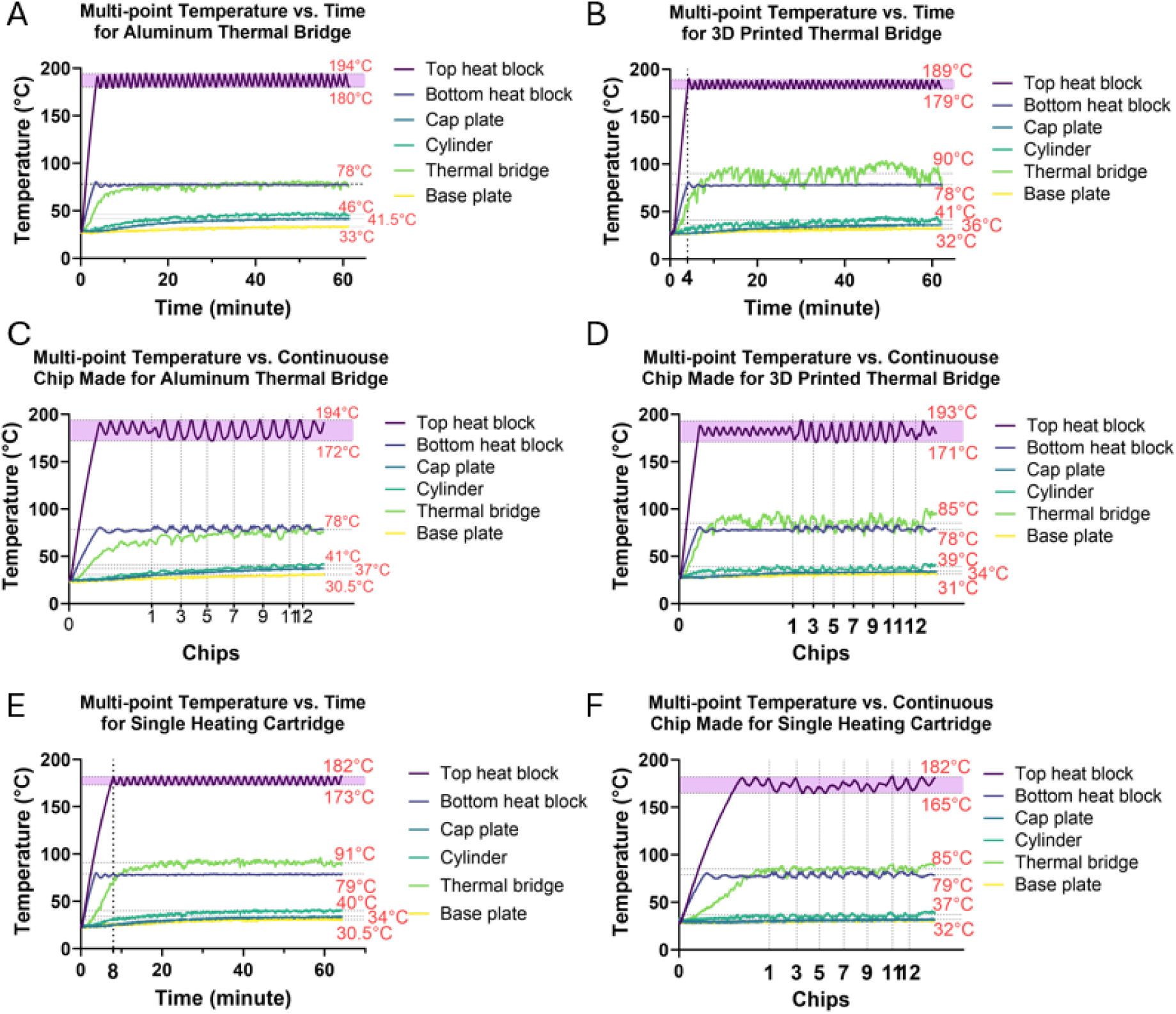
Multi-point temperature measurements of the thermoforming module under different configurations. Temperature profiles recorded at six structural locations (top/bottom blocks, cap plate, cylinder, thermal bridge, base plate) under varied configurations. (**A-B**) Aluminum thermal bridge vs. 3D-printed thermal bridge: aluminum exhibits lower bridge temperature but elevated cylinder/cap plate temperatures (∼5 °C higher), confirming poor thermal isolation despite lower bridge temperature. (**C-D**) Temperature evolution during continuous multi-cycle chip production for both bridge types, showing sustained lower cylinder/cap-plate temperatures with the 3D-printed bridge. (**E-F**) Single-heating-cartridge configuration (top block only) showing ∼2× slower thermal ramp (reaching setpoint in ∼8 min vs. ∼4 min with dual-cartridge design) and irregular temperature fluctuations, reducing thermal stability during repeated production cycles compared to the standard dual-cartridge configuration.

**Figure S6.**
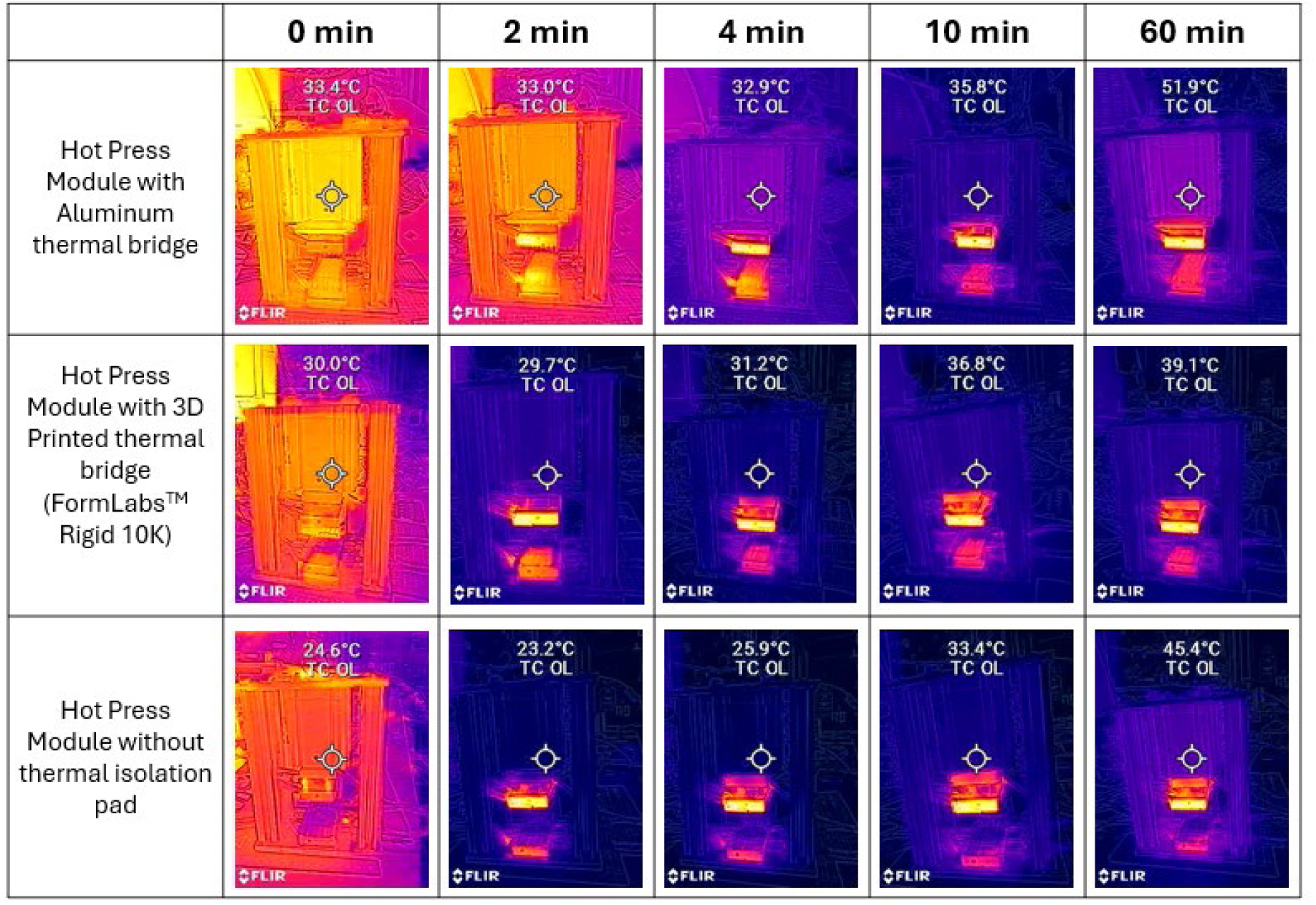
Thermographic comparison of the thermoforming module under different configurations. Thermal infrared images captured at *t* = 0, 2, 4, 10, and 60 min showing evolution of temperature distribution. Three configurations compared: (*top row*) Aluminium thermal bridge (high thermal conductivity) → progressive temperature increase in the cylinder region from 33.4 °C (*t*=0) to 51.9 °C (*t*=60 min) due to parasitic heat conduction. (*middle row*) 3D-printed polymer thermal bridge (FormLabs™ Rigid 10K resin; low thermal conductivity) → reduced cylinder temperature (39.1 °C at t=60 min) confirming superior thermal isolation. (*bottom row*) Module without thermal-isolation pad under bottom block → elevated frame and cylinder temperatures (45.4 °C at t=60 min) indicating heat loss through structural conduction. Temperature scale in images indicates measured temperature at cylinder central point.

**Figure S7.**
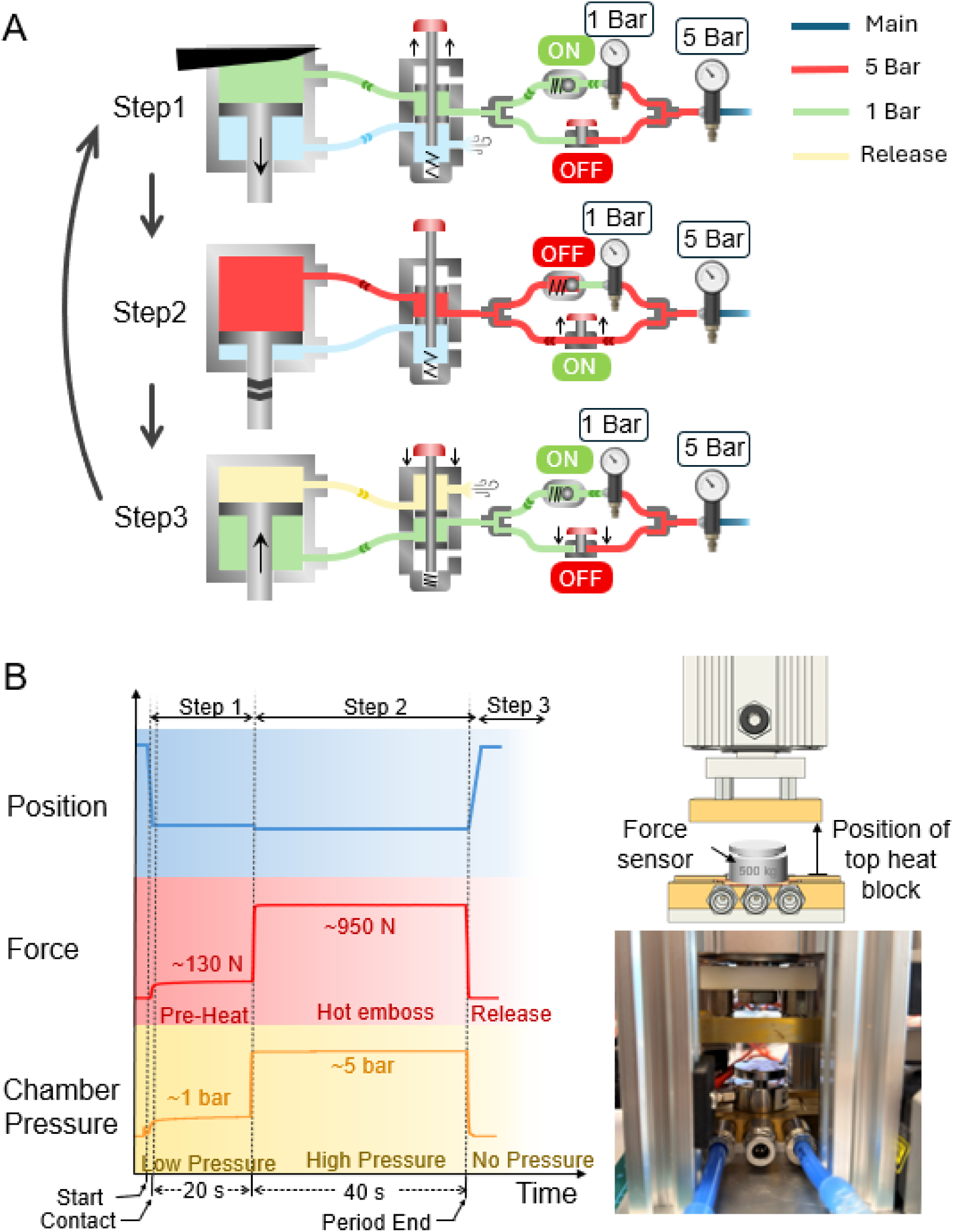
Working principle of the pneumatic and thermoforming modules. (**A**) Schematic of the dual-stage pneumatic control system. Main regulator reduces primary air supply to 5 bar; secondary regulator further reduces one branch to ∼1 bar (low-pressure line through check valve). The other branch retains 5 bar (high-pressure line controlled by shut-off valve). Both branches recombine at a Y-junction, feeding a 5/2-way solenoid valve connected to the pneumatic cylinder. This architecture enables smooth soft-start/stop motion protecting the glass mold from mechanical shock. (**B**) Temporal profile of the three-phase thermoforming cycle: Step 1 (pre-heating, ∼20 s), low-pressure air (∼1 bar, ∼130 N force) gently lowers the top heat block, heating the SEBS film above Tg; Step 2 (hot-press, ∼40 s), high-pressure air (∼5 bar, ∼950 N force) fully presses the molten SEBS into mold cavities; Step 3 (release and cooling, ∼10 s), high-pressure air is exhausted and low-pressure air retracts the top block smoothly, allowing SEBS to cooldown. Applied force is calibrated via load cell. Corresponding pressure evolution in the pneumatic cylinder extension chamber is logged, showing transition from ∼1 bar (Step 1) to ∼5 bar (Step 2) to atmospheric pressure (Step 3).

**Figure S8.**
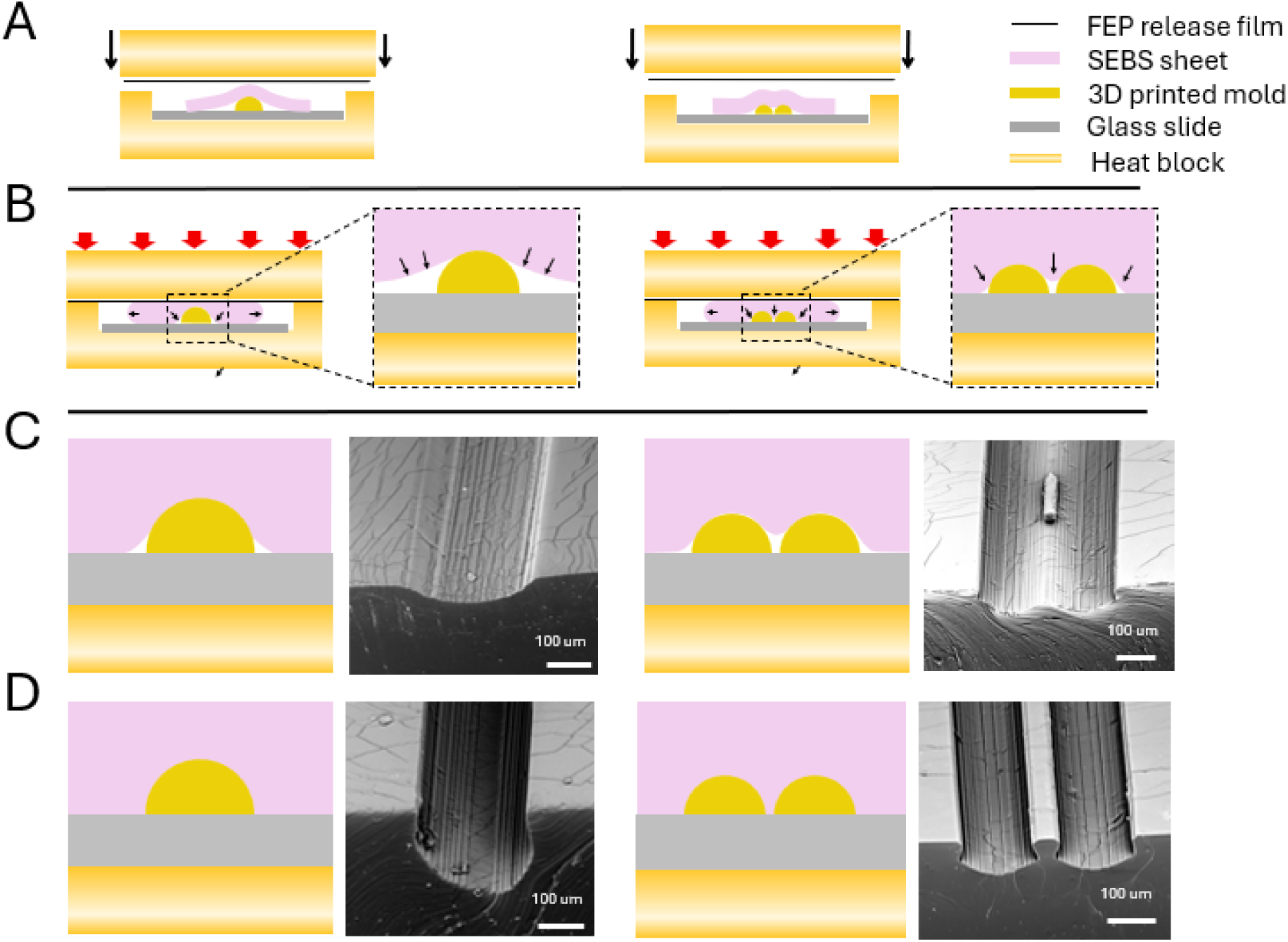
Representative failure modes in the thermoforming process. (**A**) Schematic of representative mold designs: single circular channel and closely-adjacent dual channels. (**B**) Failure mechanism schematics illustrating effects of temperature and film thickness. Left: insufficient heating → rigid SEBS fails to conform to mold, producing rounded edges and reduced forming depth. Right: thin film + narrow mold gaps → insufficient material flow into confined spaces. (**C-D**) Corresponding cross-sectional schematics and SEM micrographs. (**C**) *Left*: incomplete mold filling due to low temperature; Right: thin SEBS film fails to fill narrow inter-mold gaps. (**D**) Left: optimal heating enables sharp corner definition; Right: adequate film thickness ensures complete gap filling. Scale bars: 100 μm.

**Figure S9.**
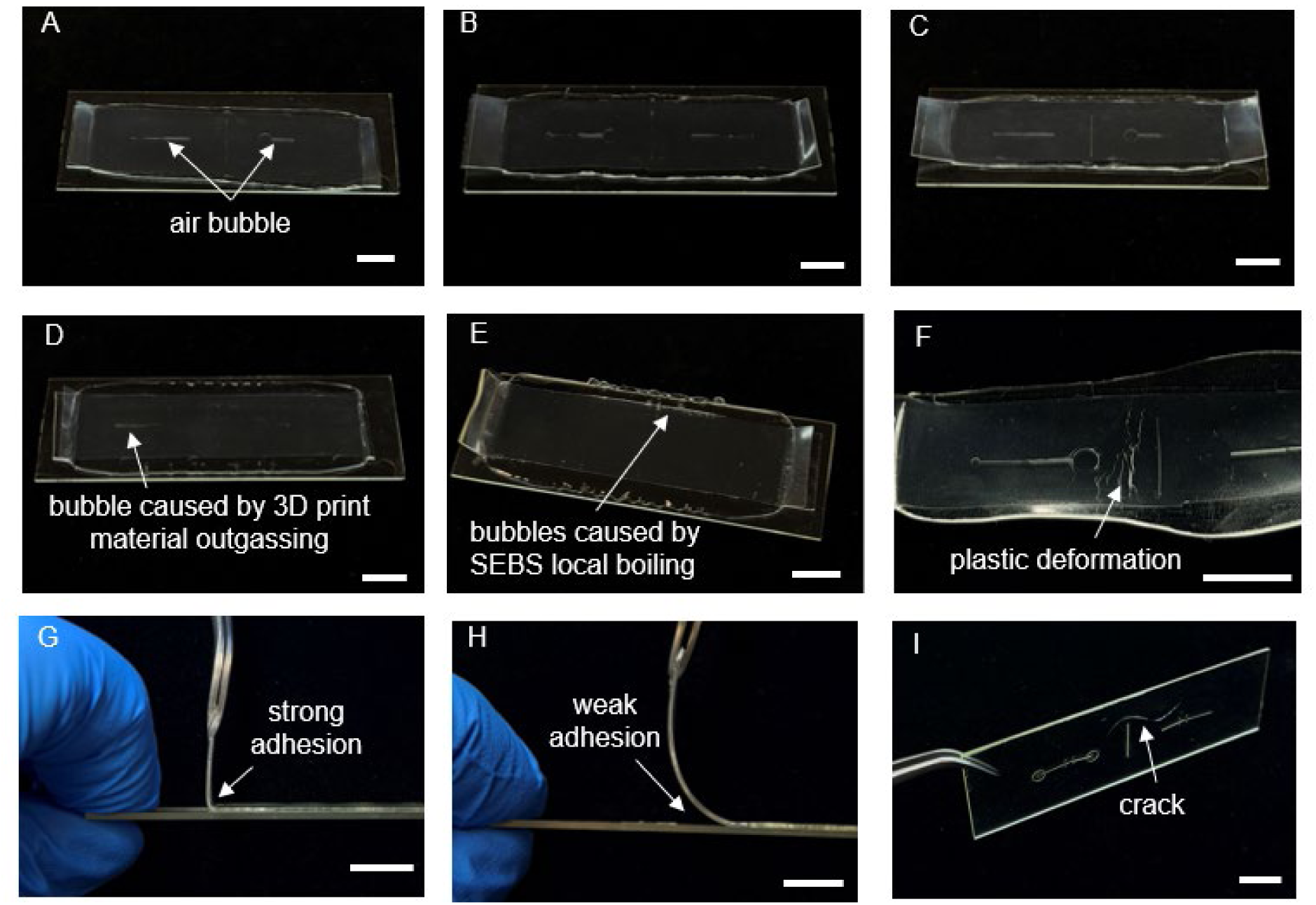
Representative failure modes in the thermoforming process. Photographs demonstrating various failure outcomes during hot-press thermoforming under suboptimal conditions. (**A**) Insufficient top-block temperature: molten SEBS fails to fully conform to mold, leaving unfilled corners and visible trapped air bubbles (arrow). (**B**) Insufficient pressing duration: SEBS lacks time for material flow, producing similar incomplete corner filling. (**C**) Bottom-block temperature insufficient: glass mold remains inadequately heated, preventing proper SEBS material flow and corner definition (rounded, unfilled corners visible). (**D-E**) Excessive top-block temperature (left and center) or prolonged heating: SEBS undergoes local boiling, generating bubbles visible as dark voids in the molded channel (arrows). Outgassing from the 3D-printed mold contributes additional bubbling and surface defects. (**F**) Film degradation at excessive temperature: SEBS remains excessively plastic after molding, deforming under minimal demold forces and sustaining permanent structural damage. (g–h) Adhesion behavior with glass mold. (**G**) Premature cooling before demolding: film adheres strongly to glass (visible adhesion at interface, arrow), risking damage to both film and mold during separation. (**H**) Optimal temperature during demolding: film detaches smoothly without structural distortion, yielding undamaged chip and reusable mold. (**I**) Cracked mold glass: results from absence of two-stage pneumatic motion; without soft-start/stop control, rapid high-pressure air influx drives top block downward abruptly, mechanically impacting and fracturing the brittle glass mold (crack visible, arrow). All scale bars: 10 mm

**Figure S10.**
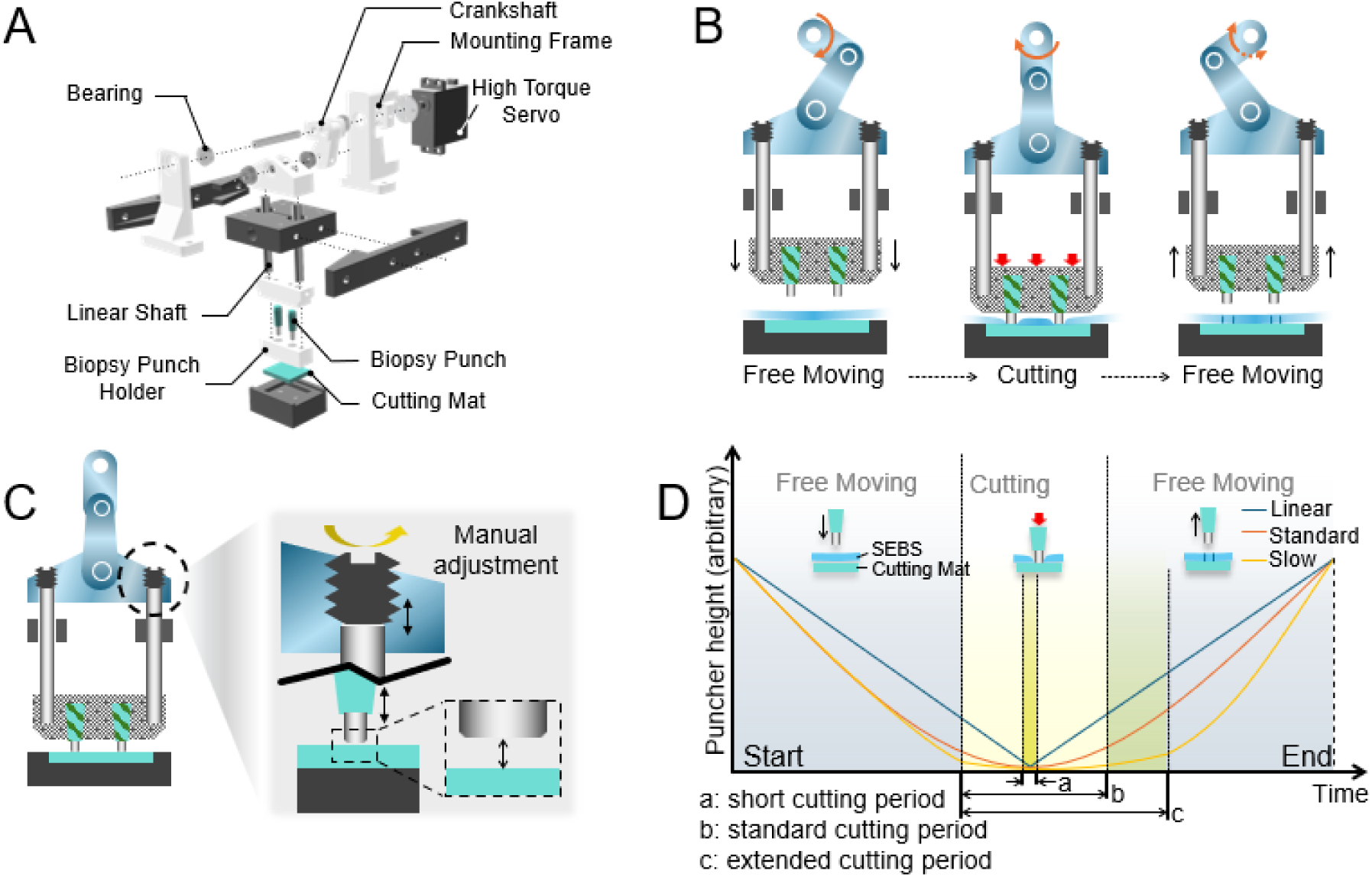
Structure and motion configuration of the puncher module. (**A**) Exploded schematic showing the high-torque servo motor (HB7545GS; peak torque 450 N·cm; speed 91 rpm), custom-designed 3D-printed crankshaft mechanism (Formlabs Form 3+), dual linear guide rails (MGN7C HIWIN), and biopsy punch assembly (KAI Medical) with interchangeable punch heads (diameters: 3.0 mm) and modular cutting mat. (**B**) Illustration of the punch motion cycle: downward stroke penetrates the SEBS film to controlled depth, leaving punched material partially attached; upward stroke completes the cycle. This prevents debris accumulation and eliminates need for continuous material removal. (**C**) Adjustable screw mechanism for fine-tuning punch penetration depth (adjustment range: ±5 mm; resolution: 10 μm) accommodating SEBS films of varying thickness (150–250 μm). (**D**) Configurable cutting-period length (time interval of punch engagement) achieved by adjusting servo motor speed and phase angle, enabling optimization for different film thicknesses and material stiffness.

**Figure S11.**
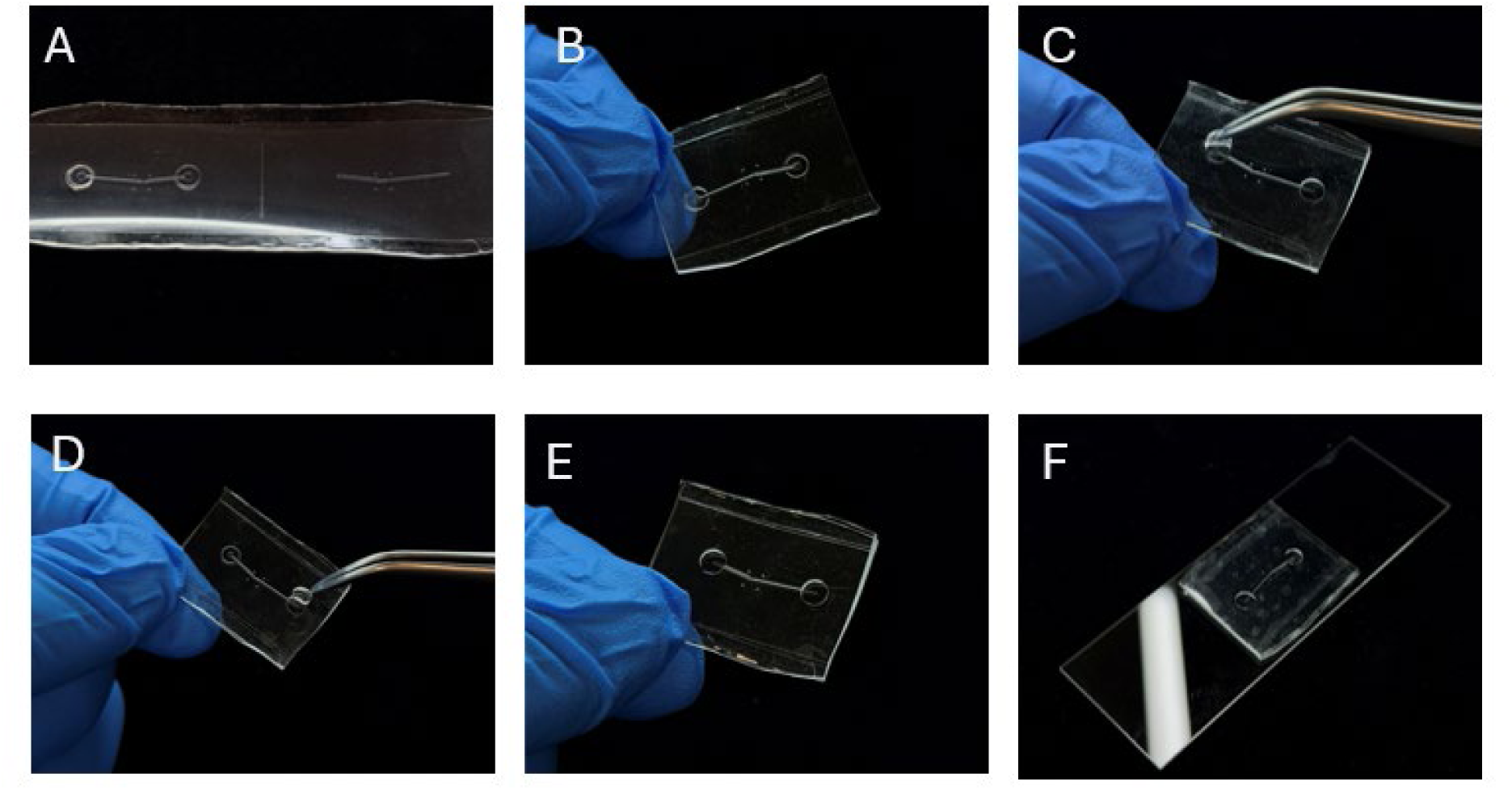
Punching and preparation of perfusion inlets and outlets in the vessel-on-a-chip. Sequence documenting the punch process and inlet/outlet preparation. (**A**) SEBS film immediately after partial punching, showing circular dimples with material partially attached. (**B**) Aligned vessel-on-a-chip with pre-cut inlet/outlet regions. (**C-D**) Punched material is gently lifted and removed using fine-tip tweezers after alignment, revealing clean circular openings. (**E**) Completed chip showing fully exposed perfusion ports (diameter: 3.0 mm). (**F**) Final vessel-on-a-chip mounted on glass slide substrate, ready for endothelial seeding and downstream perfusion experiments. Spacing between inlets/outlets adjusted to match cell seeding and perfusion experiment setup; typical spacing: 10–15 mm.

**Figure S12.**
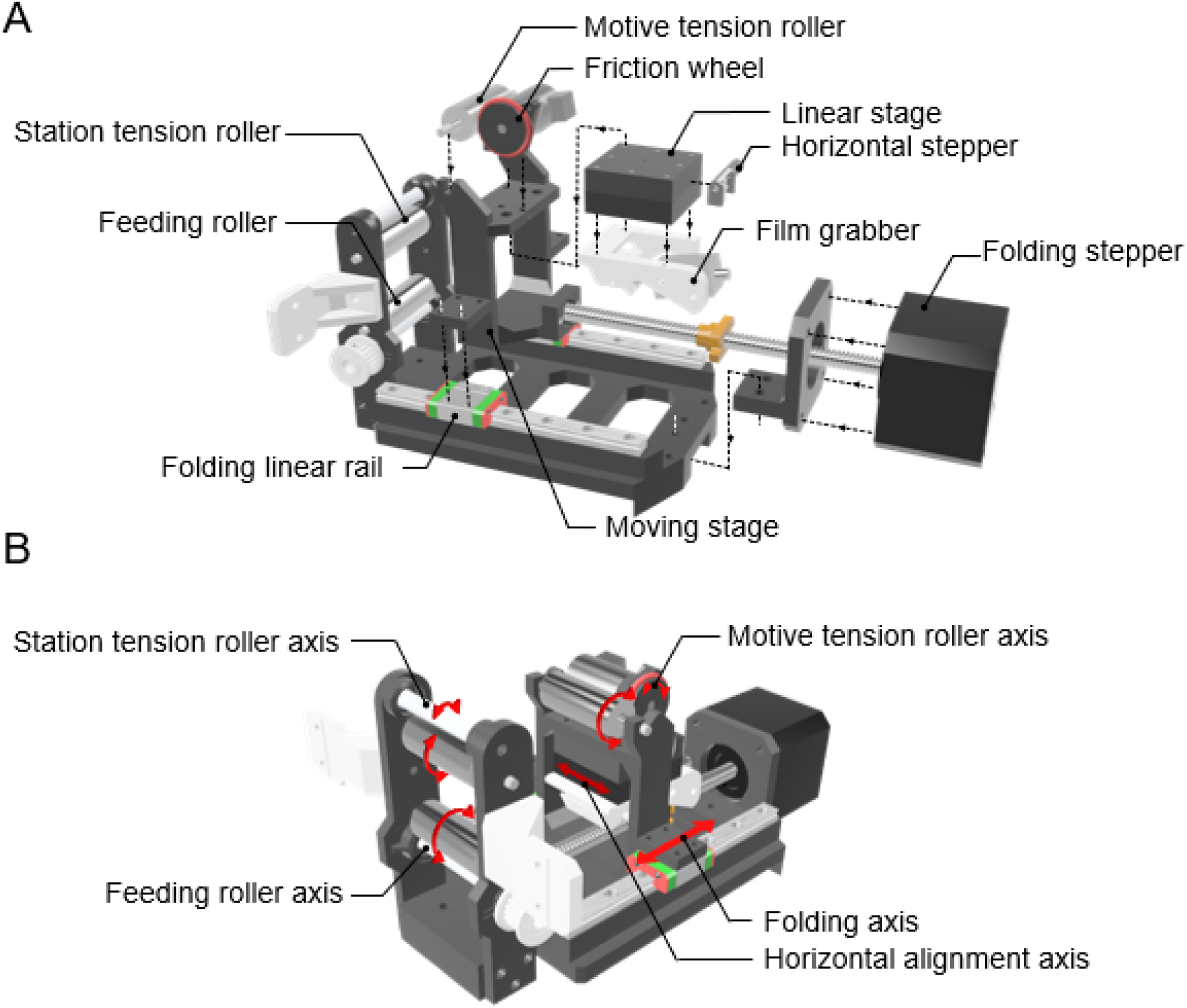
**Structure and motion configuration of the alignment module**. (**A**) Exploded schematic showing integration of five motorized axes: feeding roller (stepper motor + timing-belt drive), station and motive tension rollers (dual stepper motors), film grabber (solenoid-actuated 3D-printed clamping mechanism), moving stage (lead-screw-driven folding axis), and horizontal stage (stepper-motor-driven lateral translation on precision rails). (**B**) Motion-axis configuration diagram showing the folding axis (vertical z-motion controlling film folding height and compression), dual tension-roller axes (adjustable stretching force), feeding-roller axis (synchronized feeding), and horizontal alignment axis (x-direction lateral registration). Coordinated motion of these axes positions and aligns the two vessel halves before bonding.

**Figure S13.**
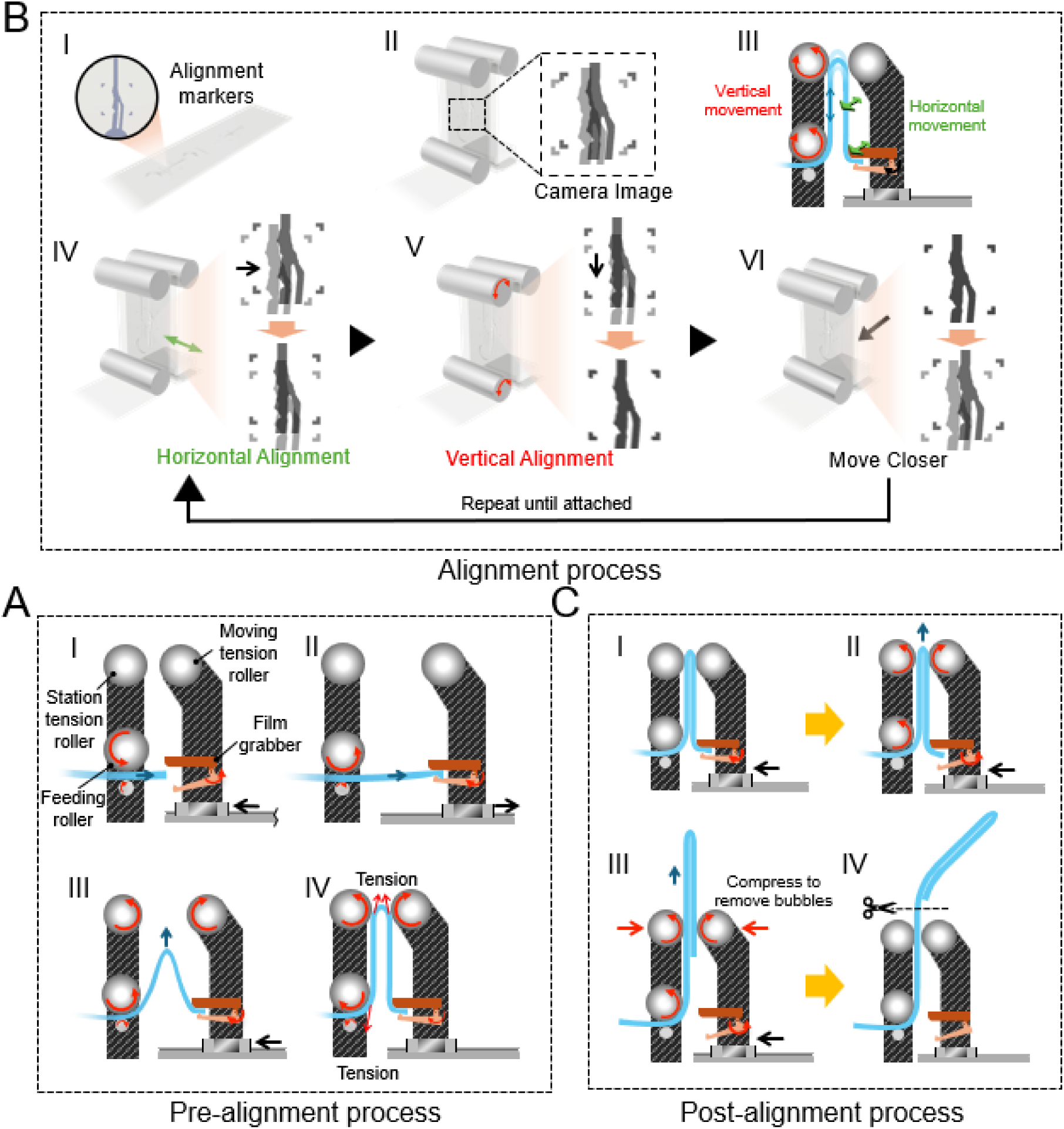
**Detailed sequence of the automated alignment process**. (**A**) Film-feeding module advances SEBS film; film grabber opens and secures the film (clamping force ∼0.85 N); grabber moves outward synchronized with feeding roller to prevent slack; film is folded by retracting stage while feeding continues, causing the film to arch upward; folded section contacts tension rollers which rotate to apply upward tensioning force stabilizing the film. (**B**) OpenCV-based vision system captures both top and bottom channel halves via reflection imaging, identifying four alignment fiducial markers per half-lumen. Misalignment offsets (Δx, Δy, Δθ) are computed; closed-loop control commands adjustments via station/motive tension rollers (z-axis) and horizontal stage (x-axis). Multiple feedback cycles (typically 3–5) progressively reduce misalignment error to <±10 μm. (**C**) Film grabber releases; tension/feeding rollers advance the aligned chip upward; folding stage applies compression force (∼ 30 N) removing air bubbles and ensuring uniform bonding; completed chip is manually separated by user, residual film retracts automatically resetting the system.

**Figure S14.**
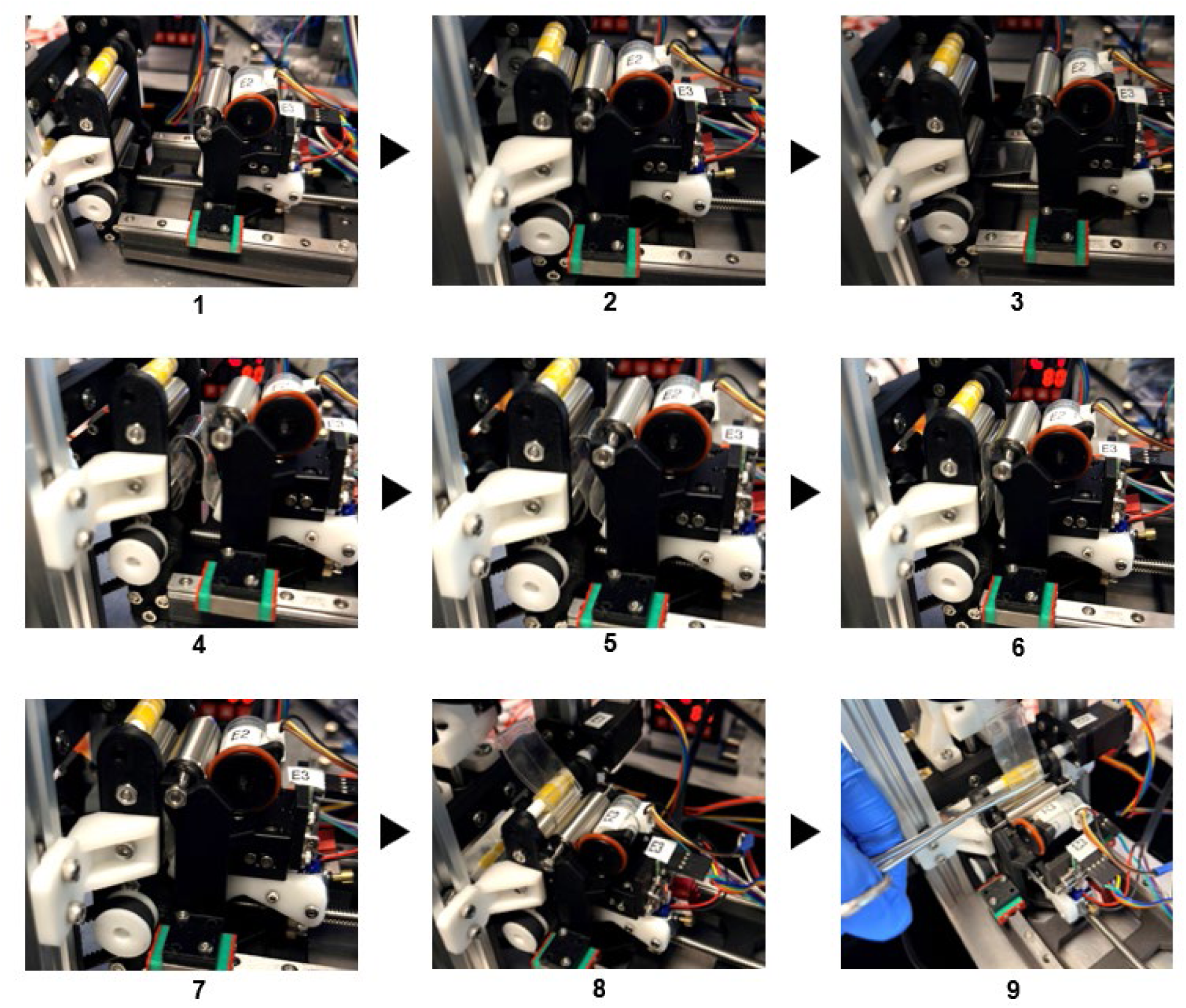
Sequential steps of a single vessel-chip alignment cycle. Nine photographs documenting the complete alignment cycle: (**1**) system ready for new cycle; (**2**) SEBS film grasped by film grabber; (**3**) film pulled outward initiating alignment; (**4**) film folded and repositioned for controlled advancement; (**5**) film extruded until front edge reaches tension roller; (**6**) film engaged and stretched by tension roller for stabilization; (**7**) full-lumen vessel-on-chip structure accurately aligned (alignment tolerance <±10 μm); (**8**) aligned chip compressed and guided through roller pair for uniform delivery; (**9**) aligned chip separated and retrieved, film retracts automatically. Total cycle time: ∼30 seconds.

**Figure S19.**
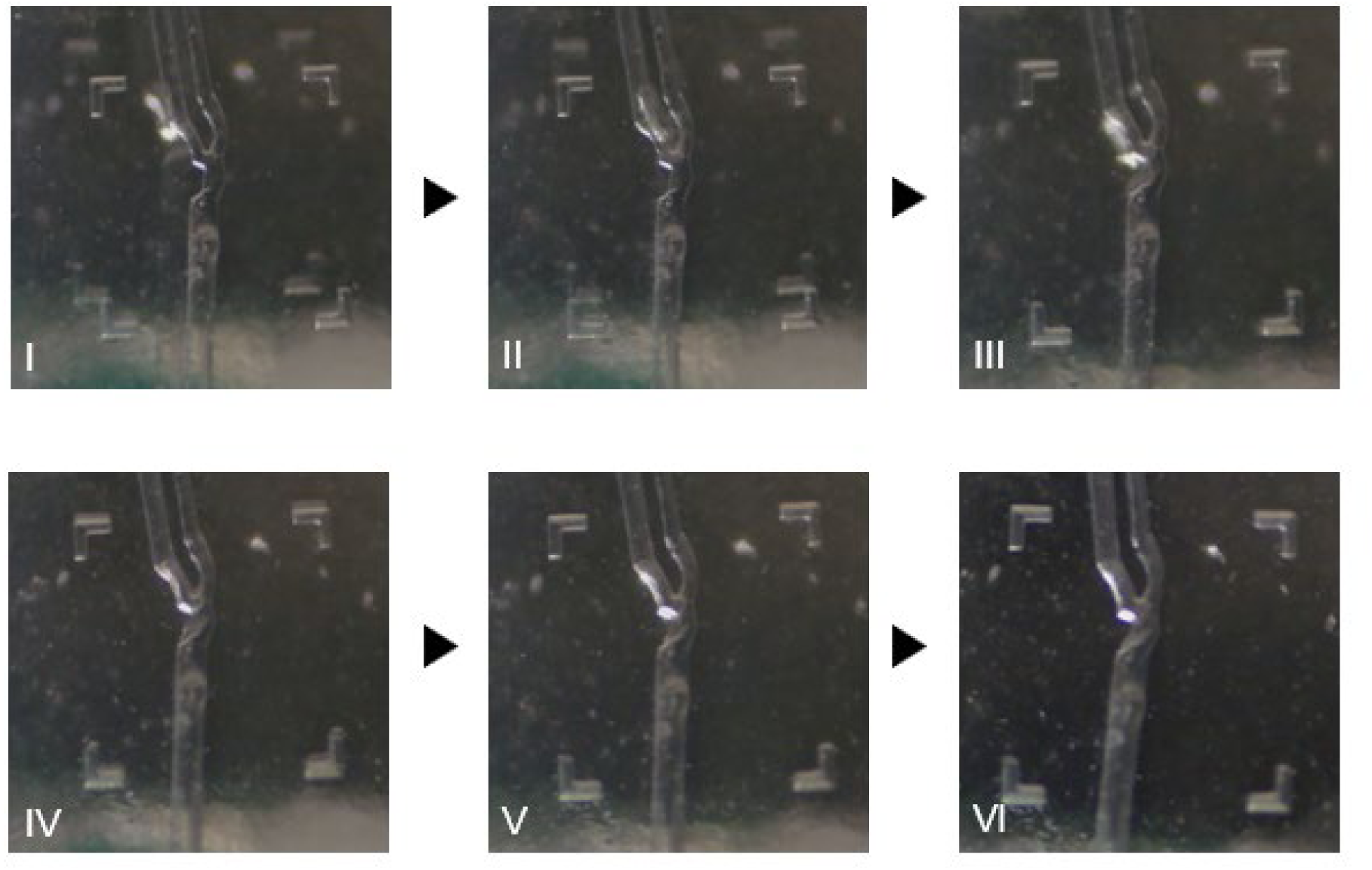
Assessment of the impact of antithrombotic agents for localised thrombosis for patient 2. (**A**) Anticoagulant effects (Group 1): Time-lapse fluorescence intensity of platelets in the presence of rivaroxaban and apixaban compared to control, showing nonsignificant changes in platelet accumulation. (**B**) Quantification of platelet translocation (Group 1): Violin plot showing the number of platelet translocation events under control and anticoagulant conditions. No significant differences are observed between the control condition and those with rivaroxaban or apixaban. (**C**) VWF-GPIb inhibition (Group 2): Time-lapse fluorescence intensity of platelets in the presence of AK2 and ARC1172 compared to control, demonstrating reduced platelet accumulation. (**D**) Quantification of platelet translocation (Group 2): Violin plot indicating a significant reduction in platelet translocation events with AK2 and ARC1172 compared to control. (**E**) Integrin αIIbβ3 inhibition (Group 3): Time-lapse fluorescence intensity of platelets in the presence of abciximab and tirofiban compared to control, showing slight inhibition of platelet aggregation. (**F**) Quantification of platelet translocation (Group 3): Violin plot showing a significant reduction in platelet translocation events with abciximab and tirofiban compared to control. (**G**) TXA_2_ and P2Y_12_ inhibition (Group 4): Time-lapse fluorescence intensity of platelet adhesion in the presence of aspirin and ticagrelor compared to control, indicating reduced platelet accumulation with ticagrelor but not aspirin. (**H**) Quantification of platelet translocation (Group 4): Violin plot showing no significant difference in platelet translocation events between aspirin and the control condition, while ticagrelor significantly reduces translocation events. Statistical significance is indicated by * = p < 0.05, ** = p < 0.01, assessed by unpaired, two-tailed Student’s t-test.

**Figure S20.**
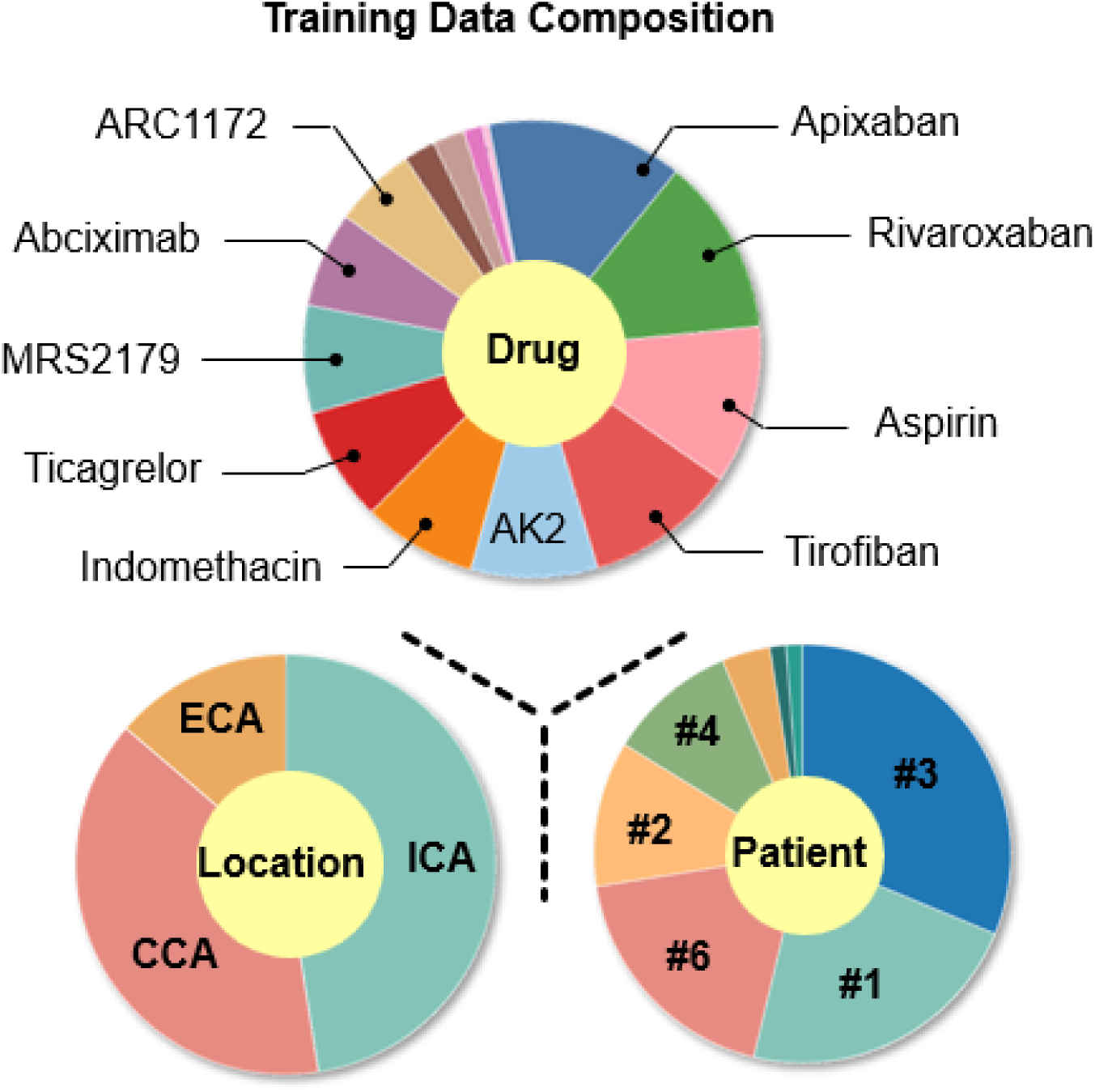
Training data composition. Distribution of training samples across drug types (top), vessel locations—external carotid (ECA), internal carotid (ICA), and common carotid (CCA) (bottom left)—and patient geometries (bottom right).

**Figure S21.**
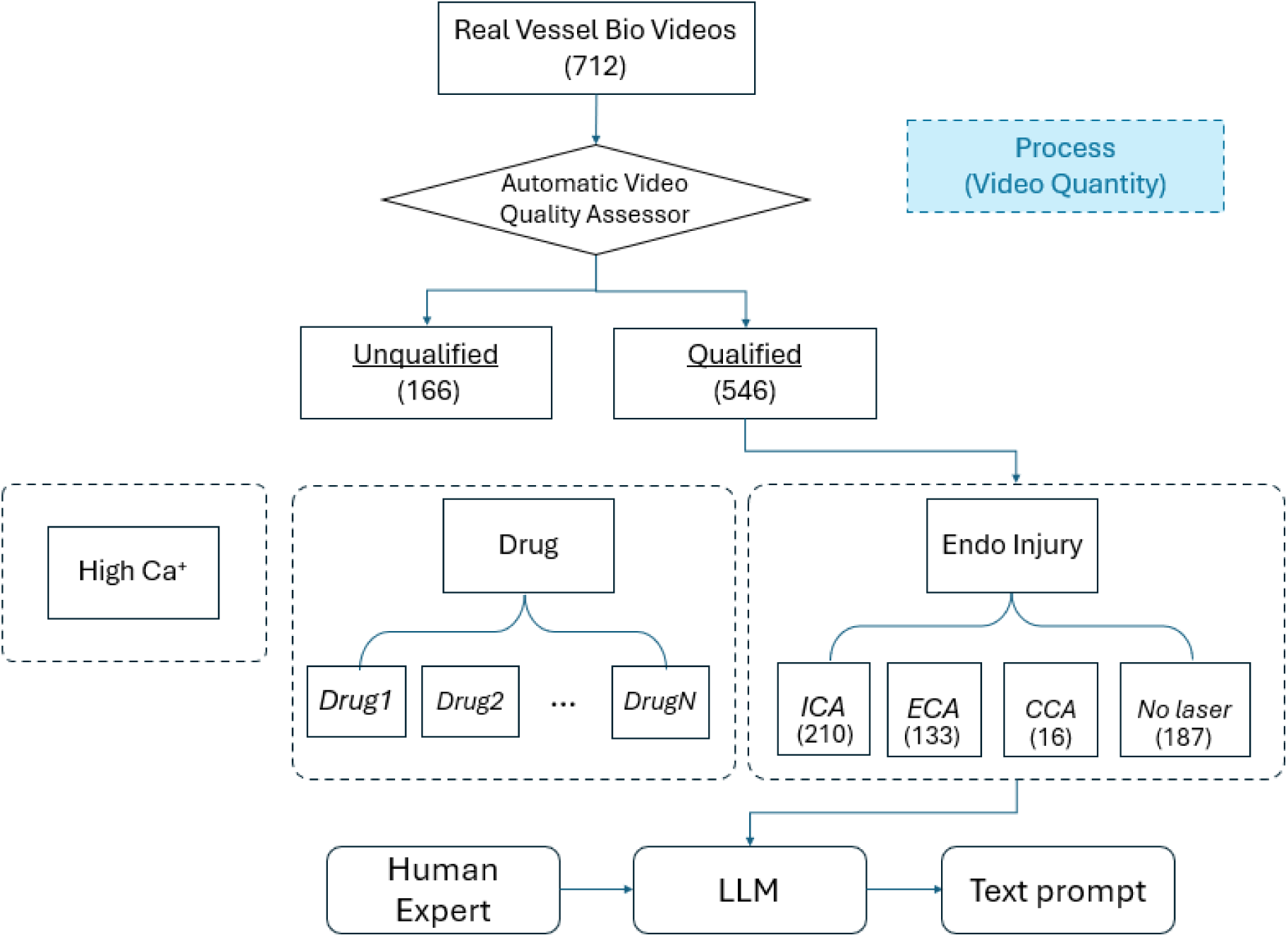
Automated video-quality check and prompt generation. Flowchart of dataset ingestion and preprocessing. A high-quality reference video with fluorescence-labeled vessels is used to calibrate thresholds. The pipeline automatically excludes clips with unexpected conditions or poor quality—e.g., out-of-focus/blurry frames, debris-laden flow, incomplete endothelial coverage, or channel artifacts—and removes them from the training set. For videos that pass quality check, a representative fluorescence frame and structured metadata (endothelial injury site, drug, calcium concentration) are provided to a large language model to synthesize standardized text prompts for training and inference. As displayed in the chart, we collected 712 biological videos with our “physical twin” in total and got 546 qualified videos after quality check.

**Figure S22.**
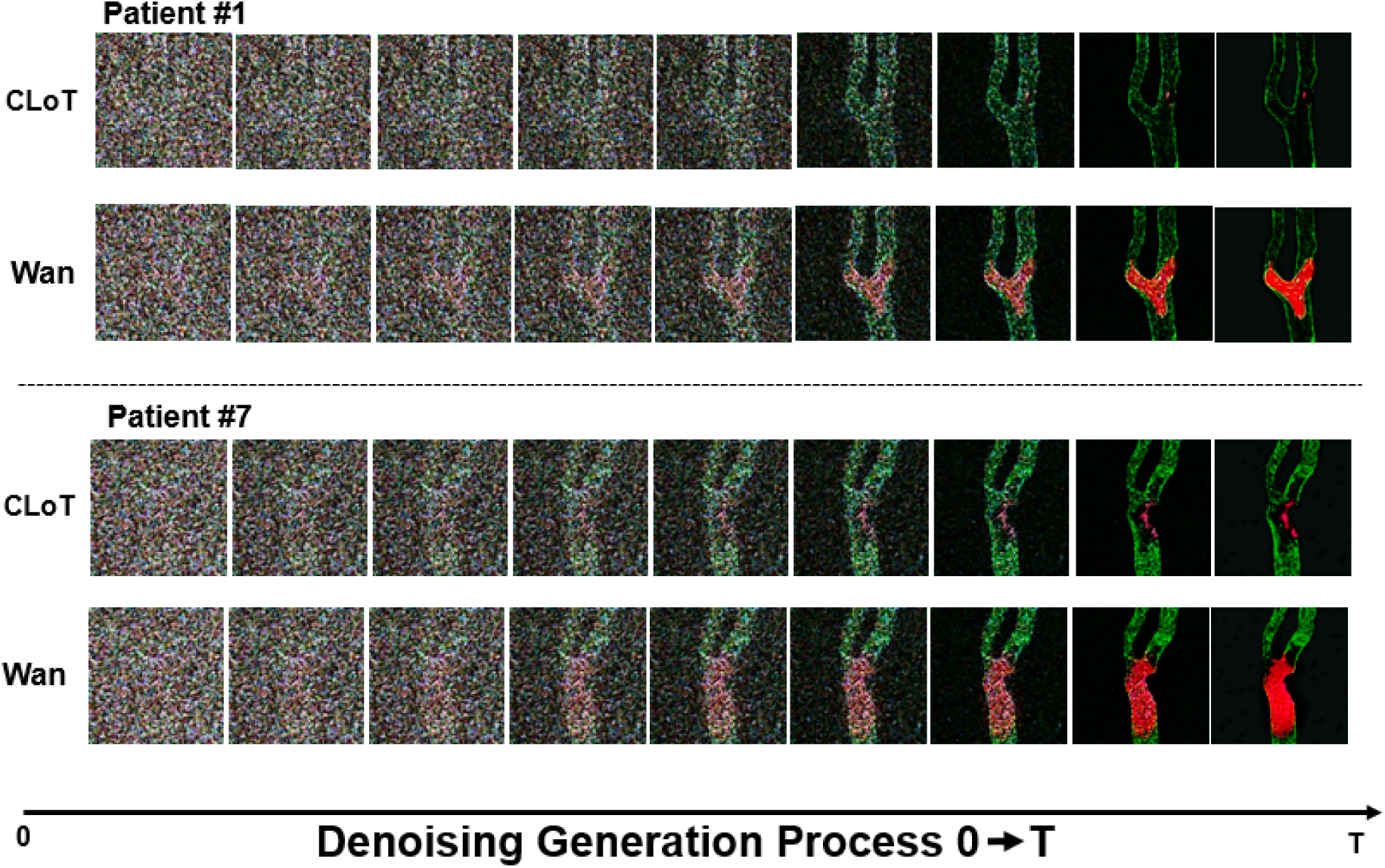
Denoising evolution of the middle frame for CLoT model compared with Wan model. Comparison of frame-by-frame denoising progression during the generative inference phase for an example test-set video (patient #1, ICA injury, rivaroxaban treatment and patient #7). Top panels show CLoT-generated frames at diffusion steps t = 0, 5, 10, 20, 30, 35, 40, 45, 49 (progressively less noisy); Bottom panels show Wan baseline generations at corresponding steps. CLoT exhibits faster convergence to realistic platelet morphology and spatial distribution, while Wan produces more diffuse, morphologically implausible structures (visible as overly smooth or fragmented aggregates lacking biological fidelity).

**Figure S23.**
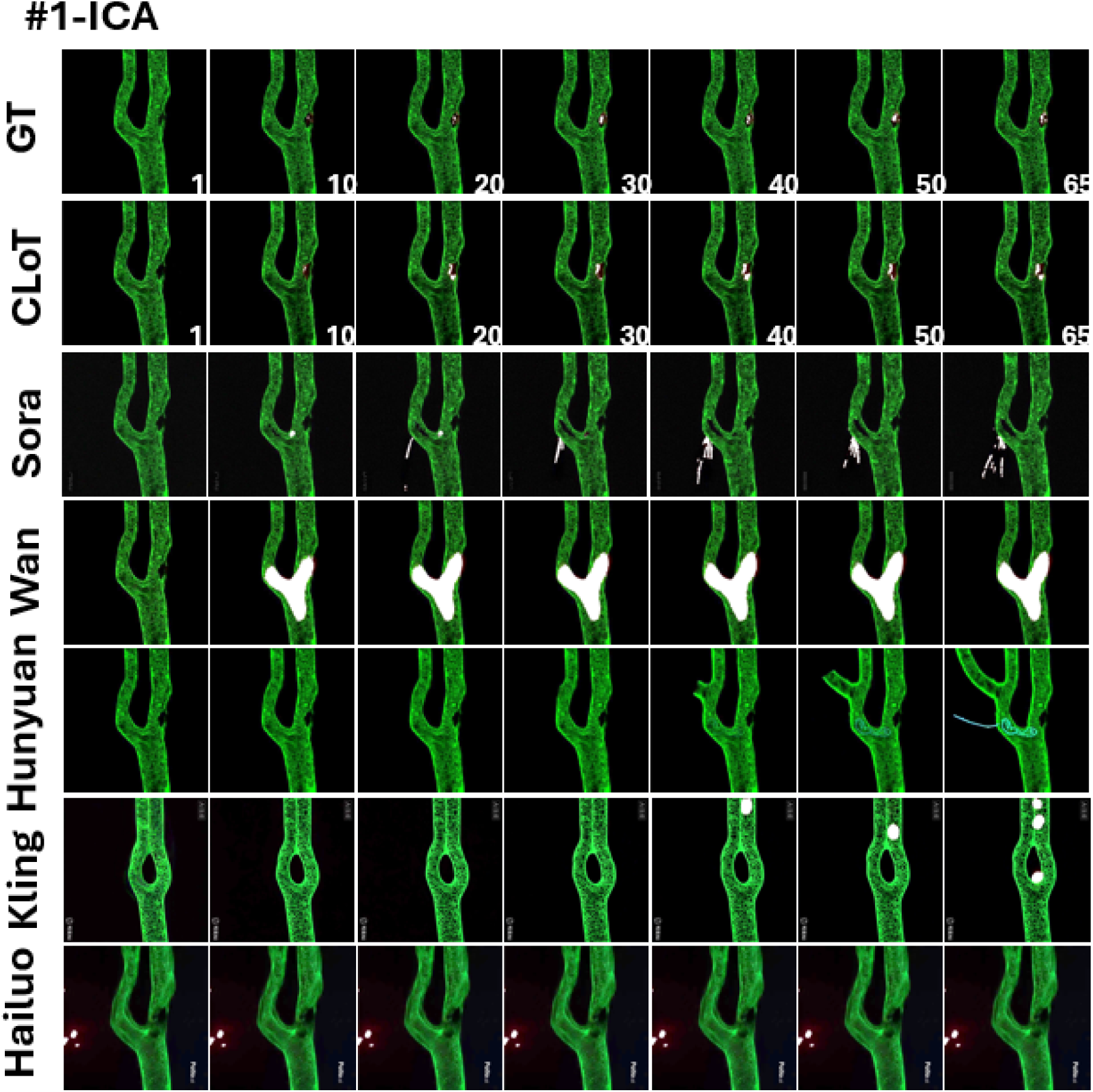
Representative frames for patient #1 from videos generated by CLoT (ours), Sora, Wan, Hunyuan, Kling, and Hailuo, compared to the “physical twin” ground-truth (GT). We display frames 1, 10, 20, 30, 40, 50, and the final frame (65). Using identical prompts and conditioning inputs, CLoT yields the closest correspondence to GT in vessel geometry and platelet-aggregation dynamics.

**Figure S24.**
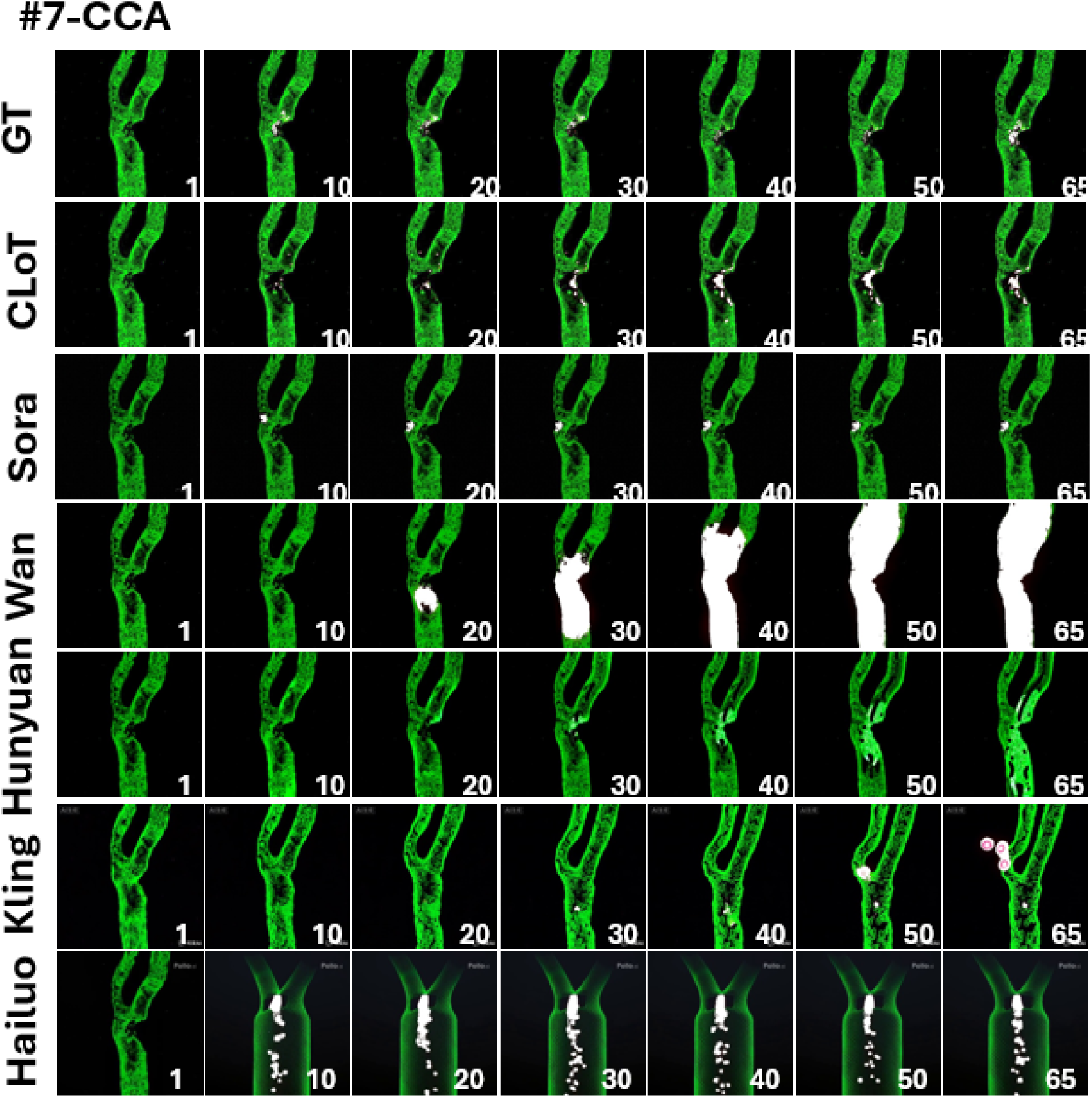
Representative frames for patient #7 from videos generated by CLoT (ours), Sora, Wan, Hunyuan, Kling, and Hailuo, compared to the “physical twin” ground-truth (GT). We display frames 1, 10, 20, 30, 40, 50, and the final frame (65). Using identical prompts and conditioning inputs, CLoT yields the closest correspondence to GT in vessel geometry and platelet-aggregation dynamics.

**Figure S25.**
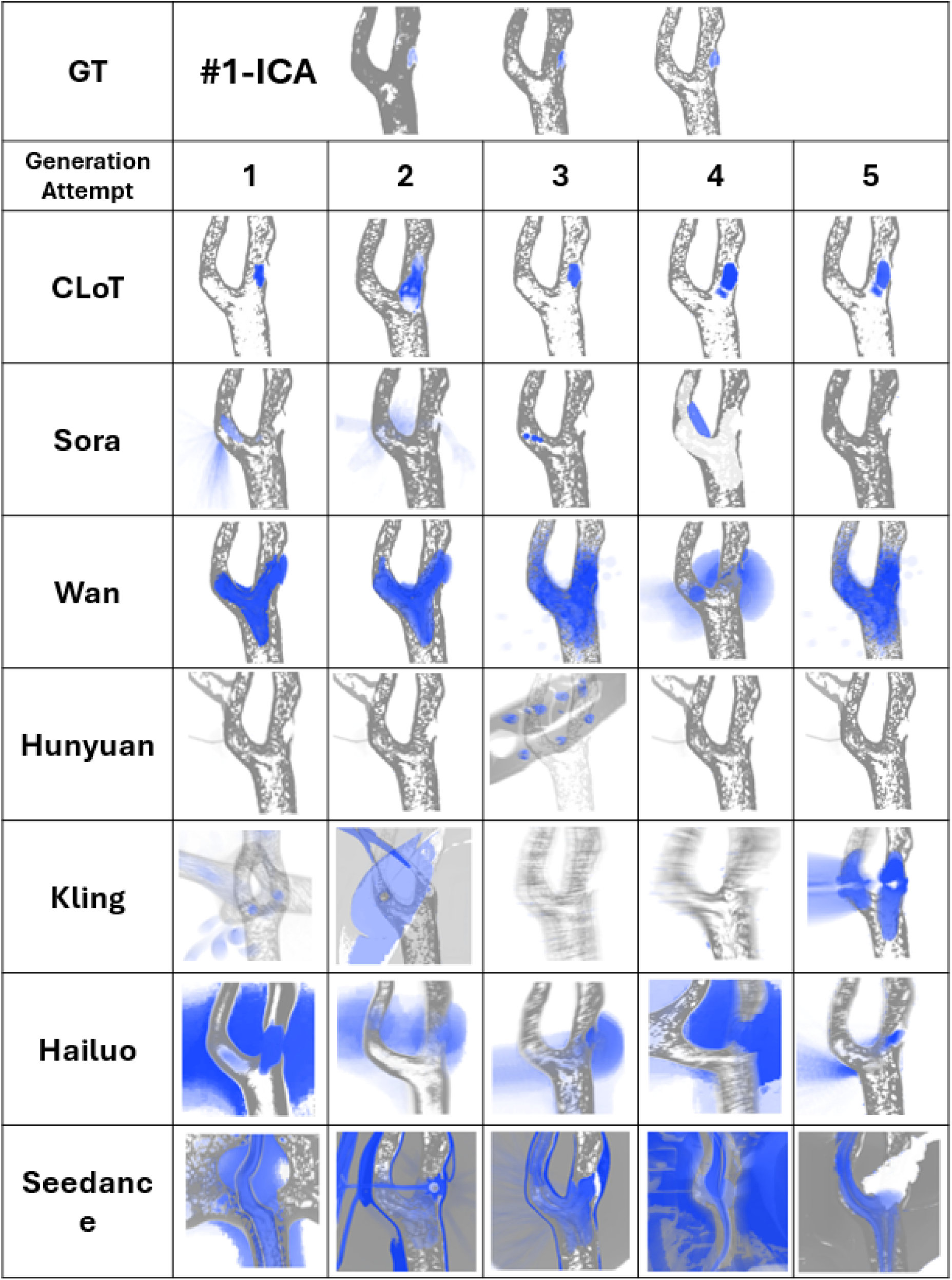
Comprehensive Replicate-Level Comparison of Platelet Activation Trajectories Across AI Models (Patient #1-ICA injury site). Temporal-trajectory (“long-exposure”) visualizations for all video replicates, Patient #1 ICA injury site. Each panel represents one 10-minute video’s cumulative platelet activation pattern (vessel structure: gray, platelets: blue; opacity reflects signal persistence). Top row: Ground truth (*n* = 3 replicates). Subsequent rows: AI model replicates (*n* = 5 generation attempts each): CLoT, Sora, Wan, Hunyuan, Kling, Hailuo, Seedance.

**Figure S26.**
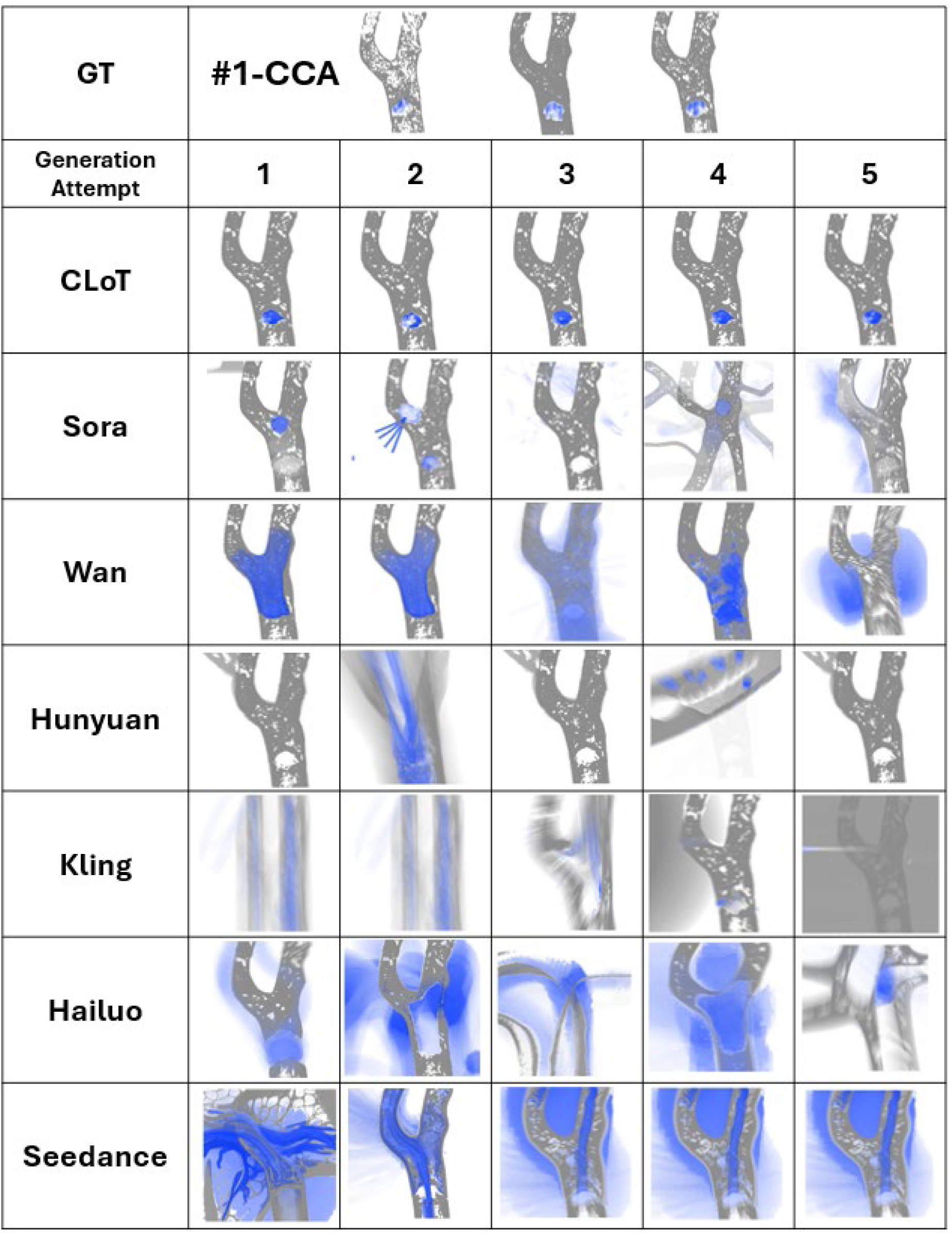
Comprehensive Replicate-Level Comparison of Platelet Activation Trajectories Across AI Models (Patient #1-CCA injury site). Temporal-trajectory (“long-exposure”) visualizations for all video replicates, Patient #1 CCA injury site. Each panel represents one 10-minute video’s cumulative platelet activation pattern (vessel structure: gray, platelets: blue; opacity reflects signal persistence). Top row: Ground truth (*n* = 3 replicates). Subsequent rows: AI model replicates (*n* = 5 generation attempts each): CLoT, Sora, Wan, Hunyuan, Kling, Hailuo, Seedance.

**Figure S27.**
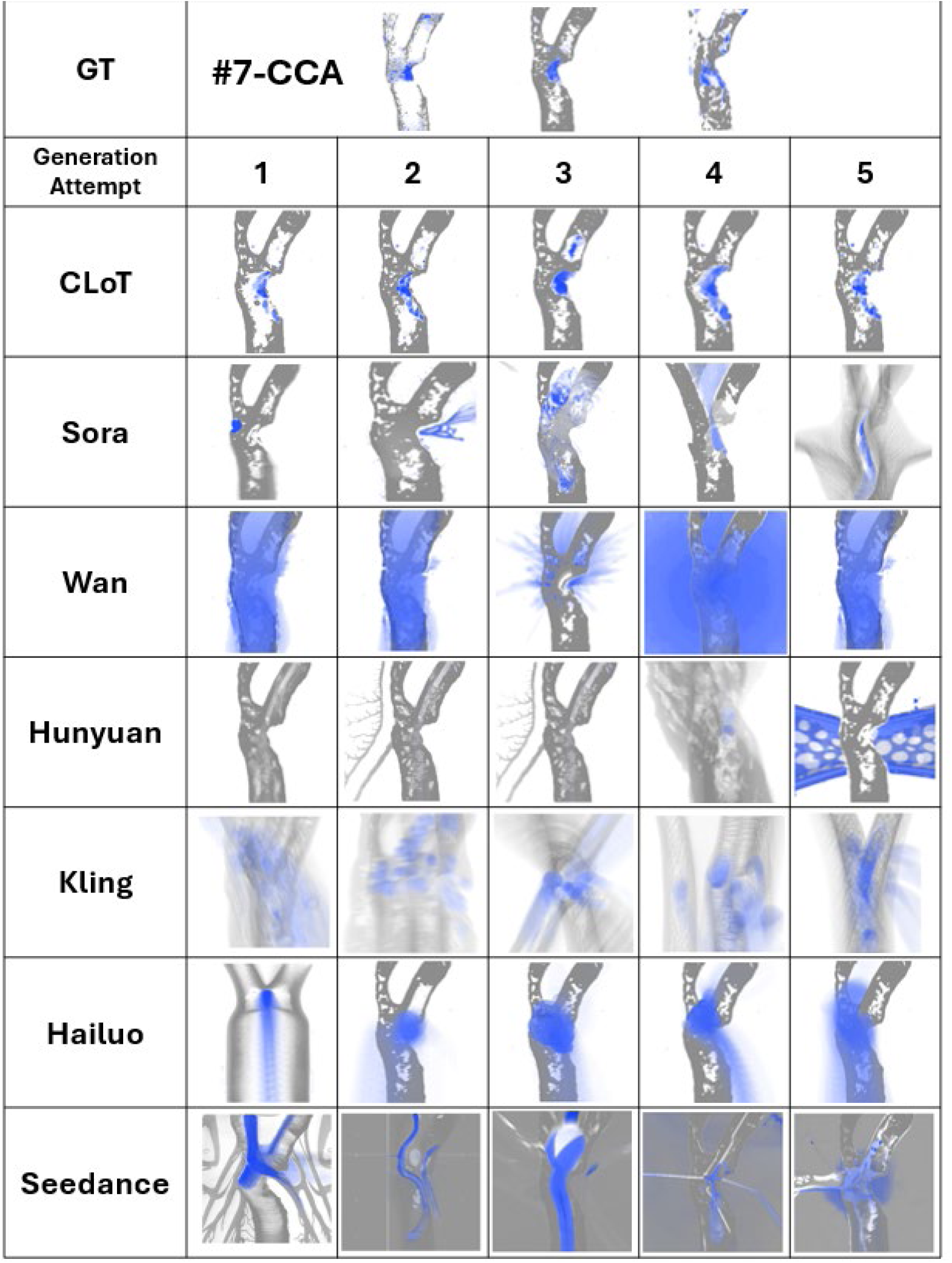
Comprehensive Replicate-Level Comparison of Platelet Activation Trajectories Across AI Models (Patient #7-CCA injury site). Temporal-trajectory (“long-exposure”) visualizations for all video replicates, Patient #7 CCA injury site. Each panel represents one 10-minute video’s cumulative platelet activation pattern (vessel structure: gray, platelets: blue; opacity reflects signal persistence). Top row: Ground truth (*n* = 3 replicates). Subsequent rows: AI model replicates (*n* = 5 generation attempts each): CLoT, Sora, Wan, Hunyuan, Kling, Hailuo, Seedance.

**Table S1.**
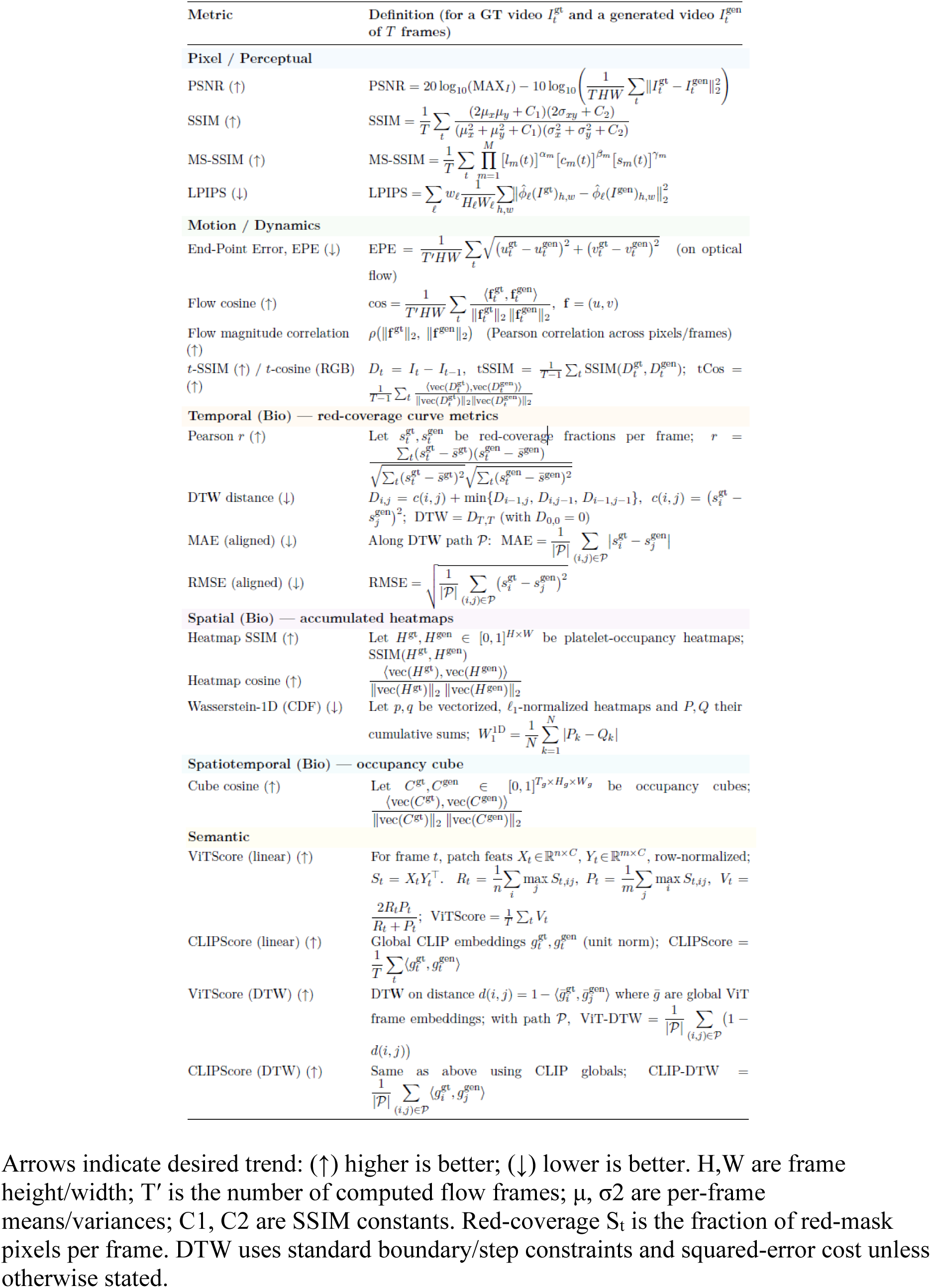
Metric definitions used for quantitative evaluation.

